# Nonlinear delay differential equations and their application to modeling biological network motifs

**DOI:** 10.1101/2020.08.02.233619

**Authors:** David S. Glass, Xiaofan Jin, Ingmar H. Riedel-Kruse

## Abstract

Biological regulatory systems, such as transcription factor or kinase networks, nervous systems and ecological webs, consist of complex dynamical interactions among many components. “Network motif” models focus on small sub-networks to provide quantitative insight into overall behavior. However, conventional network motif models often ignore time delays either inherent to biological processes or associated with multi-step interactions. Here we systematically examine explicit-delay versions of the most common network motifs via delay differential equations (DDEs), both analytically and numerically. We find many broadly applicable results, such as the reduction in number of parameters compared to canonical descriptions via ordinary differential equations (ODE), criteria for when delays may be ignored, a complete phase space for autoregulation, explicit dependence of feedforward loops on a difference of delays, a unified framework for Hill-function logic, and conditions for oscillations and chaos. We emphasize relevance to biological function throughout our analysis, summarize key points in non-mathematical form, and conclude that explicit-delay modeling simplifies the phenomenological understanding of many biological networks and may aid in discovering new functional motifs.

## Introduction

Biological regulation consists of complex networks of dynamical interactions [1–9]. For example, transcription factors regulate the production of proteins and other transcription factors [10–12], kinases and phosphatases regulate the behavior of enzymes and other kinases and phosphatases [1,4,5,8]. At a larger scale, cells regulate the growth of other cells [13,14], and populations of organisms interact with each other in ecosystems [15]. The term “network motif” has been coined to describe particularly important substructures in biological networks, such as negative feedback, feedforward regulation and cascades [2,11,16]. The function of these motifs often depends on emergent properties of many parameters, notably delays [12, 17–21], but in fact there has been no thorough treatment of network motifs with these delays included explicitly.

Here we provide a theoretical and practical basis for analyzing network motifs with explicit delays, and we demonstrate the utility of this approach in a variety of biological contexts. Previous work showed that behavior can change dramatically when explicit delay terms are incorporated in models of biological systems, as evidenced by the appearance of oscillations, for example [18,22,22–26]. These delay effects may occur at multiple spatial and temporal scales. For example, at the intracellular scale, natural [27–29] and synthetic [30,31] genetic oscillators have been shown to depend on delays, as have coupled intercellular oscillators [24,26,32] and physiological conditions such as respiration [33]. At the multicellular scale, many delayed regulation mechanisms have been examined in developmental pathways [2,12], where delays can be significant and crucial to proper formation of large-scale structure such as the vertebrate spine [22,26,34–36]. We have also shown previously that the delays expected in Delta-Notch-based lateral inhibition patterning may be important for ensuring patterning fidelity [32]. Delays in nervous sstems arise in various axon conduction and signal integration steps, and influence the presence or absence of synchronicity [37,38]. On an ecological scale, delays in interactions such as predator-prey relationships induce population booms and busts [13]. It is thus clear that delays play a significant role in regulatory networks in biological systems.

On a conceptual level, regulatory networks are often depicted as directed graphs, for example with biological molecules represented as nodes, while directional edges indicate activation and repression using regular and blunted arrows, respectively (Fig 1). Much information is left out in such cartoons. Notably, these diagrams are generally not drawn with time scales associated with particular arrows. This reflects the fact that the amount of time for a particular regulatory step may depend on multiple parameters, such as removal rates and activation thresholds [2]. Furthermore, a single arrow can often schematically represent many individual, biochemical steps [27, 39] as a way to simplify a model of the actual, more complex network.

**Fig 1.**
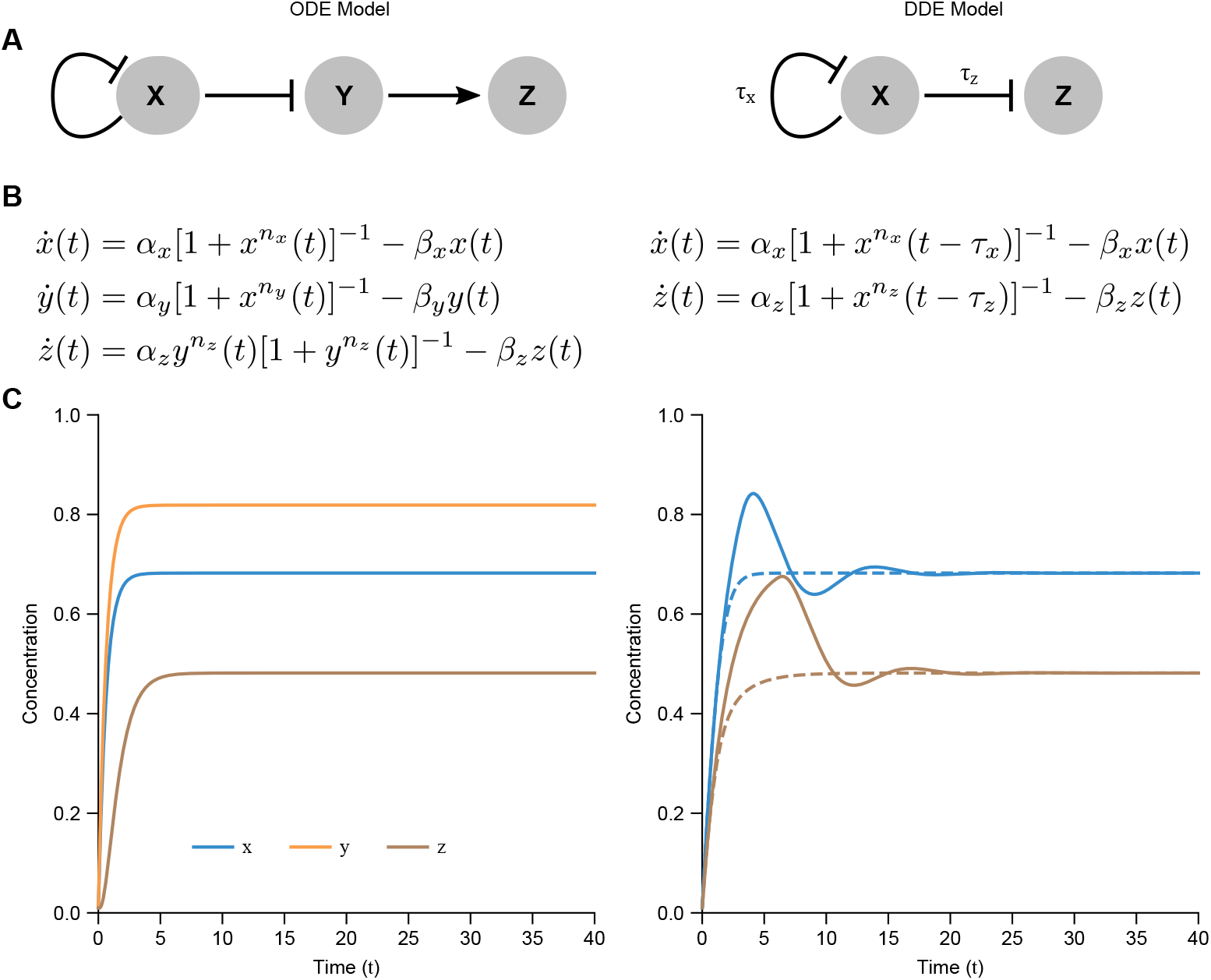
Explicit inclusion of time delays in mathematical models of network motifs can reproduce non-delay models using fewer variables and parameters, but can also lead to more complex behavior. (A) An example genetic regulatory network including genes *X*, *Y*, and *Z*, regulated by one another either positively (activation, regular arrow) or negatively (repression, blunt arrow). Model on left incorporates no explicit delay terms, whereas model on right incorporates explicit delay terms *τ*_1,2_. (B) Ordinary differential equations (ODEs) and delay differential equations (DDEs) corresponding to (A) with regulation strengths *α_i_*, removal rates *β_i_* and cooperativities *n_i_*. (C) Numerical simulations of equations in (B) for one set of initial conditions using parameters *α_x_* = 1, *α_y_* = 1.2, *α_z_* = 1.2, *β_x_* = *β_y_* = *β_z_* = 1, *n_x_* = *n_y_* = *n_z_* = 2 for the ODE simulation, and *α_x_* = 0.5, *α_z_* = 0.3529, *β_x_* = *β_z_* = 0.5, *n_x_* = *n_z_* = 2 for the DDE simulations (dashed curves: *τ_x_* = 0.8, *τ_z_* = 0.1; solid curves: *τ_x_* =3, *τ_z_* = 4). Note that delays can cause more complex dynamics (e.g., transient oscillations) compared to models in which effects are instantaneous, and where both models give the same long-term steady state behavior. Left: instantaneous effects modeled using ODEs, Right: delayed effects modeled using DDEs.

Mathematically, these regulatory cartoons are often modeled using ordinary differential equations (ODEs), in which all information is assumed to pass from one variable to another instantaneously (with relative timing often determined by a rate constant). This may be a reasonable assumption if any intermediate steps are fast, but that is not always the case. For multicellular systems, for example, the time it takes to convert an incoming signal from one cell first into gene expression, then into an external signal for another cell to measure, and finally into gene expression in that downstream cell is not negligible compared to differentiation times [32,40]. Thus ODE models of simplified networks may fail to capture real delays and their effects by oversimplification or require additional variables and parameters to predict more complex phenomena [26, 32, 40].

An intriguing solution to making the time scales and delays explicit is to use delay differential equations (DDEs). As opposed to ODEs, DDEs have derivatives which depend explicitly on the value of variables at times in the past. For example,

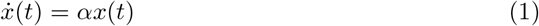

is an ODE, and

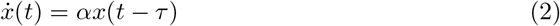

is a corresponding DDE, where *x*(*t* – *τ*) represents the value of *x* at a time *τ* units in the past, making the effect of *x* on the current rate of change of *x* delayed by a time *τ*. The time scales and delays are thus explicit (Fig 1). Multiple steps (“cascades”) within a network can then be more rigorously simplified into a single step with delay (see Fig 3). This makes interpretation of the phenomenology simpler and reduces the number of equations and parameters in the model [27, 32].

DDE models also expand the available dynamics in a model with this reduced set of equations and parameters. With ODEs, a 1-variable system such as Eq 1 can only lead to simple exponential growth or decay with no overshooting or oscillations, as guaranteed by the Poincare-Bendixson theorem [41,42]. A corresponding DDE with the same equation, such as Eq 2, can lead to stable, unstable, bistable, oscillatory, or chaotic behavior depending on parameters [20, 33, 43–47]. It may thus be possible to use simplified network models to examine comparatively larger networks without losing key information about the dynamics of an original, more detailed system.

A key challenge in using DDEs to achieve modeling simplicity with diverse available dynamics is their mathematical complexity relative to ODEs [48]. For example, a 2-component model consisting of variables *X* and *Y* can be represented as a 2-dimensional ODE system, as the system can be fully described at any given time by the two instantaneous values of *X* and *Y*. However, the analogous DDE model is in fact infinite-dimensional, as to describe such a system requires not just the two instantaneous values of *X* and *Y*, but their full historical values up to a delay *τ* in the past. Accordingly, an entire time interval −*τ* ≤ *t* ≤ 0 must be specified in the initial conditions of a DDE system for a unique solution [49]. A first-order, single-variable, linear ODE with constant coefficients such as Eq 1 has a single exponential solution, while the corresponding DDE, Eq 2, has exponential solutions corresponding to an infinite number of eigenvalues derived from a transcendental characteristic equation. It is precisely this added mathematical complexity that expands the available dynamics while using the same number of equations.

Despite the challenges, much progress has been made into analytical understanding the evolution and stability of delay differential equations [45, 50–57] as well as systems thereof [42, 58]. When analytic methods become intractable, numerical methods are available to simulate the behavior of these equations [59,60]. We see an opportunity to use DDEs to recapitulate dynamics found in ODE solutions of network motif behavior with fewer genes and thus fewer modeling parameters and equations (see Fig 1), a type of “modeling simplicity.”

In this work, we thoroughly examine the most common network motifs [2,11] with explicit delays and present an approachable, step-by-step view of the mathematical analysis in order to make such delay equations easy to use for biologists and others. We derive the complete phase space for autoregulation with delay, demonstrating explicitly the regimes in which delays are and are not important. We show that the essential behaviors of all eight [61] feed-forward motifs can be recapitulated in a single feed-forward motif with delays. We next discuss more complex situations that include two-component feedback loops, multiple delays with feedback, and arbitrary networks of interactions. A reference summary is provided in Table 3. Finally, we discuss how these results can be applied to understanding fundamental design principles of various natural biological systems.

## Results

Our results are divided up into 7 sections corresponding to 7 different regulatory networks of increasing complexity. We start with simple regulation with delay (Motif 0) and cascades (Motif I), including their demonstration of how the DDE models relates to multi-step ODE models. We then cover autoregulation (Motif II), logic (Motif III), feedforward loops (Motif IV), multi-component feedback (Motif V), and multiple feedback (Motif VI). Each section, in addition to analyzing a network of biological importance, includes new methods of DDE analysis and their implications for biological network motifs. Multiple new results are also useful for the more conventional ODE network analysis.. Each section then concludes with the “key results” for the relevant motif in non-mathematical terms, and an overall reference table (Table 3) is included in the discussion.

### Motif 0: Direct Hill regulation

As a prerequisite for understanding more complex network motifs, we first describe how we use DDEs to model simple regulation of a gene *Y* by a gene *X* as the most basic motif, and then provide a simple yet unified mathematical framework for both activation and inhibition with delay.

#### Activation and inhibition can both be modeled with a single unified function

As a concrete example of biological regulation [62], we first consider a transcription factor *x* regulating the production of a protein *y* (see Fig 2). If *x* activates *y*, the production rate of *y* increases with increasing *x*, generally saturating at a maximum rate *α*. Often there is an additional cooperativity term *n* which determines how steeply *y* increases with *x* in the vicinity of the half-maximal *x* input value *k*. In this framework, *n* = 0 is constitutive production, *n* = 1 is a Michaelis-Menten regulation, 1 < *n* < 5 is a typical biological range of cooperativity, and *n* → ∞ is a step-function regulation. If *x* represses *y*, the same conditions hold except that the production rate of *y* then decreases with increasing *x*. A standard quantitative model for this behavior is called the Hill function, and serves as a good approximation for many regulatory phenotypes in biology, including transcription and translation rates, phosphorylation, enzymatic activity, neuronal firing, and so forth [63,64]. In general there may also be a leakage rate that yields constant *y* production in the absence of *x*. The concentration of *y* also generally removed at a rate *β* proportional to its concentration. This removal term can represent many biophysical processes, such as degradation, dilution, compartmentalization, or sequestration [65,66]; for simplicity we mainly use the term “degradation.”

**Fig 2.**
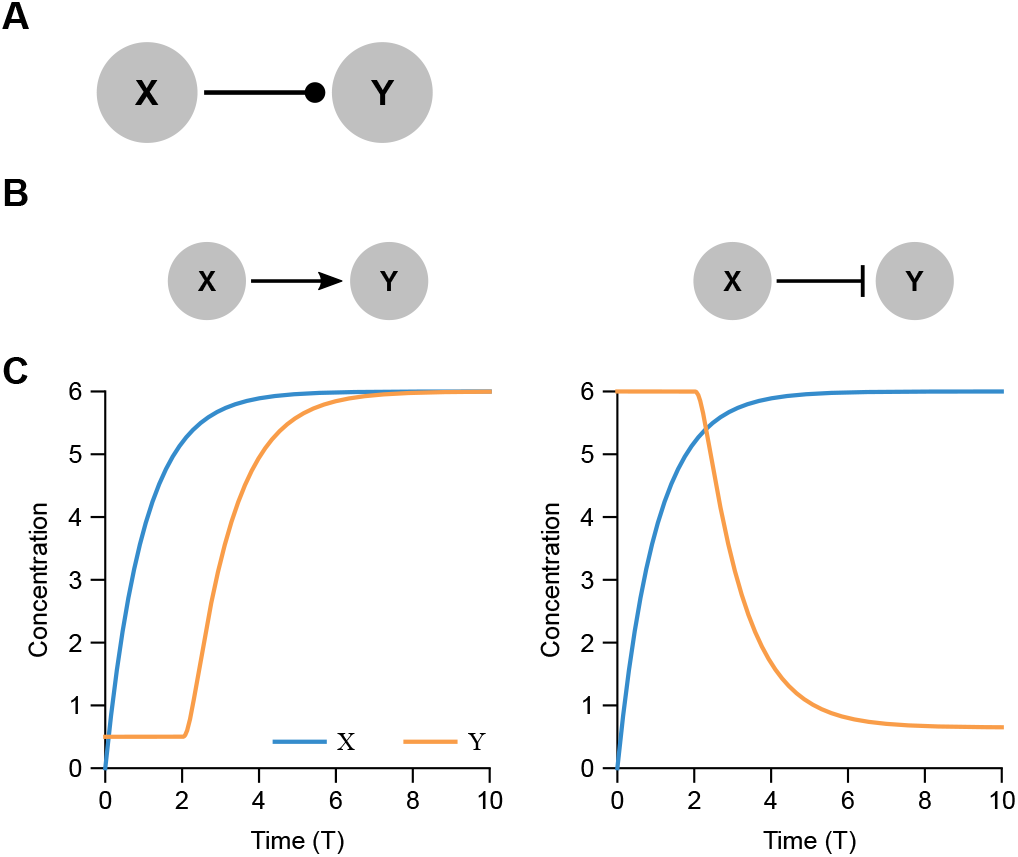
Direct regulation by activators and inhibitors. (A) The simplest regulation network consists of a single input *X* directly regulating a single output *Y*. We use a dotted arrow to represent either activation or repression, with an implied explicit delay. (B) Direct activation (left) is represented by a pointed arrow and direct repression (right) by a blunt arrow. (C) Time response (Eq 7) of activated (left) and repressed (right) *Y* following a rise in *X* that exponentially approaches a new steady state (*X* = 6) from zero (governed by 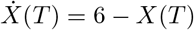). Note that the finite value of *η_X_* leads to an effective leakage slightly greater than *ϵ* for the repressor case. For the activator, *n* = −2 and 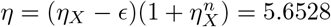 to match the *X* steady state. For the repressor, *n* = 2 and *η* = *η_X_* − *ϵ* = 5.5 to match the *X* steady state. In both cases, *γ* = 2, *ϵ* = 0.5.

Together these can be written as:

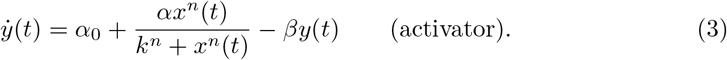

If *x* instead “represses” *y* instead of activating it, the form is similar:

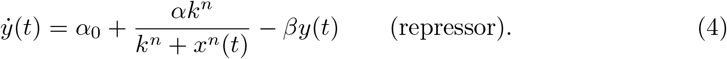

For biologically meaningful results, all variables and parameters in these equations should be real and non-negative. Note, however, that the activator case is equivalent to the repressor case with *n* < 0. In this paper we will therefore allow *n* to be negative, which is simply a notational modification used in order to combine the two cases. With this notation, the effective cooperativity is |*n*|. Thus, we have:

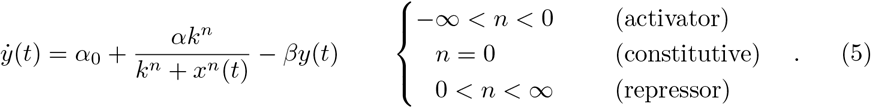

which provides a unified, single-function description for both activators and inhibitors (and constitutive expression), providing a powerful mechanism to analyze both cases simultaneously as we will do throughout this work. The activator case technically fails when *x* is identically zero, since that would imply division by zero, but the limit as *x* goes to zero causes the regulation term to be zero, which is the same result as assumed by our notation. An initial value of exactly zero for *x* can thus lead to a divide-by-zero error in simulations, but is easily avoided. The consitutive case is degenerate, in that *n* = 0, *α* ≠ 0 is equivalent to *n* ≠ 0, *α* = 0 with *α*_0_ → *α*_0_ + *α*/2.

The negative value of *n* allowed here should not be confused with the term “negative cooperativity” used in the binding kinetics literature [67], which in our notation would refer strictly to a negative value of *n* with magnitude less than one (i.e., −1 < *n* < 0). The Hill function has an inflection point for all |*n*| > 1 (see Appendix S2), which allows there to be 3 fixed points for *n* < 0 when feedback is introduced below (see Fig. S2).

#### Delays may be present in regulation, but are not modeled in removal

For an explicit delay *τ* in regulation, we would replace *x*(*t*) with *x*(*t* − *τ*) in the regulation (Hill) term, but not in the removal term, which is directly dependent on *y* itself, and thus not expected to have any delay. This form of regulation is, as noted, quite general, but for concreteness we will generally refer to quantities like *x* (or *y*) as the concentrations of some transcription factor or protein *x*. Thus we arrive at an explicit-delay model of activating or repressive biological regulation as follows:

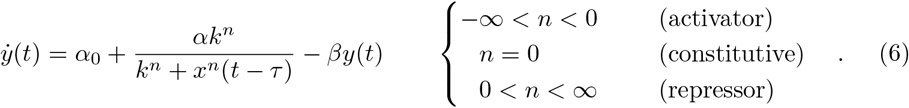

#### Nondimensionalizing yields 4 key parameters for any regulation

In analyzing various network motifs, we will generally work with a non-dimensionalized version of this equation, formed by dividing all concentrations by the half-maximal input concentration *k* and dividing times by the degradation time 1/*β*, which has the effect of measuring concentrations in units of *k* and times in units of 1/*β*. This yields a simplified equation as follows:

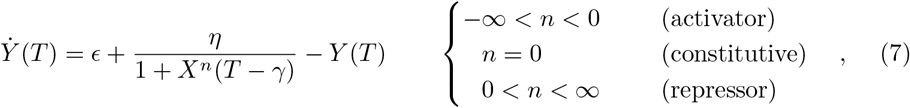

with dimensionless variables *X* = *x/k*, *Y* = *y/k*, *T* = *tβ*, and dimensionless parameters *ϵ* = *α*_0_/*kβ*, *η* = *α*/*kβ*, *γ* = *τβ*, and *n*. We thus reduce the number of parameters from 6 to 4, and as discussed below, primarily *η* and *γ* are important. In this form, *η* is a normalized “regulation strength” and *γ* is a normalized delay. Again, the consitutive case is degenerate, in that *n* = 0, *η* ≠ 0 is equivalent to *n* ≠ 0, *η* = 0 with *ϵ* → *ϵ* + *η*/2. The dynamics of direct activation and repression via Eq 7 are demonstrated in Fig 2C.

The dimensionless nature of the remaining parameters additionally implies that these parameters are “big” or “small” depending on whether they are simply large or small compared to 1, respectively. For example, *γ* = *τβ* ~ 1 implies that the delay (*τ*) and degradation (1/*β*) times are approximately equal. In subsequent sections we therefore almost always explore parameter values linearly between 0.1 and 10.

#### Key results for direct regulation

1. Time scales are specified explicitly as a normalized ratio of delay to degradation times.
2. We use standard Hill equations for regulation, with the regulation term delayed in time but no delay in the degradation term.
3. We simplify and unify the use of both activators and repressors in a single equation by allowing negative Hill coefficients to represent activators. This approach is useful for both ODEs and DDEs.
4. Nondimensionalizing regulatory functions reduces the number of parameters by measuring variables and parameters in the relevant scales.
5. Nondimensionalized paramters can be considered large or small by comparing their values to depending on whether they are greater or less than 1, respectively.

### Motif I: Cascade (sequential regulation)

A common network motif in many biological networks is the cascade (see Fig 3), a series of regulatory steps (i.e., *x* regulates *y*, which regulates *z*, etc.) [2,11,68]. Since each regulator must reach the corresponding half-maximal input value *k* before significantly affecting the next item in the cascade, each step adds an effective delay, as we discussed qualitatively above. We show here that in fact, mathematically, this cascade can be approximated well by a single regulation (Hill function plus leakage) with the addition of explicit delay (Fig 3). Furthermore, a cascade of regulation steps each with an explicit delay is well approximated by a single regulation step with an explicit delay equal to the sum of delays in each step (Fig 3). These together form the mathematical basis validating the use of DDEs with single delay terms that consolidate multiple phenomenological regulation steps.

**Fig 3.**
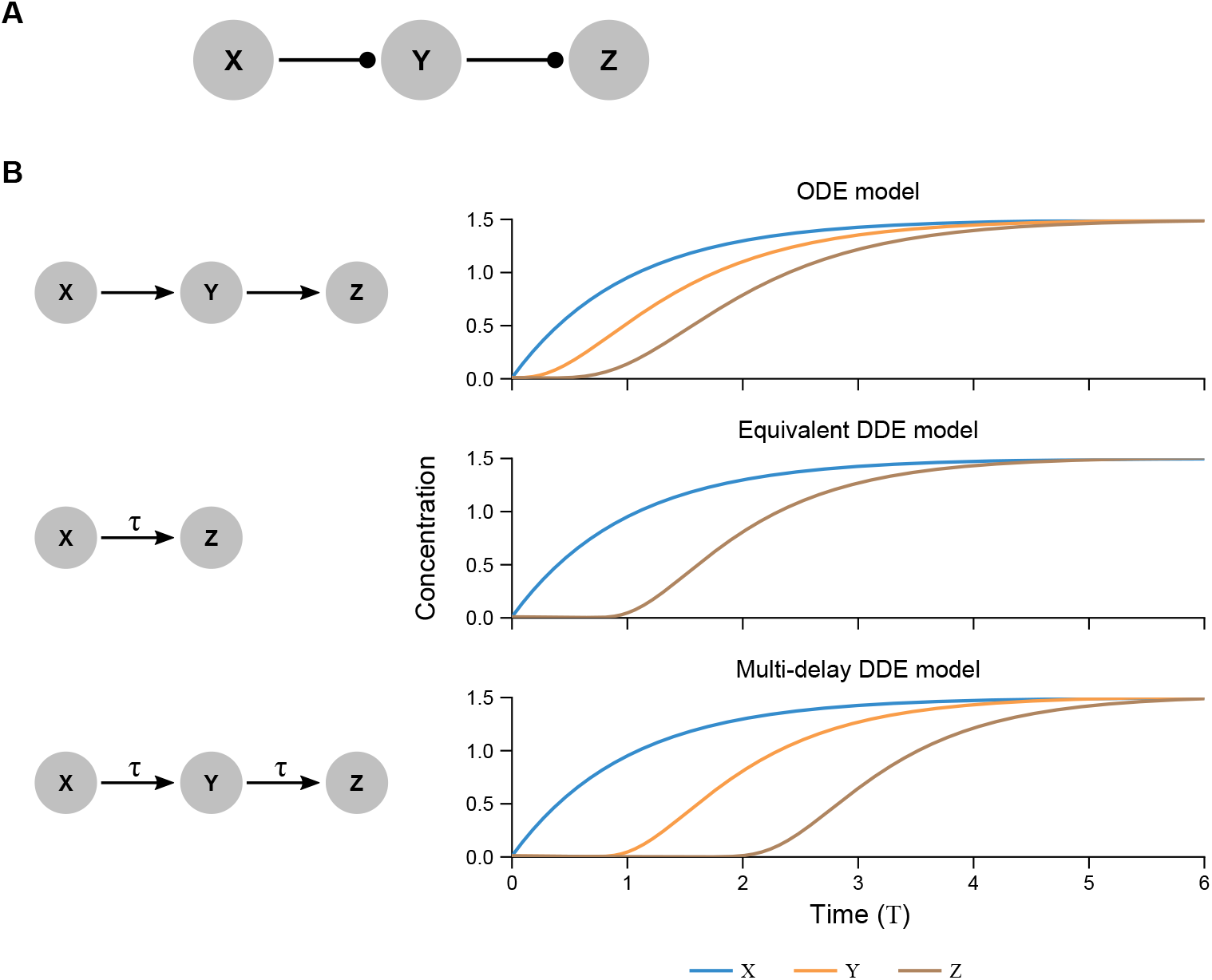
Cascades with zero, one, and multiple delays. (A) A cascade is a linear sequence of regulation steps, here *X* regulating *Y* regulating *Z*. (B) Standard models of cascades use ODEs (top), in which each step leads to a characteristic delay in the products *Y* and *Z*, which increases with each step based on the half-maximal inputs and degradation rates for each step. In an equivalent DDE model (middle), similar behavior is accomplished by replacing the explicit cascade of implicit delays with a single-step regulation including an explicit delay. A cascade in which each step contains an explicit delay (bottom) behaves analogously to delayed direct regulation (as in middle), with the final step delayed by the sum of delays in each step. *η_X_* = 0.667, *η_Y_* = *η_Z_* = 3.25, *β_X_* = 0.667, *β_Y_* = *β_Z_* = 1, *n_X_* = *n_Y_* = −2. For the bottom graph in (B), each step is governed by Eq 9 with the same parameters.

#### Delayed direct regulation approximates cascades of non-delayed regulation

For a non-delayed cascade motif in which *X* regulates *Y*, which in turn regulates *Z* (Fig 3), one can write down a nondimensional set of governing equations as follows:

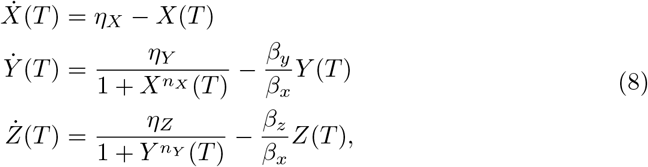

in which *β_x_, β_y_*, and *β_z_* are the dimensional degradation rates of *X, Y*, and *Z*, respectively (see Appendix S1).

We can show that an approximate single-step delayed regulation exists corresponding to this cascade by considering the following. If the system is initially at steady state *X* = *η_X_* and then *X* is perturbed, then *Y* follows with a timescale *β_y_/β_x_*. Meanwhile, *Z* continues to remain at its original steady state value until *Y* changes appreciably, which happens after an amount of time that it takes *Y* to change (*T* ≈ *β_y_/β_x_*). Thus, at early times (*T* ≪ *β_y_/β_x_*), *Y* changes much more rapidly than *Z*, and so from the perspective of *Z* we can set 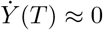. Then this pseudo-steady state value of *Y* can be inserted into the Hill term of 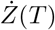, but with a delay *γ* = *β_x_/β_y_* reflecting the time that *Y* takes to change.

Finally, by matching the values of the composite function at *X* = 0, *X* = 1, and *X* → ∞ as well its the slope at *X* = 1 compared to a single Hill function with leakage, we can approximate the cascade as a single-step regulation of *Z* by *X*:

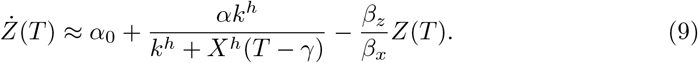

where the combined regulation parameters are (see Appendix S1)

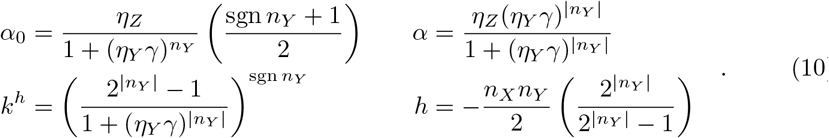

The *k* can be removed by renormalization of *X* and the degradation rates removed by renormalization of *T*. Note that because of the normalization *η_X_* = *α_x_/k_x_β_x_* (see Appendix S1), *γ* is inversely proportional to *η_X_*.

This equivalence of parameters shows several results of biological importance. First, the overall Hill coefficient *h* ≈ −*n_X_n_Y_*/2 is proportional to the negative product of the two individual Hill coefficients. Based on the signs of *h, n_X_*, and *n_Y_*, this means that two repressors or two activators act overall as an activator, whereas one activator and one repressor overall act as a repressor, as intuitively expected. Second, it also means that the cooperativity increases with the length of the cascade if cooperativities are above 1, as has been found experimentally [68]. Third, the leakage is zero for *n_Y_* < 0 and negligible (i.e., *α*_0_ ≪ *α*) for *n_Y_* > 0 if *η_Y_ γ* ≫ 1. This means that the maximum possible amount of *Y* produced during the effective delay time must be sufficient to repress *Z*. We have generally ignored this leakage term in the main text, for a more detailed discussion on leakage see Appendix S3 and Fig. S3.

#### The total delay in a cascade equals the sum of individual delays

For cascades with delays at each step, the same pseudo-steady-state analysis implies that a corresponding single-step delayed regulation can be analyzed in which the single delay is the sum of the multiple delays. In this case the effective delay from *X* to *Z* would be *β_x_/β_y_* + *γ_y_* + *γ_z_*, where *γ_y_* is the intrinsic delay between *X* and *Y* and *γ_z_* is the intrinsic delay already present between *Y* and *Z*.

Fig 3 shows simulations for a cascade of activators, perturbed from zero by a step function regulating the upstream *X*. In the ODE model, each step has a timescale of activation based on the cascade of degradation rates. The equivalent DDE model shows a single-step regulation from *X* to *Z* where the rise of *Z* is delayed to the same time scale as *Z* in the ODE model. The multi-delay cascade shows that additional delayed steps appear approximately identical to earlier steps in the cascade but shifted in time.

#### Key results for cascades

1. Multiple steps can be simplified to a single step with explicit delay in the regulation term.
2. Composed Hill functions are approximately equivalent to a single Hill function with leakage.
3. Shortening cascade leads to leakage term (even if there was none for the individual steps), but the leakage is small if the final-step regulation strength is smaller than the degradation rate of the initial step (*α_y_*/2*k_y_β_y_* ≫ 1).
4. The total delay in a cascade is approximately equal to the sum of all single delays and all degradation times.
5. The cascade overall is repressive when the number of individual repressing steps is odd; otherwise, the cascade is activating.

### Motif II: Autoregulation

Autoregulation describes the situation in which a single biological species (e.g., a transcription factor) regulates its own production (Figs 4A-C). It is one of the most common network motifs in biological regulatory networks [2, 69, 70]. In this section we analyze autoregulation with delay, deriving many predictions for its behavior, including a complete phase space based on all parameters.

**Fig 4.**
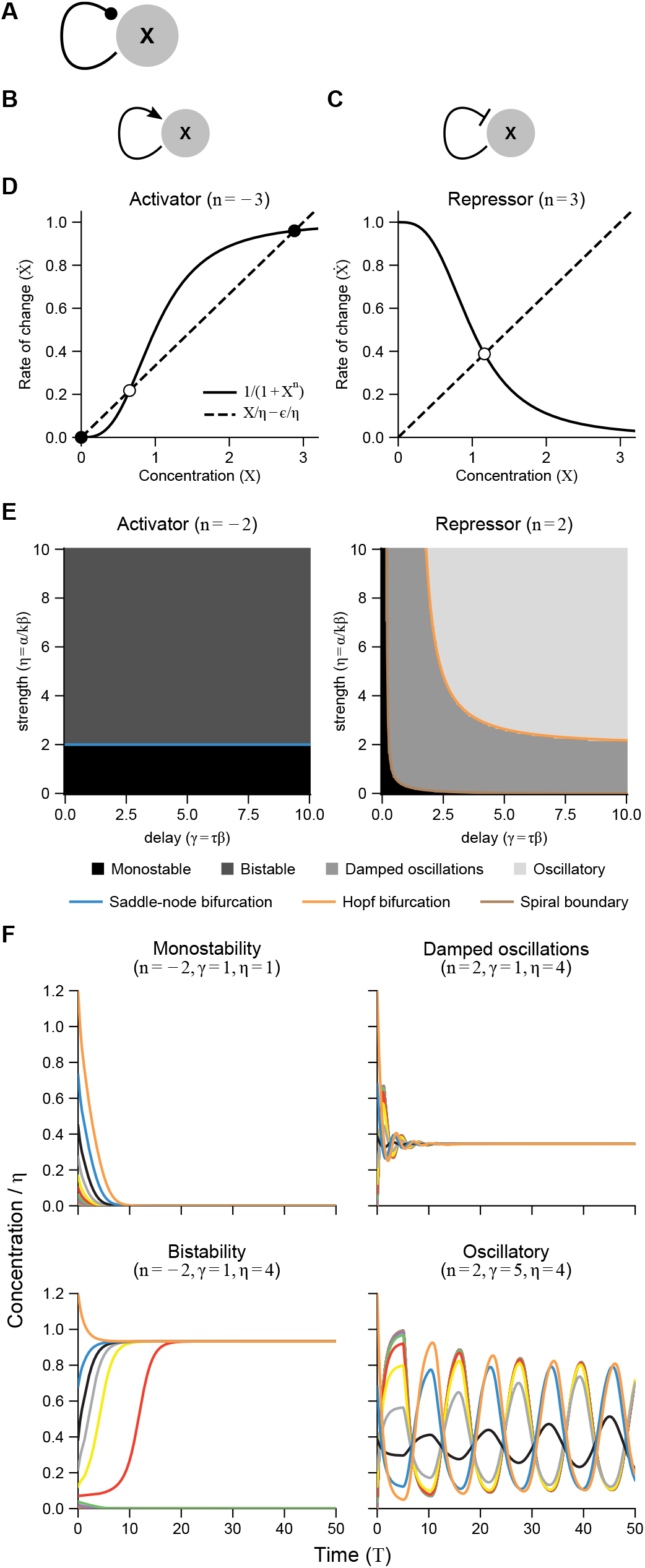
The complete phase diagram for the autoregulation network motif has analytically derivable parameter regions for bistability, monostablility, monostablility with damped oscillations, and oscillations. (A) The autoregulation network motif, with a dotted arrow indicating either (B) self-activation or (C) self-repression. The two cases are given by *n* < 0 (activation) and *n* > 0 (repression). (D) Stable (black circles) and unstable (white circles) fixed points for activator and repressor cases are given by the intersection of the regulation (solid) and degradation+leakage (dashed) lines. Note that activators can have 1, 2, or 3 fixed points, whereas repressors always just have one. (E) Parameter space showing all possible behaviors for autoregulation with delay. Shading shows simulation results (with an interval of 0.1 for both *γ* and *η* axes) and curves show the analytically derived bifurcation boundaries. See Fig. S1 for cases −3 ≤ *n* ≤ 3. (F) Representative simulation curves for the four qualitatively different behaviors, with different colors representing different initial conditions (see Materials and Methods). *ϵ* = 0.

#### The complete phase space for autoregulation demonstrates quantitative and qualitative importance of delays

Based on Eq 5, the governing equation for such a system with delayed regulation is given by setting *Y* = *X* (output equals input) in Eq 7:

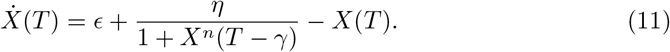

This equation has four parameters (*η, γ, ϵ*, and *n*). As outlined below, much of the behavior can be understood from just two, *η* and *γ*.

Since leakage must be small relative to regulation (i.e., *ϵ/η* ≪ 1) for regulation to be strongly effective (“activated” rate much greater than “non-activated” rate), we will focus here on the case with no leakage (*ϵ* = 0). We treat non-negligible leakage in the supplements (see Appendix S3). Note that in general Eq 11 has no closed-form solution. What we can do, however, is solve for the phase boundaries, or bifurcations, of this equation. This tells us the types of dynamical behavior of *X*(*T*) to expect in different parameter regions. To do this, we first determine the fixed points of the system described by Equation 11, and linearize around these fixed points. Next, we can determine the eigenvalues of the corresponding linear dynamical system and, by solving for when the real parts of these eigenvalues equal zero, we identify the system’s phase boundaries. We then also confirm these analytical results with numerical simulations.

#### Fixed points do not depend on delays

Fixed points are values *X*(*T*) = *X*\* for which *X* does not change with time. By definition, we can substitute 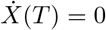 and *X*(*T*) = *X*(*T* − *γ*) = *X*\* into in Eq 11, indicating that the production term must exactly cancel out the degradation term (Fig 4D). For repressors (*n* > 0) there is only a single fixed point regardless of the parameter values so long as they remain biological. For activators (*n* < 0), there can be 1, 2, or 3 fixed points (2 fixed points is a border case for −*n* > 1). Explicitly, the fixed point values are given by

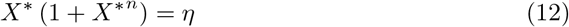

or *X*\* = 0 (for activators only). Again, we assumed *ϵ* = 0 for simplicity. Note that these fixed points only depend on 1/*η* = *βk/α* and *ϵ/η* = *α*_0_/*α*, and have no dependence on the delay time *τ*. This is as it should be, because the fixed points are fixed in time by definition, and should have no dependence on previous times.

#### Linearization is sufficient to determine bifurcations in qualitative behavior

By focusing on small disturbances from the fixed points *δX*(*T*) = *X*(*T*) − *X*\* ≪ 1, one can linearize Eq 11 (see Appendix S2) around its fixed points. Assuming a solution *δX*(*T*) = *A* exp(λ*T*) yields a transcendental equation for the eigenvalues λ termed the characteristic equation of Eq 11:

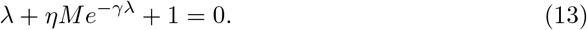

The constant function of the fixed point value *M*(*X*\*) is defined as

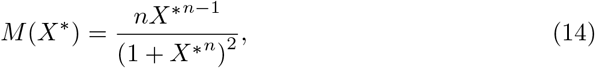

whose sign is given by *n* (positive for repressors and negative for activators), since all the terms besides *n* are non-negative.

To determine whether the fixed points are stable, we solve for the conditions in which the eigenvalues λ cross the imaginary axis, from negative to positive. This is true when Re λ = 0, either at λ = 0 (termed a saddle-node bifurcation) or at λ = *iω* (termed a Hopf bifurcation).

#### Saddle node bifurcations determine bistability in autoactivators

For the saddle-node case (λ = 0), Eq 13 and Eq 12 reduce to (see Appendix S2):

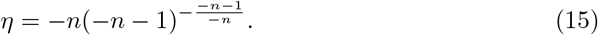

which gives real, positive (i.e., biologically meaningful) *η* for *n* < −1, meaning the saddle-node bifurcation occurs only for autoactivation and not for autorepression. Note that Eq 13 for λ = 0 is equivalent to the condition that the degradation rate (normalized here to 1) and the production rate as functions of *X*\* are tangent (i.e., the two lines in Fig 4D activator case are tangent to one another at a single point). If the production rate as a function of the fixed point is any steeper (*η* any larger), two non-zero fixed points are created, a stable one with high *X*\*, and an unstable one with lower *X*\*. For −*n* > 1, this means the system becomes bistable, since the state *X*\* = 0 is always stable.

We can view Eq 15 as a function of −*n* (see Fig. S2). As a reminder, *n* < 0 for activators so the biological cooperativity is |*n*| = −*n*. From this perspective, the equation gives the value of *η* for onset of stability at a given *n*. The maximum value of this function is *η* = 2 at −*n* = 2, meaning the system is always bistable above *η* = 2 regardless of the cooperativity (as long as −*n* > 1). The minimum is *η* = 1 which is the last real value of the function at −*n* = 1, meaning bistability never occurs for −*n* < 1 or for *η* < 1; for very large negative n, the bistability boundary also approaches *η* = 1. Because the function is non-monotonic, it is also possible (although biologically perhaps unrealistic except on an evolutionary timescale), to hold a value of 1 < *η* < 2 and decrease *n* to a point where the bistability is lost, and then decrease *n* still further until bistability is gained again. This can all be seen in the bifurcation curves in Fig. S2.

For the limiting case −*n* =1, λ = 0 implies *X*\* = 0 and *η* = 1, which is actually the limit of Eq 15 as −*n* approaches 1. Then for *η* < 1, the origin is stable, and for *η* > 1, only one new stable high-*X* fixed point is created, and the origin becomes unstable. For 0 < −*n* < 1, the value of *M*(*X*\* = 0) approaches −∞ as *X*\* approaches 0, which makes λ > 0 near the origin unless *η* = 0 (no production of *X*; see Eq 13). The origin is thus unconditionally unstable, and there is one other fixed point which is stable (unless *η* = 0, in which case the origin is the only fixed point). Finally, for *n* = 0, the regulation term is really just a leaky expression term, and we have *η* = 2*X*\*, which is the only fixed point, and which is unconditionally stable. The origin is not a fixed point at all, as it will not be for *n* > 0, the repressor case.

#### Hopf bifurcations determine oscillatory behavior in autorepressors

For the Hopf bifurcation (λ = *iω*), Eq 13 and Eq 12 result in two equations, one for the real parts of Eq 13 and one for the imaginary parts (see Appendix S2).

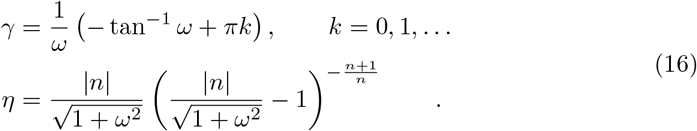

Equations 16 represent a series of curves parameterized by *ω*, but there is a single outermost curve at *k* = 1 (*γ* is always negative if *k* = 0) giving real, positive values for both *γ* and *η*. For values of *γ* and *η* greater than this boundary, *X* oscillates in a stable limit cycle. This boundary exists for both positive and negative *n*; however, for negative *n* (activators), the boundary lies entirely above the bistability boundary, which can be found by taking the limit as *ω* → 0 (for which *γ* → ∞), and equals exactly the bistability boundary given by Eq 15. This means that at least one eigenvalue is positive and real, so no oscillations are observed for activators. For repressors, any value of *n* > 1 will have a boundary (see Appendix S2) given by Eq 16 beyond which the system oscillates. This boundary has both horizontal (*η*) and vertical (*γ*) asymptotes, given as:

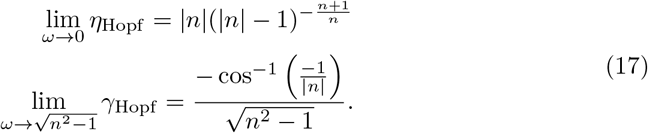

Thus there is both a minimal required regulation strength regardless of delay, as well as a minimum required delay regardless of regulation strength in order to achieve oscillations. Note though that the vertical (*γ*) asymptote approaches zero as *n* → ∞, meaning a very large regulatory strength could achieve oscillations even with minuscule delay if cooperativity is extremely steep. No oscillations occur for *n* ≤ 1, since oherwise Eq 16 would give non-real *η*.

The oscillation period can be approximated by linearization near consecutive maxima and minima of the oscillation (see Appendix S2). This method yields a period of approximately 2(*γ* + 1), or 2(*τ* + 1/*β*) in dimensionful terms (compare also to [26]). Biologically, this means that the concentration *X* is pushed from high to low (or vice versa) after the delay time (*τ*) has elapsed and the concentration has equilibrated to its new value via degradation (1/*β*).

Damped oscillations are expected when the largest eigenvalue satisfying Eq 13 has non-zero imaginary part. This is never true for *n* < 0, and is true for *n* > 0 above the curve denoted “spiral boundary” in Fig 1E and Fig. S1. This curve is given as follows:

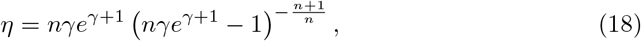

which is derived in Appendix S4. It approaches *η* = 0 for large delay and has a vertical asymptote at *nγϵ*^*γ*+1^ = 1, so there is some non-zero delay that has non-oscillating decay for any strength *η*.

Putting together the derived boundary curves yields a *η*-*γ* phase space for each *n*, displayed in Fig 4E and Fig. S1 along with simulations showing that this analytical treatment matches the dynamical behavior.

#### Leakage prevents oscillations and bistability

With leakage present (Appendix S3), the phase space becomes 3-dimensional, with Fig 4E charts as 2-dimensional slices at *ϵ* = 0. With small leakage, this 2-dimensional slice changes only slightly in a quantitative manner, but the qualitative portrait remains the same. Large leakage, on the other hand, effectively drowns out the regulation, preventing both oscillations (for negative autoregulation) and bistability (for positive autoregulation). The full treatment with leakage and the corresponding phase portraits are presented in the supplements (Appendix S3, Fig. S3).

Note that the explicit delay present in the model allowed us to capture *an entire phase space of oscillators, switches, and housekeeping (unregulated) genes in a single equation with only two important free parameters*. This phenomenological model can thus capture a large range of possible situations with relative simplicity. We arrived at these results despite the added complexities of DDEs relative to ODEs without having to analytically solve the differential equations. This same procedure can be applied to more general regulatory networks, which we examine next.

#### Key results for autoregulation

1. The regulatory strength *η*, leakage *ϵ*, and cooperativity *n* determine the available steady states, while the time delay *γ* determines the dynamics but does not affect steady state behavior.
2. As analyzed in Appendix S3, the leakage *ϵ* is easily incorporated into these phase boundaries as well, resulting in a 3D phase diagram for each *n*. Leakage primarily serves to drown out the regulation, preventing oscillations and bistability at high *ϵ*.
3. The cooperativity *n* is incorporated into the analytically derived phase space boundaries. No specific parameter values had to be assumed, and no linear approximations needed to be made to the nonlinear regulation.
4. The entirety of the phase space for a self-regulating gene, both activator and repressor, are contained within a single phase space of only two primary free parameters (*η* and *γ*), while *n* serves a more modulating role as long as |*n*| > 1. (There is also no chaos, as can be proven [42].)
5. Some cooperativity is necessary (i.e., |*n*| > 1) in order to realize bistability or oscillations (i.e., these behaviors are impossible for |*n*| ≤ 1). For a range of regulation strength (1 < *η* < 2), the bistability of self-activation is not achievable for a range of cooperativities (see Fig. S2).
6. Damped oscillatory behavior occurs in autorepressors for small but non-zero delay and regulation strength, meaning that regulation will overshoot or undershoot the target fixed point for some initial conditions. No damped behavior is ever possible for autoactivators, meaning there can be no overshoot.
7. Sustained oscillations require a minimum delay greater than zero, since the Hopf bifurcation curve has both vertical and horizontal asymptotes. Higher cooperativity decreases the required minimum delay, which approaches zero (albeit for infinitely high regulation strength) for step-function regulation (infinite cooperativity).
8. Both oscillations and bistability require a minimum regulatory strength, given by the saddle-node bifurcation line in autoactivators (bistability) and by the horizontal asymptote in autorepressors (oscillations).
9. Oscillation period in autorepression is approximately given by twice the delay time plus the degradation time.

### Motif III: Logic

So far we have only discussed regulation in which a single input regulates a single output. We represent this 1-variable regulation using a delayed Hill function. To represent networks in which two or more inputs regulate a single output (Fig 5), we must choose a 2-variable function. Many such functions are used in biology to describe specific mechanistic scenarios [71].

**Fig 5.**
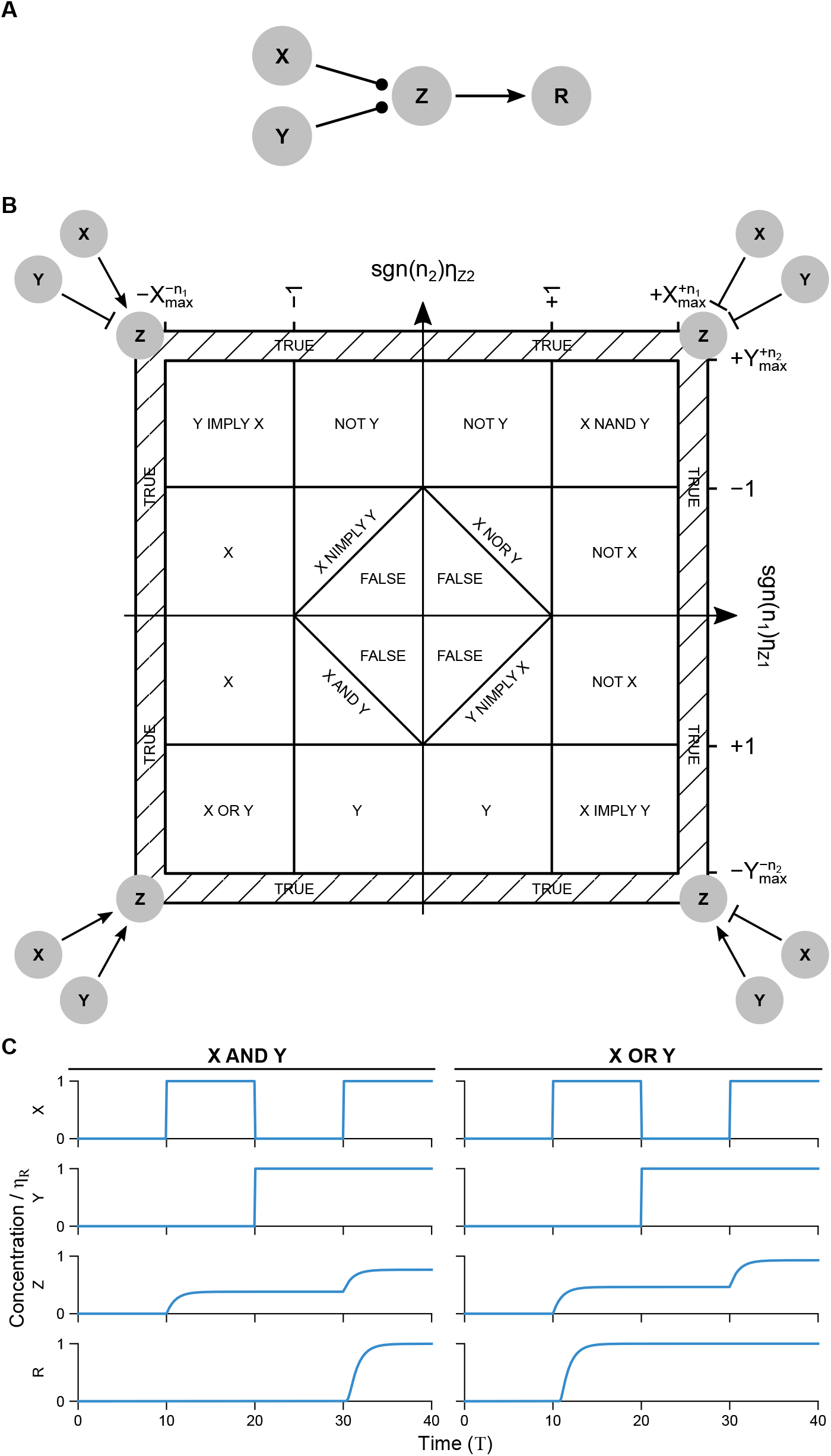
A simple approximation for digital logic using a sum of Hill terms recapitulates all monotonic logic functions in a single parameter space. (A) A prototypical regulatory network involving logic where X and Y both regulate Z, which must integrate the two signals using some logic before it can in turn activate a downstream reporter R. (B) Parameter space showing regions where regulation approximately follows 14 of the 16 possible 2-input logic functions depending on the strength of the two single-variable Hill regulation terms (*η*_*Z*1_: regulation of Z by X, and *η*_*Z*2_: regulation of Z by Y). Network logic can be smoothly altered by varying the parameters (*η*_*Z*1_, *η*_*Z*2_), with a change of sign in (*n*_1_, *n*_2_) required to switch quadrants. The bottom-left quadrant shows that very weak regulation in both terms leads to an always-off (`FALSE`) function, weak regulation in one arm only leads to single-input (X, Y) functions, strong regulation in both arms leads to an `OR` function, and regulation too weak in either arm alone to activate an output but strong enough in sum leads to an `AND` function. The other three quadrants are related by applying `NOT` to one or both inputs, with function names related by de Morgan’s law [72] `NOT`(X `OR` Y) = `NOT` X `AND NOT` Y. In particular, X `IMPLY` Y = `NOT`(X) `OR` Y, X `NIMPLY` Y = X `AND NOT`(Y), X `NOR` Y = `NOT` X `AND NOT` Y, and X `NAND` Y = `NOT` X `OR NOT` Y. Truth tables for all 16 logic gates are provided in Table S2 for reference. The two non-monotonic logic functions, X `XOR` Y and X `XNOR` Y, are those 2 of 16 not reproduced directly using this summing approximation. They can be produced by layering, e.g., `NAND` gates [72]. (C) Representative time traces for `AND` (*η*_*Z*1_ = *η*_*Z*2_ = 0.9) and `OR` (*η*_*Z*1_ = *η*_*Z*2_ = 1.8) gates with *n*_1_ = *n*_2_ = −2, *n*_3_ = −20, *η_R_* = *η*_*Z*1_ + *η*_*Z*2_. The function sgn(*n*) = +1 when *n* > 0, sgn(*n*) = −1 when *n* < 0.

One class of 2-variable functions that has found widespread use in describing natural [2, 65,73–76] and synthetic [77–81] biological networks is the logic-gate description. In analogy to digital electronics [72], logic gates in biology determine whether the output is at high concentration (“on”) or at low concentration (“off”) depending on whether the inputs are on and off. For example, commonly found two-input logic gates include the `AND` gate (output on if and only if both inputs are on) and the `OR` gate (output on if either or both inputs are on).

Despite the importance of specifying 2-input functions, these functions are not always drawn explicitly in biological schematics (e.g., Fig 5A) [2,65,73,74] as they are in digital electronics [72]. In keeping with this lack of convention, we will not draw logic gates in our schematics, but any time regulation proceeds from two inputs to one output a logic function is implied. In this section we provide a specific function that covers 14 out of 16 possible 2-input logic operations, and show that these operations form a continuous 2D parameter space.

#### A summing function characterized by two parameters reproduces all 2-input monotonic logic gates

To begin our analysis of logic functions, we start by writing down some nondimensionalized equations corresponding to Fig 5A. We assume that the degradation constants (*β*) for *Z* and *R* are equal for simplicity, and that there is no leakage. In addition, we allow the input genes to follow arbitrary dynamics, in keeping with their role here as arbitrary network inputs.

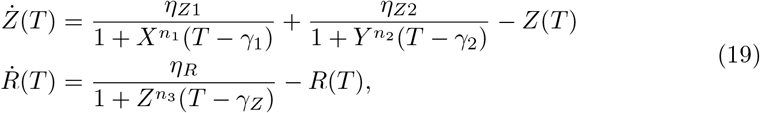

and where the notation as before is with *n_i_* > 0 for repressors and *n_i_* < 0 for activators. The nondimensionalization is the same as for autoregulation (see Appendix S5). In particular, *Z* = *z/k_z_* is concentration of *z* normalized to its half-maximal input to *r*. *R* = *r/k_r_* for arbitrary constant *k_r_* that could refer to, for example, the half-maximal input of *r* on some downstream gene if *r* is a transcription factor.

We have made a choice to describe the regulation of *Z* due to *X* and *Y* using separate additive Hill terms. Other formulations are possible. For example, the product of two Hill coefficients can be used [82] or *X*^*n*_1_^ and *Y*^*n*_2_^ could be summed in the denominator of a single Hill function [65] or some combination thereof [75,76]. These all represent subtly different biological situations. For example, Eq 19 most closely describes two independent promoters for *Z*, one activated by *X* and one by *Y*, while combining *X* and *Y* in the denominator of a single Hill function most closely describes two transcription factors that bind a single promoter for *Z*. The benefit of using two separate Hill functions is that it is mathematically tractable, and as we will see below, can describe many logic functions simply by tuning of regulatory strengths.It is limited, however, in that it necessarily has multiple states for *Z*, which are only rectified to a set of binary states in *R* (Fig 5C). It also deals poorly with ratiometric inputs, as we will note in the next section on feedforward motifs. Previous work has shown that a single combination of terms can similarly produce almost as many logic functions [75,76]. The analysis done here using a sum of Hill terms for this and later motifs can be reproduced with few changes using, for example, multiplicative terms (see Appendix S5).

Based on the normalization chosen for nondimensionalization, we can make a surprisingly simple characterization of the logic resulting from this type of regulation based on the idea of dynamic range matching [83]. Every regulator in Eq 19 appears in a sum with unity in the denominator of the Hill function for the protein it is regulating. This means that, in order to be an effective regulator, the normalized value of the regulator must be able to go both above and below 1. This is in fact the reason that we explicitly include the reporter *R* in this discussion. Defining the output of the logic circuit as “on” or “off” requires some reference point of activity between on and off, which is provided as *k_z_*, the half-maximal input of *Z* to *R*. After normalizing *Z* = *z/k_z_*, this midpoint becomes *Z* = 1.

We assume that as effective regulators, *X* and *Y* can both take values above and below 1. We will further assume that *Z* can also take on values below 1, in order to be a good regulator of *R*. For this to be true, *η*_*Z*1_ ≪ max(*X*^|*n*_1_|^) and *η*_*Z*2_ ≪ max(*Y*^|*n*_2_|^). If these conditions are not met, *Z* will activate *R* regardless of the inputs from *X* and *Y*, indicated by the areas marked TRUE in Fig 5.

Let us say that *X* and *Y* settle on steady-state values *X* * ≡ *η_X_*, *Y*\* ≡ *η_Y_*. If we assume these values arbitrarily, then they serve as the inputs to the logic gates. A value of *η_X_* or *η_Y_* significantly greater than 1 is “high” (true, 1) in that it is seen as above the unity regulatory threshold in the equation for 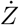, and “low” if much less than 1, for the converse reason. We then want to look at whether *Z*^*^ is greater than or less than 1, which similarly serve as “high” (true, 1) and “low” (false, 0) values, respectively, as used in 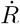.

For example, the case where *n*_1,2_ > 0 (two repressors), the steady state value of *Z* is given as a function of *η_X_, η_Y_, η_Z1_*, and *η*_*Z*2_, as shown in the column labeled *Z*\* in Table 1. We can then look at the value of *Z*\* when the values of the inputs *η_X_* and *η_Y_* are much smaller or much larger than 1, under the above-given assumptions that 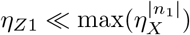 and 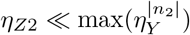. For example, if both *η*_*Z*1_ > 1 and *η*_*Z*2_ > 1, then the output simplifies to the last column in Table 1. The table overall then represents the truth table for a NAND gate. Calculations for all conditions are given explicitly in Table S1.

**Table 1.** `NAND` Logic function for two repressors based on Eq 19.

| $\eta_X$ | $\eta_Y$ | $Z^*$ | $Z^*$ for $\eta_{Z1,2} > 1$ |
| --- | --- | --- | --- |
| $\ll 1$ | $\ll 1$ | $\eta_{Z1} + \eta_{Z2}$ | $> 1$ |
| $\ll 1$ | $\gg 1$ | $\eta_{Z1} + \eta_{Z2}/\eta_Y^{n_2}$ | $> 1$ |
| $\gg 1$ | $\ll 1$ | $\eta_{Z1}/\eta_X^{n_1} + \eta_{Z2}$ | $> 1$ |
| $\gg 1$ | $\gg 1$ | 0 | $< 1$ |

By simply changing the parameters *n*_1_, *n*_2_, *η*_*Z*1_, and *η*_*Z*2_, the logic function can be changed. If instead of having *η*_*Z*1_ > 1 and *η*_*Z*2_ > 1, they are both < 1 but their sum > 1, then the table looks like a `NOR` gate. If their sum is < 1, then the table looks like a `FALSE` gate (i.e., output is always low). If *η*_*C*1_ > 1, *η*_*C*2_ < 1, then it looks like `NOT` A. If *η*_*C*1_ < 1, *η*_*C*2_ > 1, then it looks like `NOT` B. If we relax the assumptions that *η*_*Z*1,2_ are below the maximum of the inputs, then it looks like a `TRUE` gate (i.e., large enough *η*_*Z*1_ or *η*_*Z*2_ implies the output is always “on”).

This logic can be extended into all possible combinations of activators and repressors, and combined into a convenient chart, shown in Fig 5B. The chart is symmetric on interchange of the two inputs (*η*_1_ ↔ *η*_2_, *n*_1_ ↔ *n*_2_). It is also anti-symmetric (that is, application of logical `NOT`) by conversion of an activator to a repressor or vice versa (*n_i_* → −*n_i_*). Fourteen of the 16 possible two-input logic functions are represented in the chart. All fourteen are monotonic functions of *X* and *Y*, in that *Z*(*X, Y* =1) > *Z*(*X, Y* = 0) and *Z*(1, *Y*) > *Z*(0, *Y*). The two non-monotonic gates, XOR and XNOR are not represented by this simple logic motif, because summing (addition of Hill terms) is monotonic. However, they can be constructed from connecting several of these gates [72, 83].

#### The logic parameter space can be divided into AND-type, OR-type, and single-input functions

Only the 8 regulatory functions on the diagonals of Fig 5 (excluding `FALSE` and `TRUE`) make use of both inputs. These eight can be further divided into positive or negative regulation on each arm (4 possibilities) in conjunction with an `AND` or `OR` gate (8 possibilities total). Specifically, “`OR`-type” logic applies when both *η*_1_ > 1 and *η*_2_ > 1 (`OR, NAND`, and both `IMPLY` gates), and “`AND`-type” logic applies when both *η*_1_ < 1 and *η*_2_ < 1 (`AND, NOR`, and both `NIMPLY` gates). This can be seen mathematically by using Boolean logic simplification. For example, *X* `NOR` *Y* = `NOT` *X* `AND NOT` *Y*. Similarly, *X* `NAND` *Y* = `NOT` *X* `OR NOT` *Y*.

It is important to note that this logic scheme is an approximation, and in reality the sum of two Hill terms provides a form of fuzzy logic [84]. That is, if the inputs *X* or *Y* are close to 1, or if the regulatory strengths *η*_*Z*1,2_ are close to 1, then *Z* will also be close to 1 for some input combinations. Thus the boundary lines in Fig 5 are soft, and become exact as the cooperativities become large. Linearizing the *R* fixed point around *Z* =1, one can conclude that |*n*_3_| ≳ 2(1 − *ε*)/*δ* is required for an input *δ* above 1 to yield an output *ε* below 1. Thus for highly digital logic, it must at least be true that |*n*_3_| ≫ (*δ* = 1, *ε* = 0), and in the limit |*n*_3_| → ∞ (*δ* → 0) *R* becomes a step function, as in the case of exact (rather than fuzzy) logic.

Our analysis thus far of the logic motif (Fig 5) with two separate inputs has focused on steady state outcomes, and therefore only exhibited trivial dependence on delays. Since there is no feedback in this network, we can mathematically time-shift the definition of *X*(*T*) and *Y*(*T*) independently so that no delays remain in either input. This changes however when the two inputs *X* and *Y* are not independent. As a simple example, if *X* and *Y* represent the same input source (i.e., *X* ≡ *Y*), then a difference of delays between the two regulation arms cannot be shifted out entirely. This type of regulation, in which a single input regulates a single output through two arms, is the feedforward motif, which we turn to next. We will show in particular that this inability to remove the difference of delays is essential to the motif’s behavior.

#### Key results for logic

1. Logical functions can be determined by comparing the strength of regulatory inputs to the half-maximal input of the regulated output. In non-dimensional form, all strengths can be compared to 1.
2. Both the strengths of the inputs and the strength of the output on a reporter must be considered to derive a logic function.
3. Boolean logic gates can be concisely described in a simple two-dimensional parameter space given by sgn(*n*_1_)*η*_1_ and sgn(*n*_2_)*η*_2_.
4. Out of 16 possible 2-input logic gates, 14 are generated by a sum of two Hill functions; only the two non-monotonic logic gates (`XOR` and `XNOR`) are not.
5. From an evolutionary perspective, the fact that all these logic gates fit in a single parameter space implies easy conversion of `AND`-type logic and `OR`-type logic, with a sign flip required on Hill coefficients needed to add `NOT`s to the logic functions. This is similar to results for other functions describing a subset of logic gates [75, 76].
6. For independent inputs, logic has only trivial dependence on delays, which shift one input relative to another in time.

### Motif IV: Feedforward loop

The feedforward motif (see Fig 6) is a non-cyclic regulation network in which an input, *X* regulates an output, *Z*, via two separate regulation arms. One arm is “direct,” in which *X* regulates *Z* in a single step, while the second arm is “indirect,” with *X* regulating an intermediate *Y*, which in turn regulates *Z*. In this way, the first arm “feeds forward” past the cascade (see Fig 6A-B). The motif is found commonly in biological networks [11,85], comprising about 30% of three-gene regulatory interactions in transcriptional circuits [86].

**Fig 6.**
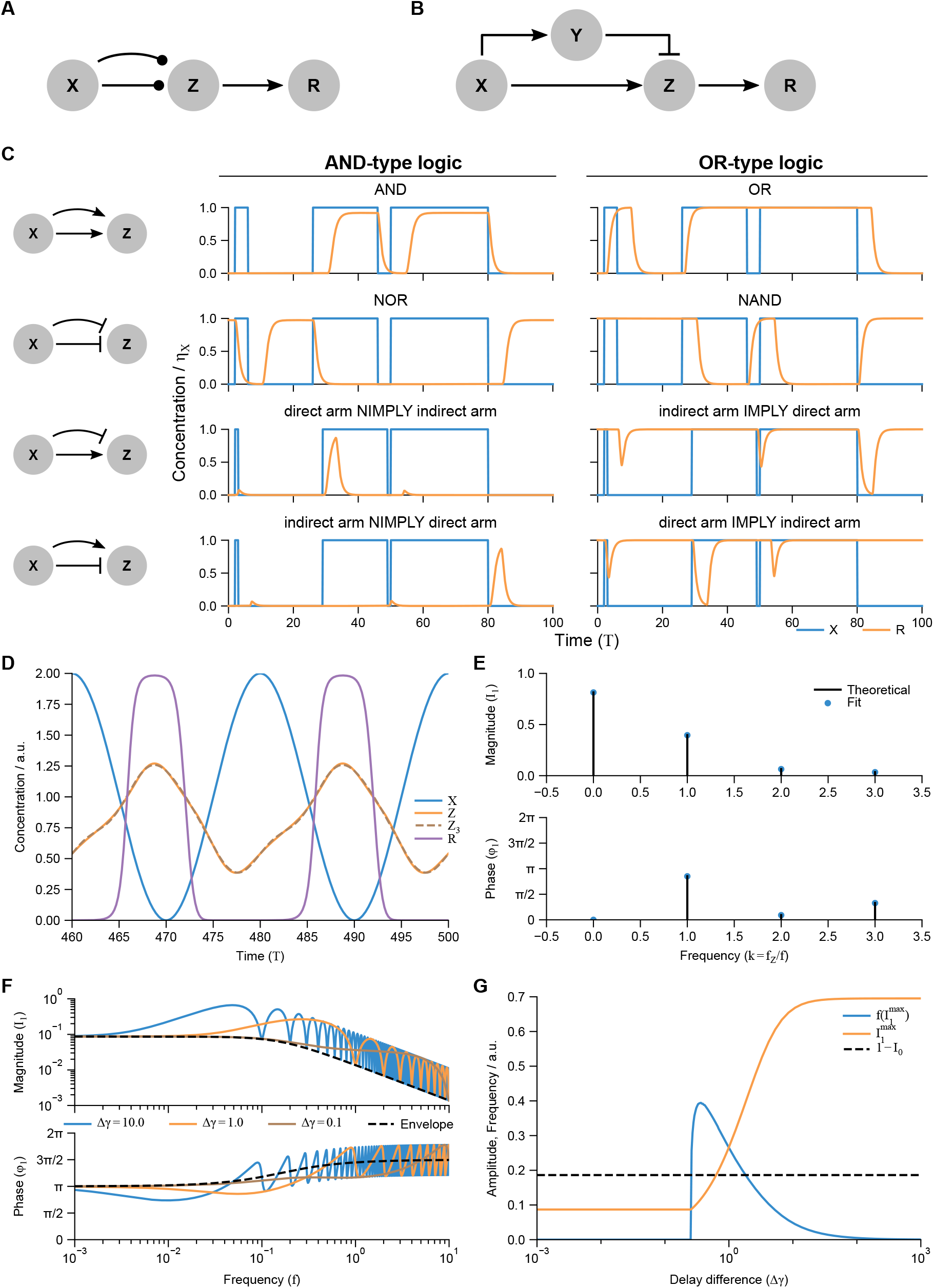
The feedforward network motif owes its primary functions to a difference in regulatory delays. (A) The feedforward motif with delays, in which a single output *Z* is controlled by an input *X* via two regulatory arms with differing delays. The straight, short arrow represents the “direct arm” with delay *γ*_1_ and the longer, curved arrow represents the “indirect arm” with delay *γ*_2_ > *γ*_1_. (B) The ODE model for an incoherent (type 1) feedforward motif, one of 8 possible networks in which the intermediate gene *Y* is modeled explicitly in the indirect arm. (C) Simulations of the four feedforward motifs with AND-type and OR-type logic (see Fig 5 and Table S2) in response to short and long gain and loss of input signal. Blue curves: inputs (*X*), orange curves: reporter *R* activated with high cooperativity by *Z*. Note that the bottom two rows demonstrate pulse generation, while the top two rows filter short signals. (D) Response of an incoherent feedforward motif to oscillatory input after initial transients have died away. *Z*_3_ is a 3-frequency Fourier approximation of *Z* (see E). (E) Fourier decomposition of *Z* from (D) by Eq 30 and by a numerical fit to the data in (D). (F) Frequency scan (Bode plot) of (D) for 3 values of Δ*γ*, with the theoretical envelopes from Eqs 31,32. (G) The maximum amplitude of the motif in (D) over a range of Δ*γ* and the corresponding frequencies at which the maxima occur. *Z* goes above 1 (activation threshold for *R*) for a small range of Δ*γ*. For (C-F), *η*_1_ = 0.9, *η*_2_ = 0.7, *n*_1_ = 2, *n*_2_ = −2, *n*_3_ = −20, *η_R_* = 2, *A* =1. For (D), *f* = 0.05, Δ*γ* = 4.

The different combinations of activation and/or repression from the two terms generate what are termed “coherent” (regulation arms are both activating or both repressing the output) or “incoherent” (one activation arm and one repression arm) feedforward motifs. These behaviors additionally depend on the logic (Fig 5) associated with how the downstream element responds to its inputs [2, 61, 65]. Standard classification of feedforward loops based on which regulatory arms are activating or inhibiting have been used in the literature (see Table 2). Here we show how a DDE model maps onto these characterizations and provide analytical solutions to pulse and frequency responses of feedforward loops.

**Table 2.** Standard classification of feedforward loops [61]. Each type of feedforward loop has two regulation arms (see Fig 6A), one fast-acting (short delay) and one slow (larger delay). Each arm either activates or represses the output. This leads to the four types of feedforward loops listed here. ODE models have twice as many distinct feedforward loops as the DDE model, because the slow regulation arm in the DDE model has only one step instead of two, and hence two designations are referenced in the leftmost column for each type.

| Feedforward loop designation | Fast regulation | Slow regulation |
| --- | --- | --- |
| Coherent types 1, 4 | Activation | Activation |
| Coherent types 2, 3 | Repression | Repression |
| Incoherent types 1, 4 | Activation | Repression |
| Incoherent types 2, 3 | Repression | Activation |

#### Feedforward dynamics depend only on the difference in time delay between two regulatory arms and their 2-input logic

ODE models of this motif provide equations for each of the multiple regulators in such a pathway, but we show here that the essential ingredients are the production of a difference in delays between the two inputs to *Z*, in addition to the logic function between the two inputs and the output *Z*. With a DDE model, we can drop the intermediate *Y*, focusing just on the input and output proteins, *X* and *Z*, using two regulating terms with different delay times (Fig 6A-B).

Looking at Fig 6A, we can write down a simple equation for the regulation of *Z*, by setting *Y* = *X* in Eq 19 (see Appendix S6):

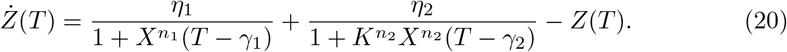

Because each regulation term is controlled by the same input (*X*), we cannot in general normalize the half-maximal input to 1 in both terms as we did for logic (Eq 19); *K* = *k*_1_/*k*_2_ represents the ratio of the two input scales. We will assume that the level of *X* has arbitrary dynamics, as in our discussion of logic in the previous section.

If one shifts the input variable backwards in time by *γ*_1_ 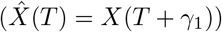, we see that the behavior of the feedforward equation actually depends only on the difference in regulatory delays, Δ*γ* = *γ*_2_ − *γ*_1_ rather than each delay independently:

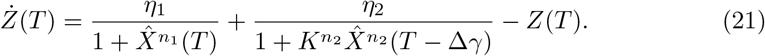

For clarity we will from now on refer to 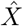 as *X* without the hat unless specified.

To understand the behavior of Eq 21, we will analyze the output of *Z* to two types of input *X*: a step input, and a continuous oscillation. The response of a system to these two types of inputs, respectively termed step response and frequency response, are often used in control theory to characterize an input-output relation [87].

#### Step-pulse response of feedforward motifs can be solved analytically and demonstrates filtering and pulse-generation behaviors

We considered a step input (Fig 6C) to include both a step on (from low to high *X*) as well as a step off (from high to low *X*):

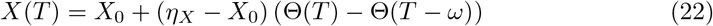

where the Heaviside function is defined such that Θ(*T*) = 1 for *T* < 0 and Θ(*T*) = 0 for *T* > 0. Eq 22 represents a square input pulse of width *ω* starting at *X*_0_ and reaching to *η_X_*. This is an on-pulse if *η_X_* > *X*_0_ and an off-pulse if *η_X_* < *X*_0_.

The lack of feedback in feedforward loops and the fact that the square-pulse input of Eq 22 takes on only two values simplify the form of Eq 20 significantly, allowing it to be solved explicitly (see Appendix S6). The solution is:

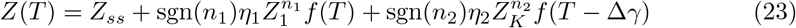

in which

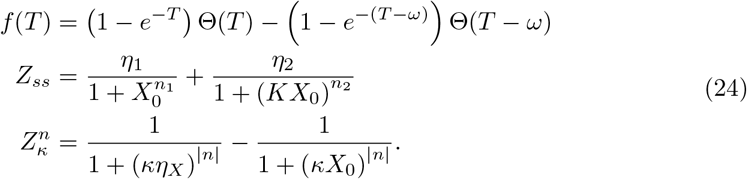

Here, *Z_ss_* is the steady-state value of *Z*, 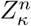 are the magnitudes of deviation from steady state due to each arm, and *f*(*T*) specifies where the responses from the two regulation arms are active. In terms of the original *X* (as opposed to the time-shifted 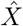), the output in Eq 24 is shifted to the right in time by *γ*_1_ relative to the input original input *X*(*T*).

There are several interesting results to note in this equation. First, the magnitude of the Hill coefficients only serve to modulate the strength of the input, *η_X_* and *X*_0_; they have no other effect because the input already has infinitely steep transitions between on and off. Likewise, the ratio of half-maximal input levels in the two arms (*K* = *k*_1_/*k*_2_) only modulates the strength of the responses.

Second, the *η* values only appear in the baseline steady-state value *Z_ss_* (determining the logic of the steady state output) and in the combinations sgn(*n_i_*)*η_i_*, which as noted in Fig 5 are the parameters dictating the regulation logic. Based on Fig 5, there are 8 important logic functions which respond significantly to both inputs and are therefore important for feedforward loops: four `AND`-type functions and four `OR`-type functions.

Third, the function *f* only serves to indicate where the responses from the two regulation arms are active. If *ω* ≫ 1, then *f* equals 0 at the beginning and end of the pulse (*T* = 0, *T* = *ω*) and equals 1 in the middle of the pulse (0 ≪ *T* ≪ *ω*). It is always zero outside the pulse. All responses, contained in the function *f*, decay with time, so the system both starts and ends at the steady state *Z_ss_*. That is, feedforward responses must be driven externally, unlike autoregulation. Overall there are four responses to the input: beginning at *T* = 0, *T* = *ω*, *T* = 0 + Δ*γ*, and *T* = *ω* + Δ*γ*.

For coherent feedforward motifs, input pulses must be sufficiently wide (*ω* ≫ 1) to fully activate the output *R*. This implies that coherent feedforward motifs act as filters against short signals, either short on signals (`AND` and `NAND`) or short off signals (`OR` and `NOR`). The delay difference Δ*γ* determines how quickly the response starts (`AND` and `NOR` logic) or returns to baseline once the pulse is over (for `OR` and `NAND` logic), so that large Δ*γ* maintains a response long after the input pulse has subsided.

For incoherent feedforward motifs, the output *R* is only fully activated if the pulse is wide (*ω* ≫ 1) and the delay difference is also wide (Δ*γ* ≫ 1), in which case a pulse of output is formed. The output pulse is formed on either the on-step (slow arm `IMPLY` fast arm, and slow arm `NIMPLY` fast arm) or the off-step (fast arm `IMPLY` slow arm, and fast arm `NIMPLY` slow arm). This means that incoherent feedforward motifs act as pulse generators. The delay difference Δ*γ* determines the minimum width of input pulse that will produce a response in *R*.

A limitation in this description of feedforward loops is due to our choice of sum-of-Hills logic function. It has been shown that incoherent feedforward motifs show fold-change detection (FCD) for large input ranges *X* ≫ 1 [71], in which the response to an input s is not dependent on the absolute values before (*X*_0_) and after (*η_X_*) the step input, but only on the ratio (*η_X_*/*X*_0_). This is not true for the current description. One can see this by replacing *X* in Eq 21 with *F* * *X*_0_, where *F*(*T*) = *X*(*T*)/*X*_0_ is the ratiometric input change. Assuming *X* ≫ 1, there is an overall prefactor *X*_0_ on which the dynamics of *Z*, and in turn *R*, depend. An alternative incoherent logic function such as *X*(*T*)/(1 + *X*(*T*) + *X*(*T* − Δ*γ*) automatically has *X*_0_ drop out for large *X* with the same substitution *X* = *X*_0_*F*. Thus, while the sum-of-Hills logic framework works well for logical behavior, it is not ideal for describing ratiometric scenarios.

Many of the behaviors about feedforward motifs described in this section corroborate earlier results from ODE models of feedforward loops [2,2,61,61,65,71,85,86,88,89], but are derived here with just a single equation. In particular, it is well known that coherent feedforward loops have filter capabilities and that incoherent feedforward loops can act as pulse generators [2,61,65] or stripe generators when regulation is dependent on space instead of time [90,91]. Furthermore, we can clearly demonstrate that the feedforward loop dynamics depend entirely on the difference of delays in the two regulation arms, a fact that can only be seen indirectly with ODE models [65].

#### Frequency response of feedforward motifs can be solved analytically and demonstrates low- and band-pass filtering capabilities

There are many examples in which biological regulatory networks encode information as the change in frequency of an oscillating input [92, 93]. It has been suggested that these signals can be more robust to noise than encoding information in the absolute concentration of inputs [94–96]. The fact that feedforward motifs can filter inputs based on the width of square pulses suggests that they may have more general frequency-filtering capabilities. We therefore include here as well an analysis of the frequency response of this motif to sinusoidal input (Fig 6D-G). We will show that the frequency dependence of the output follows the outline of a universal transfer function curve, independent of the logic or delay difference.

Instead of the step input analyzed above, here we consider a sinusoidal input

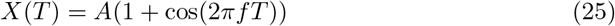

which oscillates between zero and twice the amplitude 2A at a frequency *f*. Since *X*(*T*) is periodic with frequency *f*, and Eq 20 has no explicit time-dependent terms, every term in the equation must be periodic with a fundamental frequency of *f* as well. We can thus decompose the first regulatory term in a Fourier series as follows:

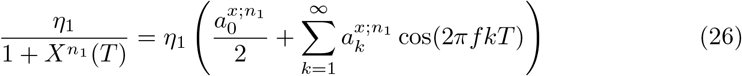

where the Fourier coefficients 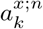 (with *n* = *n*_1_) are determined as

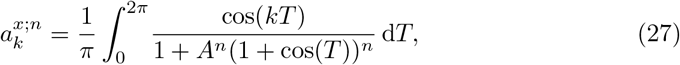

for all *f* > 0 (see Appendix S6). In the trivial case *f* = 0 all coefficients are zero except *a*_0_, which is 2 times the steady state value of Eq 21, slightly different in value than the limit as *f* → 0. Since the input is chosen to be a pure cosine, there are no sine terms in the Fourier expansion, which would otherwise add terms 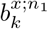 sin(2*πfkT*). The same Fourier decomposition can be done for the second regulatory term, by using *n* = *n*_2_ instead of *n*_1_, setting *T* → *T* − Δ*γ* in Eq 26, and replacing *A* → *AK* in Eq 27. We will not have need to discuss *k* = *k*_1_/*k*_2_ (not to be confused with the frequency multiplier *k*) further, but one may keep in mind that all values of 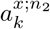 below reflect this modification in *A*.

We can similarly decompose *Z*(*T*) into Fourier components with unknown coefficients 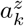 and 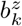:

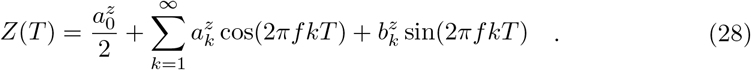

Because the regulatory terms and *Z* all have the same fundamental frequency, we can solve for the output coefficients 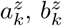 in terms of the input coefficients 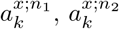 (see Appendix S6).

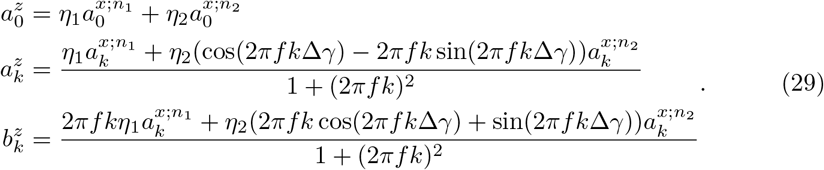

Finally, the Fourier decomposition can also be written in terms of magnitude and phase instead of cos and sin components. In this format (see Appendix S6),

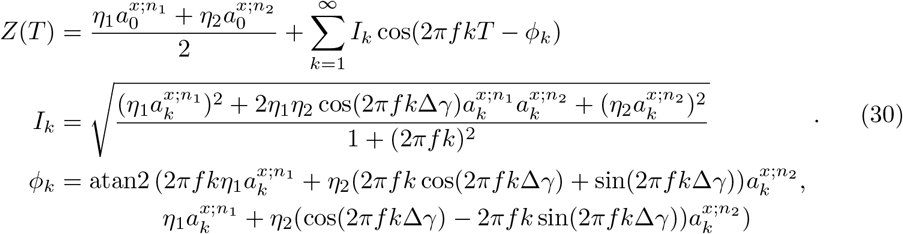

The magnitude of the output is symmetric to interchange of the two regulation arms (*η*_1_ ↔ *η*_2_, *n*_1_ ↔ *n*_2_), while the phase is not.

The magnitude *I_k_* decreases monotonically in *f*, as

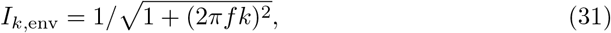

modulated by the sinusoidal numerator with period 1/*k*Δ*γ* (and which equals *I_k_*(*f* = 0) for frequencies an integer multiple of 1/Δ*γ*). Although harder to see explicitly in Eq 30, a similar behavior applies to the phase *ϕ_k_* as well. This can be found by looking again at frequencies an integer multiple of 1/Δ*γ*, in which case the phase reduces to

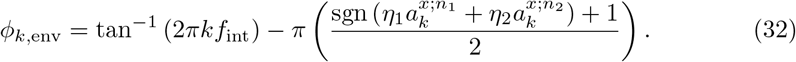

The amplitudes and phases as a function of frequency (also called Bode plots [87]) and their envelopes are plotted in Figure 6F.

At frequencies *f* that are integer multiples of 1/*k*Δ*γ*, the cosine in *I_k_* equals ±1. Because 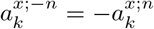 (see Eq 27, Appendix S6), the output magnitude *I_k_* then increases to a local maximum for coherent feedforward motifs (sgn(*n*_1_) = sgn(*n*_2_)) and decreases to a local minimum for incoherent feedforward motifs (sgn(*n*_1_) = sgn(*n*_2_)). The opposite holds for frequencies that are half-integer multiples of 1/*k*Δ*γ*. For the special case of perfectly balanced incoherent feedforward motifs (*η*_1_ = *η*_2_, *n*_1_ = −*n*_2_, *K* =1), the magnitudes *I_k_* decrease to zero for frequencies that are half-integer multiple of 1/Δ*γ*; otherwise, the maxima (for coherent) and minima (for incoherent) equal *I_k_* (*f* = 0).

Interestingly, the delay difference Δ*γ* only appears in the cosine and sine functions of *I_k_* and *ϕ_k_*. The monotonic envelopes that ignore these periodic effects for both the magnitude (Eq 31) and phase (Eq 32) do not depend at all on the delay difference Δ*γ* (Fig 6F). Thus for any delay difference, the frequency at which the magnitude envelope decreases to one-half its value at *f* = 0 is found at 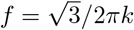, with an envelope phase shift of −*π*/3.

Despite this non-dependence of the envelope on Δ*γ*, the absolute maximum magnitude obtained for incoherent feedforward motifs does depend on Δ*γ*, as does the frequency at which this maximum response occurs. The dependence on delay difference is particularly strong near Δ*γ* = 1 (Fig 6G). This is because for Δ*γ* ≫ 1, the envelope decreases by half when *f* is many multiples of Δ*γ*, and thus past many local maxima, and so the local maximum occurs at 0 < *f* ≪ 1 with a maximum above *I_k_*(*f* = 0). For Δ*γ* ≪ 1, on the other hand, the envelope decreases by half before a single sinusoid is complete, so the maximum is at *f* → 0.

At Δ*γ* ≈ 1, the first sinusoid reaches its maximum while the envelope is descending most steeply, and the maximum occurs at *f* ≈ 1 with a value highly dependent on Δ*γ*. To activate *R*, *X* must go above 1 for at least part of the output oscillation, meaning that *I*_1_ must go above 1 − 〈*I*〉 = 1 − *I*_0_. Looking at Fig 6G, we see that for large Δ*γ* ≫ 1, this occurs for small frequencies, and the feedforward loop acts as a low-pass filter. However, there is a window of Δ*γ* ≈ 1 in which 1 − *I*_0_ is large, at modest frequencies (0.1 < *f* < 10), corresponding to the hump in magnitude, e.g., around *f* = 0.5 for Δ*γ* = 1 in Fig 6F. The incoherent feedforward motif thus acts as a bandpass filter for Δ*γ* ≈ 1, with ineffective regulation outside of modest frequencies.

For coherent feedforward motifs, the frequency at which the first minimum in output response occurs follows the same logic, but the maximum response is always found at *f* → 0. Since the maximum response is at low frequencies, coherent feedforward motifs act as low-pass filters.

#### Key results for feedforward loops

1. Results for feedforward loops modeled by DDEs essentially replicate results known from ODEs, but demonstrate explicitly and intuitively that their behavior depends only on logic and delay diffference between regulatory arms.
2. There are 8 possible feedforward motifs, corresponding to the 4 `AND`-type functions and the 4 `OR`-type functions on the diagonals of Fig 5B.
3. The only relevant parameters for feedforward motifs are the logic (sgn(*n*_1_)*η*_1_, sgn(*n*_2_)*η*_2_) and the difference in delays (Δ*γ*). The smaller delay merely leads to an overall shift in the output equal to that delay.
4. Step-pulse responses of feedforward motifs are composed of 4 independent responses: when the regulation arms are (1) both on, (2) both off, and (3,4) one on and one off.
5. The importance of synchronized (both on or both off) versus anti-synchronized (one on, one off) inputs on the two arms is determined by the difference in delays. High delay difference means inputs are usually seen as anti-synchronous by the output, while low delay difference means inputs are usually seen as synchronous.
6. Incoherent feedforward motifs generate pulses in response to step inputs, with the delay difference determining the minimum width of input pulse that will produce a response.
7. Coherent feedforward motifs act as filters against short signals, with delay difference determining how quickly the response starts (`AND`-type logic) or returns to baseline once the pulse is over (`OR`-type logic).
8. Fold-change detection can occur for feedforward loops, but requires logic functions that are not in the sum-of-Hill format.
9. Incoherent feedforward motifs can act as band-pass filters, responding most strongly when the input wavelength exactly mismatches the delay difference (half-integer multiples of 1/Δ*γ*).
10. Coherent feedforward motifs act as low-pass filters and respond to higher frequencies most strongly when the input wavelength matches the delay difference time (integer multiples of 1/Δ*γ*).

### Motif V: Multi-component feedback

Many natural networks include two-component loops [65], in which, for example, a transcription factor *x* regulates a transcription factor *y* which in turn regulates *x* (Fig 7A). In this section we analyze the effect of explicitly modeling two steps in a regulation loop as compared to modeling direct autoregulation.

**Fig 7.**
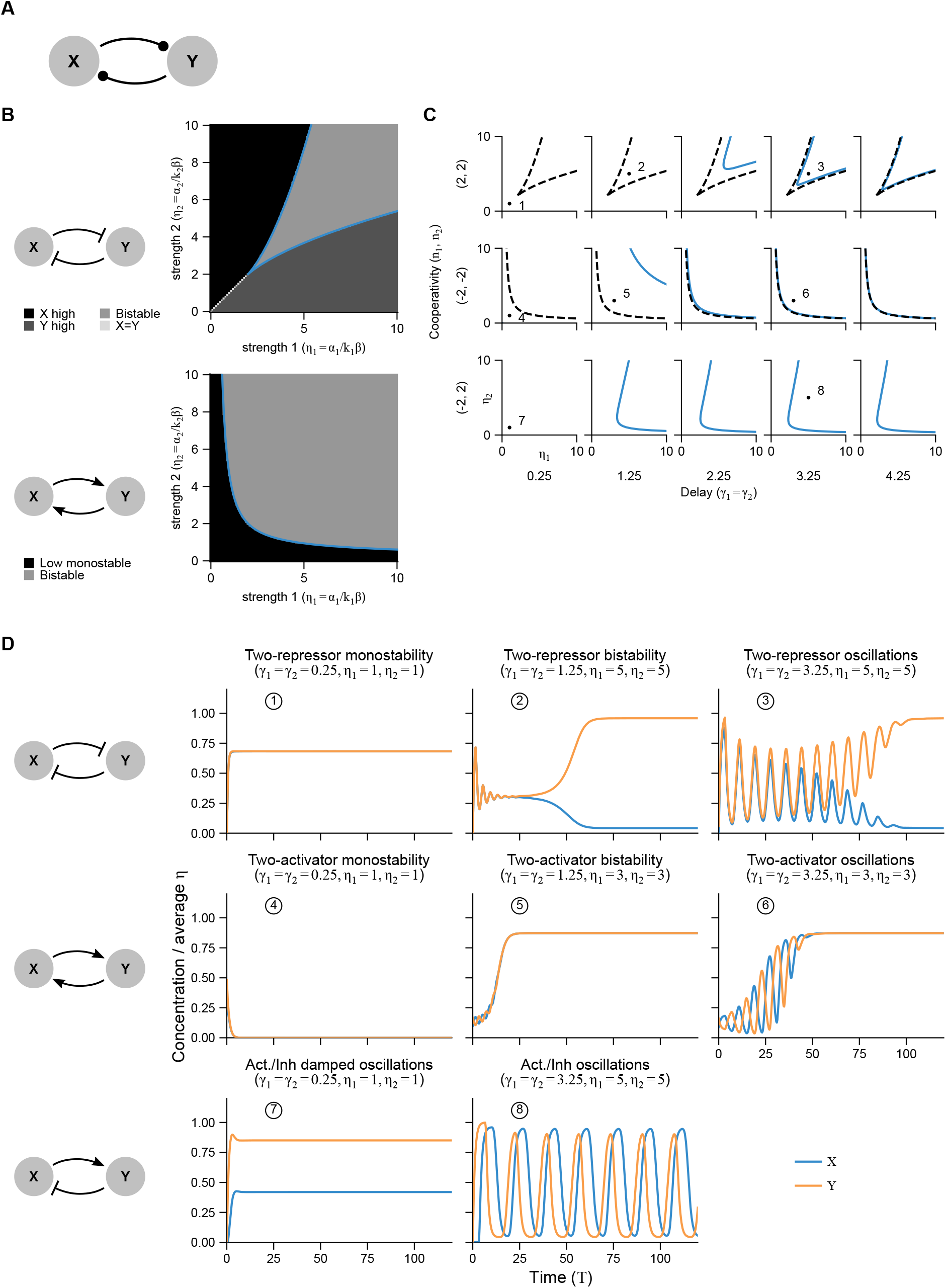
Two-component autoregulatory loops reproduce behaviors of autoregulation, but have additional behaviors describing the relative dynamics of the components. (A) A two-component loop network motif, which is similar to autoregulation but with two explicitly modeled genes instead of one. (B) Parameter space of the two regulatory strength parameters showing phase diagram for a loop composed of two activators (cross-activating) or two repressors (cross-inhibitory). Shading shows results of simulations (with an interval of 0.1 for both *γ* and *η* axes); blue curve is the analytically derived phase (bifurcation) boundary from Eqs 36. (C) Parameter space varying both strength parameters. Because the Hopf bifurcations depend only on the total delay and transient oscillations most prominent for equal delays, we show only the cases *γ*_1_ = *γ*_2_. Blue curves show analytically derived Hopf bifurcations. Black dashed curves are the bifurcation boundaries from (B). Except for the activator/repressor case, all these curves lie above the bistability boundary given by the black curve in (B), meaning oscillations are always transient. (D) Representative simulations for specific initial conditions showing all possible qualitative behaviors for a two-component loop with two activators, two repressors, or one activator and one repressor. Hill coefficients equal 2 for repressors, −2 for activators.

#### Two-component feedback loops shows additional behaviors not predicted from simple autoregulation

We can write down the governing equations for such a system, using the same normalizations as before (see Appendix S7), assuming for simplicity zero leakage and that *x* and *y* have the same degradation rates *β*. This is equivalent to the 3-step cascade Eq 8 with equal degradation rates and setting *Z* = *X* to form the loop.

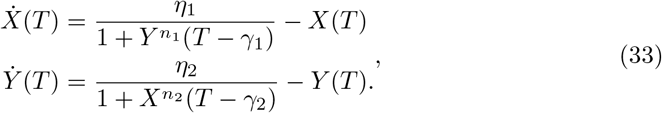

The fixed points are given by *η*_1_ = *X*(1 + *Y*^*n*_1_^), *η*_2_ = *Y*(1 + *X*^*n*_2_^) or (for two activators) *X* = *Y* = 0. Linearizing around these fixed points and assuming ansatz solutions of the form *δX*(*T*) = *A* exp(λ_1_ *T*), *δY*(*T*) = *B* exp(λ_2_ *T*), we find the set of characteristic equations

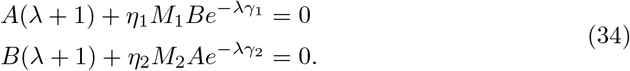

where λ_1_ = λ_2_ ≡ λ must hold for the ansatz to be true for all time. This can be rewritten in matrix form, with a matrix *J* times the vector (*A, B*)^⊤^. Diagonalizing *J* results in two characteristic equations (see Appendix S7)

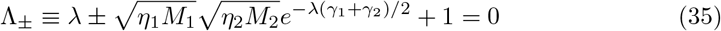

for two corresponding eigenmodes *v*_+_ = (1, *Re^iϕ^*)^⊤^ and *v*_−_ = (−1, *Re^iϕ^*)^⊤^, respectively, where *B/A* = *Re^iϕ^* is the ratio of *δX* to *δY* components in magnitude-phase notation (see Appendix S7). Overall, the eigenvectors indicate a phase difference between *X* and *Y* of *ϕ* for *v*_+_ and *ϕ* + *π* for *v*_−_. When the Λ_+_ equation is satisfied for Re λ = 0, *v*_+_ bifurcates and when the Λ_−_ equation is satisfied for Re λ = 0, *v*_−_ bifurcates.

#### Saddle-node bifurcations determine bistability modes

For saddle-node bifurcations, at which bistability begin, we set λ = 0. This can only be satisfied by the Λ_+_ equation if *n*_1_ < 0, *n*_2_ < 0 (two activators), and by the Λ_−_ equation if *n*_1_ > 0, *n*_2_ > 0 (two repressors). A feedback loop of an odd number of repressors cannot have a saddle-node bifurcation. As for linear cascades (see Fig 3), the overall autoregulation is repressive only if it contains an odd number of repressors. The bistability boundary for both cases is then given by (Appendix S7)

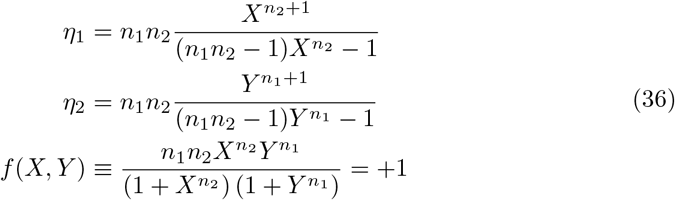

Here each *η* in the bifurcation boundary is determined by the steady state value of one of the variables, while the third equation is a monotonic mapping between *X* and *Y* that restricts the value of *η*_1_ relative to *η*_2_.

The phase difference *ϕ* = 0, thus we see that a two-activator loop bifurcates into fixed points where *X* and *Y* are both high or both low, because the components of *v*_+_ are of like sign, whereas a two-repressor loop (*v*_−_) bifurcates into fixed points where *X* is high and *Y* is low or vice versa. This can be seen clearly in the data (Fig 7B,D).

#### Hopf bifurcations determine transient oscillations in modes restricted by bistability

For Hopf bifurcations, at which oscillations begin, we set λ = *iω*. The boundaries are given for both eigenmodes by (Appendix S7)

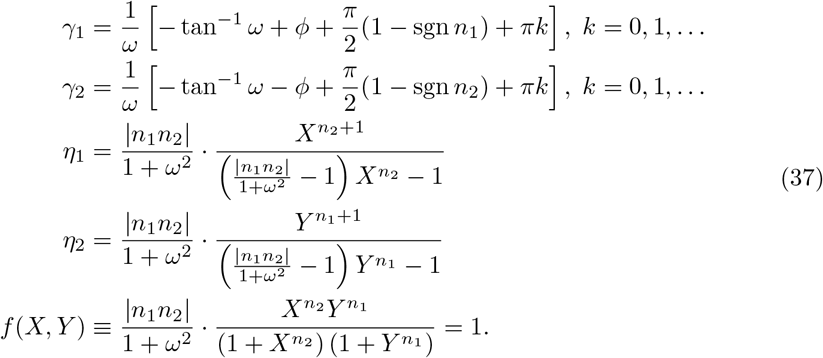

As before, *f* = 1 provides a monotonic mapping between *X* and *Y* that restricts *η*_1_ relative to *η*_2_. These curves always lie entirely above the bistability boundary found above. Thus, oscillations in *v*_+_ are not seen for two activators nor in *v*_−_ for two repressors, since those eigenmodes already have positive eigenvalues. The alternate eigenmode, however, does show oscillations in each case.

Thus, the primary Hopf bifurcation for two activators occurs for *v*_−_ (at *k* = 0) and the primary Hopf bifurcation for two repressors occurs for *v*_+_ (at *k* =1). Because bistability has occurred, however, there are distant stable fixed point, making these oscillations only transient. For one repressor and one activator, the primary bifurcation occurs at *k* = 0 for *v*_−_, with a phase of −*π*/2 if *n*_1_ > 0, *n*_2_ < 0 (*Y* leads *X*) and *π*/2 if *n*_1_ < 0, *n*_2_ > 0 (*Y* lags *X*) Because there is no bistability bifurcation, these are sustained oscillations.

Based on Eqs 37, we can see that the primary Hopf bifurcations are equivalent for differing values of the delays as long as the average delay is equal, because the oscillation frequency *ω* of the eigenmodes only depends (implicitly) on the average delay:

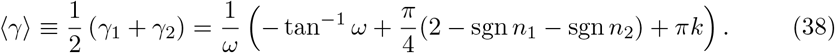

In particular, oscillations are possible for activator-repressor loops even when one delay is zero, as long as the sum of delays is greater than zero. On the other hand, the phase difference between the oscillations depends only on the difference in delays given the value *ω*:

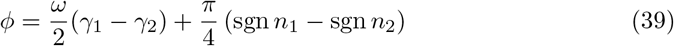

Together with the additional phase difference of *π* for *v*_−_, this implies that for equal delays, there are synchronous oscillations for two-repressor loops, anti-synchronous oscillations for two-activator loops, and *π*/2-shifted oscillations for activator-repressor loops. This phase relation also holds off the bifurcation boundary, where λ = *μ* + *iω*, as *ϕ* does not depend on *μ* (see Appendix S7). Note also that the overall phase in the expressions for *γ* are 0 for two repressors, *π* for two activators, and *π*/2 for activator-repressor loops.

Finally, one can also conclude from Eq 34 that (see Appendix S7)

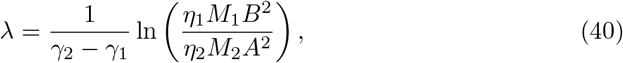

which holds for both the real and imaginary parts of λ. This implies that the oscillations grow most quickly and have the highest frequency when the delays are nearly equal (*γ*_1_ = *γ*_2_). This consequently makes the transients most noticeable for equal delays. All these results can be seen in the simulated data (Fig 7B,D).

#### Key results for feedback loops

1. Two-component feedback is very similar to direct autoregulation, but with new behaviors noticeable, particularly in transient behavior.
2. Transient synchronous oscillation are possible for double repressor loops with equal delays.
3. Transient anti-synchronous oscillation are possible for double activator loops with equal delays.
4. Sustained oscillations exist for repressor/activator loops.
5. Transient oscillations are most pronounced when the delays are equal, and more pronounced in double-repressor loops than in double-activator loops.
6. Oscillation requirements on regulatory strengths are determined by the average delay.
7. Phase in oscillations between components and time extent of transient oscillations are determined by the difference in delays.

### Motif VI: Double feedback

Thus far, every motif we have analyzed contained at most one feedback loop. Multiple feedback has been described to lead to interesting dynamics, such as excitablity [97] and chaos [82]. A full analysis of this motif is beyond the scope of this work, for reasons described below. In this section we focus on the possible chaotic dynamics that may arise when multiple (and most simply, double) feedback is present.

Chaos [41] is a major class of dynamics besides multi-stability and oscillations of potential importance to biology, characterized by oscillatory-like, unpredictable long-term behavior. It has been suggested that biological systems may have evolved to avoid chaos [98,99], but also that chaotic behavior can arise in disease states [33,100]. For example, Mackey and Glass (of no known relation to author) modeled irregular fluctuations in pathological breathing patterns and blood cell counts [33] using a delayed feedback model. Epilepsy has likewise been modeled as a chaotic disorder [101]. Much mathematical modeling has been done on gene networks that can give rise to chaos in both non-delayed [98,102,103] and delayed [82,104,105] models. We thus sought to determine under what conditions the delay-based models we studied in this paper can give rise to chaotic dynamics, if at all, and if so what a minimal chaos-producing motif may be.

#### Chaotic behaviors are prevented in monotonic regulation with linear or cyclic network topology

In fact, none of the motifs analyzed so far can ever produce chaotic dynamics. This is because cyclic network topology (no more than one loop) and monotonic regulation guarantee that a system of delay equations will not have any chaotic solutions [42]. All motifs we covered so far meet these criteria. Either non-monotonic feedback (as in the Mackey-Glass equation [33]) or non-cyclic topology has the possibility to produce chaotic dynamics.

The sum-of-Hills convention we used for expressing logic behavior excludes the two non-monotonic two-input logic gates, `XOR` and `XNOR` if the two inputs refer to different sources. However, this in not the case if a single source is used for both inputs (a feedforward loop). Specifically, the sum-of-Hills logic is non-monotonic if both the signs and the magnitudes of the Hill coefficients differ (see Appendix S8). If this feedforward loop is then connected in an overall feedback loop, it is equivalent to a two-arm autoregulation (Fig 8), from which we might expect chaotic solutions for some parameters. In fact, this double feedback can yield chaotic dynamics even with monotonic regulation (Fig 8)

**Fig 8.**
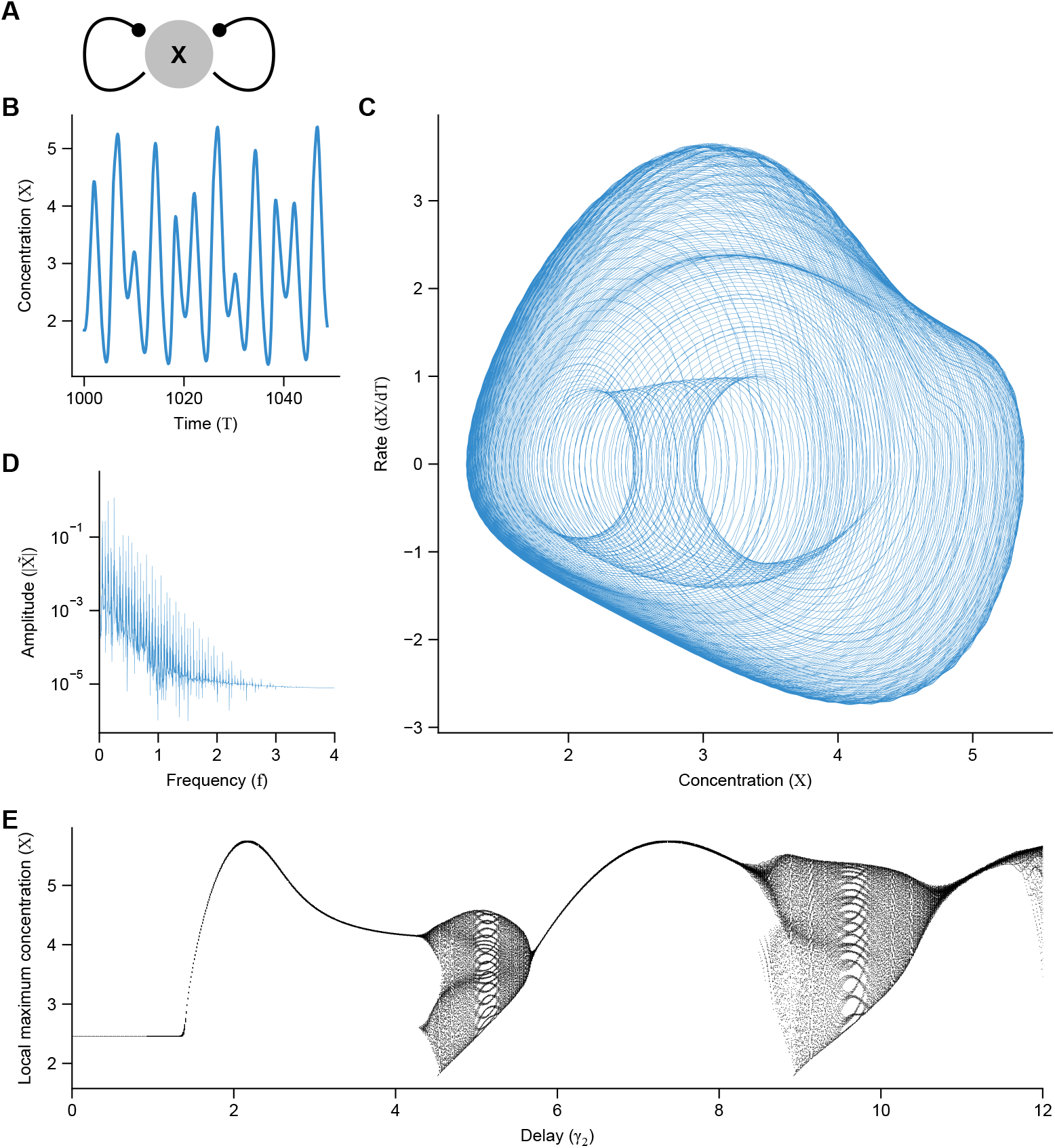
Double negative feedback induces chaotic behavior when the difference in delay times is significant. (A) Cartoon of double feedback. (B-E) focus on negative feedback, where chaos can occur. (B) Time trace of chaotic dynamics after initial transients. (C) Trace of dynamics in phase space, with the derivative on the vertical axis. While a simple oscillator would trace a loop (possibly with multiple sub-loops if the waveform is complicated), the chaotic dynamics appear to fill an entire region. (D) Fourier transform of chaotic dynamics show many peaks, indicating that there are no simple set of frequencies underlying the dynamics. (E) Bifurcation diagram for double negative feedback, with local maxima plotted. Simple oscillations intersperse chaotic regimes, where local maxima with a range of values are found. *η*_1_ = *η*_2_ = 16, *n*_1_ = 5, *n*_2_ = 2, and *γ*_1_ = 1. For (B-D), *γ*_2_ = 9.5.

The governing equation for such a double feedback motif is thus given by setting the output and both inputs of the logic equation (Eq 19) to *X* (and letting *K* = 1 for simplicity):

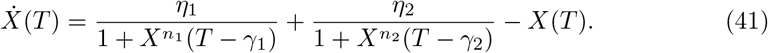

#### Chaotic behavior is possible for non-monotonic feedback with multiple feedback and disparate delays

From our simulation, we only found evidence for chaotic solutions if the delays in the two arms differ substantially. Simulation results for double negative feedback is shown in Fig 8 for *n*_1_ = 5, *n*_2_ = 2. Fig 8 illustrates several pieces of evidence that the dynamics presented are chaotic. First (Fig 8B), the time trace of *X*(*T*) does not appear to settle down to a fixed point, and oscillates but with local maxima that are variable in height. Second (Fig 8C), a phase-space trace of 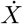 versus *X* occupies an entire region in phase space, rather than being confined to a 1-dimensional loop as would be expected for non-chaotic oscillation. Note that this implies chaos for many initial conditions, although we only used a single initial condition [41]. Third (Fig 8D), the Fourier transform of the dynamics exhibit a large number of peaks, indicating that the oscillations are not made up of a simple combination of frequencies.

A parameter space in which *γ*_2_ is varied relative to *γ*_1_ = 1 (Fig 8E) shows regions of chaotic behavior interspersed with simple oscillatory dynamics, akin to chaotic behavior in many systems [41]. We only explored the parameter space for *γ*_1_ = 1, but both delays appear to be important, not only their ratio or difference. In particular, there are no oscillations for both delays less than one. A full exploration of the two-delay parameter space is left to future work.

We also found chaotic dyanmics for dual positive/negative feedback (non-monotonic regulation) with *η*_1_ = 15, *η*_2_ = 1, *n*_1_ = 11, *n*_2_ = −3 and delays as in Fig 8 (see Fig. S4). Qualitatively, the time dynamics appear more pulsatile (Fig. S4B) than dual negative feedback. This makes sense with previous reports of dual positive/negative feedback demonstrating excitable behavior [97]. The attractor also has a somewhat different profile, notably containing a hole in the covered phase space (Fig. S4C), and the Fourier spectrum has a denser set of peaks (Fig. S4D). Given the consistent presence of an attracting (stable) fixed point (*X* = 0), dual positive feedback cannot support chaotic solutions [41].

Note that chaotic solutions are not expected even if the delays are different but the strengths and Hill coefficients are identical (i.e., *η*_1_ = *η*_2_ ≡ *η* and *n*_1_ = *n*_2_ ≡ *n*). One can see this by linearizing 41 and noting the *η* and *M* parameters can be pulled out in front of the delayed terms. Then assuming an ansatz solution of *A* exp(λ*T*) results in

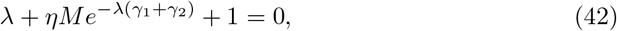

which is equivalent to the characteristic equation for autoregulation (Eq 13) with *γ* = *γ*_1_ + *γ*_2_. Thus the bifurcation boundaries will follow similar rules to autoregulation, ruling out chaotic solutions. The same holds for unequal strengths and Hill coefficients but equal delays, simply by replacing the coefficient *ηM* → *η*_1_*M*_1_ + *η*_2_*M*_2_. Since dual positive feedback likewise does not support chaotic solutions (see above), for chaotic solutions to occur via double feedback, there must be at least one negative feedback arm, the delays must differ, and either the cooperativities or strengths must differ. If biological systems evolved to avoid chaos [98, 99], it may be that these conditions are selected against, even if the double-feedback motif is not. The parameter space over which the fixed point is stable has been analyzed by Mahaffy and Zak [44], showing it to be in general formed by intersections of an infinite number of bifurcation curves.

For further exploration of double negative feedback and other delayed network motifs that exhibit chaotic dynamics, see Suzuki, *et al*. [82].

#### Key results for double feedback

1. Double feedback with a difference in delays between autoregulatory arms can lead to chaotic dynamics.2. For chaotic solutions, the two feedback arms must have different cooperativities and differ strongly in their delays.
2. Feedback need not be nonmonotonic for chaotic solution.
3. For chaotic solutions, at least one feedback arm must be repressive.
4. Simpler regulatory motifs cannot generate chaos.
5. A systematic analysis of the complete parameter space of double feedback is likely to be very complex [44] and is left for future work.

## Discussion

### The origin of delays and their effect on network motifs

The complex network of interactions in biological systems are often simplified for analysis by focusing on small network motifs that aim to capture their essential dynamics [2, 11, 16]. However, ignoring mechanistic detail can introduce effective delays into the interactions, and ignoring these delays in a mathematical model can lead to mischaracterization. In this work, we systematically studied delay differential equation (DDE) models of basic network motifs and compared them to the corresponding ordinary differential equation (ODE) models that lack explicit delays.

DDEs are well suited to characterizing the behavior of effective delays, because they incorporate explicit lags into dynamical governing equations without reference to the intermediate steps that give rise to them. In this work, we demonstrated mathematically that ODE-modeled, multi-step cascades and directly delayed, DDE-modeled cascades are nearly equivalent. Specifically, a cascade with no explicit delays can be approximated well as a single regulatory step with delay. Similarly, we showed that feedback cascades have very similar behavior to direct autoregulation. This establishes the familiarly mechanistic, ODE-modeled basis [30, 106, 107] for the more phenomenological, DDE-modeled regulation we focused on in this work.

Some details are lost in the DDE modeling simplification, such as the relative dynamics of components, or how the phenomenological, single-step parameters arise from the mechanistic steps. At the same time, however, DDE models show rich dynamics such as oscillations and chaos with fewer equations, variables, and parameters. Using simplified models based on DDEs we were able to provide complete phenomenological definitions of some of the most basic motifs, including cascades, logic, feedforward loops, autoregulation, multi-component feedback, and double feedback (Table 3). We further showed that the delays often determine key motif properties, such as the oscillation period in negative autoregulation, and pulse width and frequency cutoff in incoherent feedforward motifs. Interplay of multiple delays (e.g., their sums and differences) then play a similar role in multi-component feedback, for determining absolute and relative oscillation periods, and in multiple feedback, for determining chaotic dynamics.

**Table 3.**
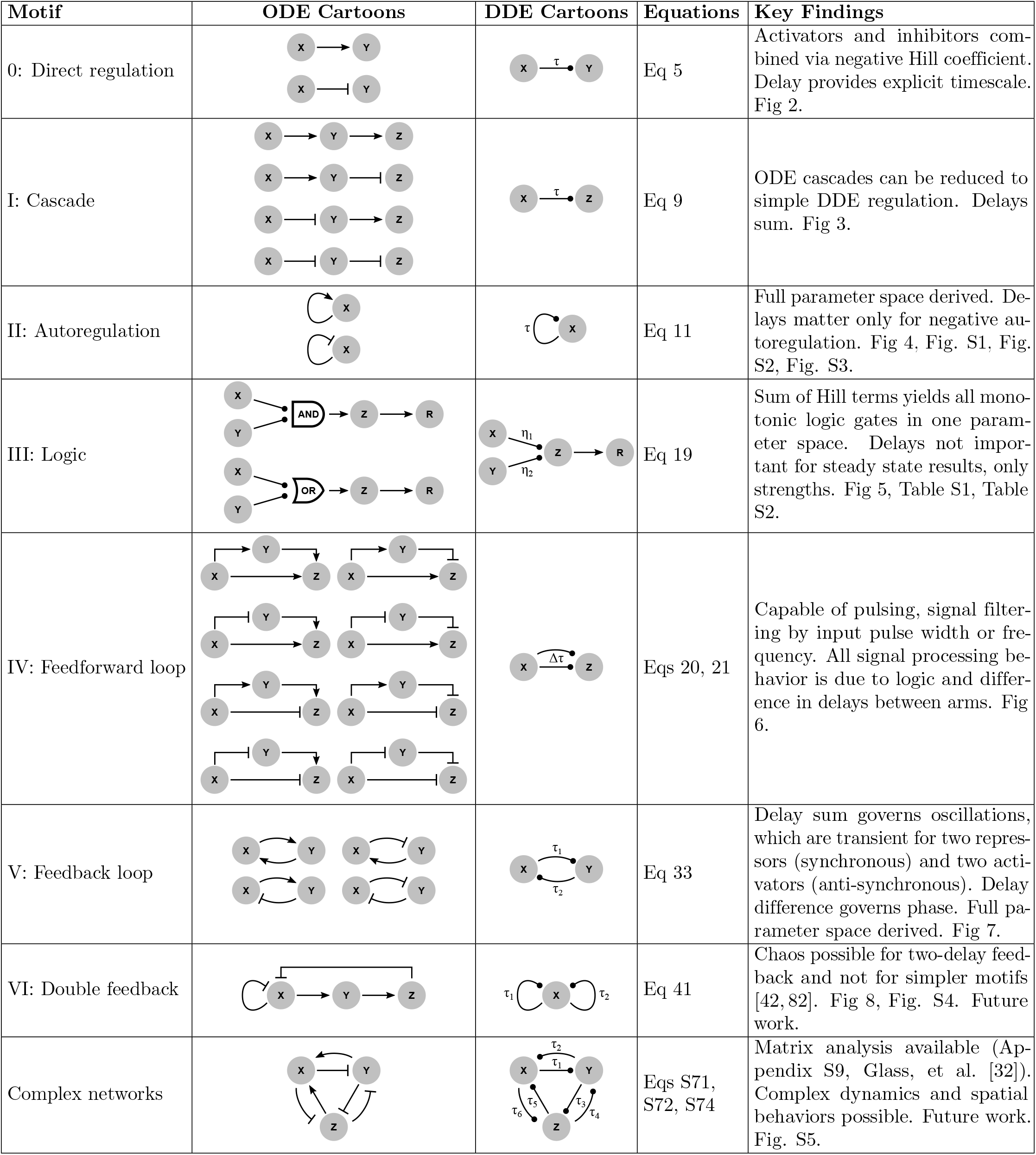
Summary of differences between ODE and DDE network regulation models and key findings. Each DDE cartoon provides a simplified, unified model for several ODE cartoons.

| Motif | ODE Cartoons | DDE Cartoons | Equations | Key Findings |
| --- | --- | --- | --- | --- |
| 0: Direct regulation |  |  | Eq 5 | Activators and inhibitors combined via negative Hill coefficient. Delay provides explicit timescale. Fig 2. |
| I: Cascade |  |  | Eq 9 | ODE cascades can be reduced to simple DDE regulation. Delays sum. Fig 3. |
| II: Autoregulation |  |  | Eq 11 | Full parameter space derived. Delays matter only for negative autoregulation. Fig 4, Fig. S1, Fig. S2, Fig. S3. |
| III: Logic |  |  | Eq 19 | Sum of Hill terms yields all monotonic logic gates in one parameter space. Delays not important for steady state results, only strengths. Fig 5, Table S1, Table S2. |
| IV: Feedforward loop |  |  | Eqs 20, 21 | Capable of pulsing, signal filtering by input pulse width or frequency. All signal processing behavior is due to logic and difference in delays between arms. Fig 6. |
| V: Feedback loop |  |  | Eq 33 | Delay sum governs oscillations, which are transient for two repressors (synchronous) and two activators (anti-synchronous). Delay difference governs phase. Full parameter space derived. Fig 7. |
| VI: Double feedback |  |  | Eq 41 | Chaos possible for two-delay feedback and not for simpler motifs [42, 82]. Fig 8, Fig. S4. Future work. |
| Complex networks |  |  | Eqs S71, S72, S74 | Matrix analysis available (Appendix S9, Glass, et al. [32]). Complex dynamics and spatial behaviors possible. Future work. Fig. S5. |

When simplifying networks using delay-based models, one must decide which and how many regulation steps to contract. If all possible cascades are “contracted” to delayed direct regulation and all looped cascades are contracted to delay direct autoregulation, then every network boils down to its topology (i.e., the number of loops) along with the strength, delay, cooperativity (including sign specifying activation or repression), and leakage associated with every remaining regulation. Changing the topology, however, fundamentally changes the motif. For example, we showed that replacing the cascade arm of a feedforward loop with a delayed, single-step regulation reproduces feedforward behavior (Fig 6). However, if an autoregulatory loop existed within this cascade (e.g., addition of self-activation to *Y* in Fig 6B), this would represent a fundamentally different network with likely different behavior. Importantly, if cascades are contracted without the addition of a delay, our results suggest that the model likely risks mischaracterizing the dynamics unless the individual steps are fast (i.e., production surpasses downstream half-maximal input much faster than it decays; in normalized terms, *η* ≫ 1). If one is interested in modeling behavior of specific components relative to one another, these specific components can be incorporated without contraction.

This ability to contract a network into a much smaller equivalent network (while preserving its topology) may aid in discovery of large-scale motifs that have important functions or that might otherwise be perceived as statistically insignificant. The motifs in this paper were originally discovered by scanning biological networks for subnetworks of *n* nodes that occur more frequently than expected by chance [11,16] (this in fact being the authors’ original definition of network motifs). However, this discovery method becomes increasingly difficult for *n* ≳ 5 due to the combinatorial scaling of the number of possible motifs [108]. Performing a similar search on a contracted network would be equivalent to searching the original network for larger motifs, and places a stronger emphasis on topology than on number of components involved (e.g., *X* → *Z* being equivalent to *X* → *Y* → *Z*). While this can be done without DDEs, the introduction of delays allows one to perform the contraction while maintaining key information about the original dynamics.

While our work has focused on a limited subset of network motif topologies, the same techniques can be applied to other more complex networks. Expanding on the eigenmode analysis we performed for multi-component feedback loops, one can describe the linear behavior of an arbitrary network near its fixed points by a matrix form of the characteristic equation (see Appendix S9, Fig. S5), 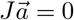 with 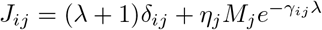. Diagonalizing *J* will yield a characteristic equation for each eigenmode, with a total number of eigenmodes corresponding to the number of components modeled in the network. Each mode can show bifurcations, yielding a potentially qualitatively rich set of dynamics. Not every mode will be equally important; those corresponding to the largest eigenvalues having the biggest effect on the results.

Consider for example, lateral inhibition patterning, in which each cell represses the production of a protein Delta in immediately neighboring cells [26, 32, 40]. In a one-dimensional tissue, the diagonal entries of *J* are λ + 1, the off-diagonal entries are filled, and all other entries zero. We have previously shown that this system gives rise to an alternating spatial pattern in which “errors” in the pattern corresponding to unwanted eigenmodes are removed by increasing delay [32] to a bifurcation boundary in which only one desired mode has positive λ. Thus, large-scale networks modeled using the presented framework can provide predictions regarding rich dynamics of relevance to complex biological phenomena with relative modeling simplicity.

### Limitations of the current analysis and future directions

We have endeavored in this work to present a unified, comprehensive view of biological network motifs and delays therein, to analyze the effect of delays on the most common motifs, and to ascertain where such delays may or may not be important for biological function. Nonetheless, several aspects of these explicit-delay models have been left out of our analysis for simplicity and present rich areas for future study. We detail here such limitations of the present analysis.

First, we did not incorporate any noise in our simulations, in order to clarify the analysis of the underlying equations. Biological systems are intrinsically noisy [109–111], and noise is known to affect dynamics of delay equations in sometimes unexpected ways [23, 105, 112, 113]. It would be important to include an analysis of noise in future work or in modeling specific biological systems. Reassuringly, our previous work on delays in lateral inhibition [32] showed no significant effects when the amplitude of the noise is significantly smaller than the strength of regulation. It therefore seems reasonable to expect that the main conclusions presented in this paper hold for similarly weak noise, although this has yet to be tested.

Second, all our equations have used a constant delay. It is likely that in many biological contexts delays will be variable in time [43,114]. Others have studied the effect of variable and stochastic delays [43,113,115] (another form of noise), which complicate the picture we portray here, and whose effects may provide new insight into mechanisms of biological regulation [113]. For example, where we saw that multi-component feedback loops have transient oscillations that are most prominent where all delays are equal, delay distributions may expand the parameter regimes in which such transients are significant if the delay distributions overlap.

Third, for all feedback or looped networks, we ran our simulations with initial conditions constant in time for all times smaller than zero (“constant histories”). Non-constant histories may show interesting effects on their own. For example, simulations of autorepression using Eq 11 show in-phase and anti-phase locking with constant histories regardless of the initial value, while sine-wave histories with randomized phase do not (see Fig. S6).

Fourth, we focused almost entirely in this work on single-variable Hill-function regulation and linear degradation. While this covers many biological networks [65], it cannot represent all scenarios [33, 71]. For example, we noted that the sum-of-Hills logic recapitulates many known behaviors of feedforward loops, but a single-term function is needed to demonstrate fold-change detection [71]. Furthermore, zeroth-order [116] and nonlinear [117] degradation kinetics certainly play many roles not covered here, nor did we consider cases where removal terms include a delay. Similarly, we did not include any diffusion terms or partial derivative important in many pattern formation networks [118].

Finally, with the exception of Eq 41, we have not studied any equations in this work that have both feedback and multiple delays. This type of equation applies to any situation in which a protein, e.g., regulates its own production through two mechanisms, with each mechanism having its own delay. Mahaffy, *et al*. analyzed a linear version of such a DDE [44], showing that whether or not a fixed point is stable depends on the generally non-trivial intersection of an infinite set of bifurcation curves. Analyzing the situation in the nonlinear biological context is likely to be very challenging [44, 46, 52].

On a related note, Eq 41 is the only equation we studied that shows any indications of chaotic behavior for any parameters. This is not surprising for the non-feedback motifs, where there can only be transient dynamics for constant inputs (at steady state all variables depend algebraically on the upstream regulators, and one can simply plug in the input values to find the output) [65]. In general, the Poincaré-Bendixson theorem [41] proves a lack of oscillations for ODEs of dimension (number of components) less than 2 and lack of chaos for dimension less than 3. However, this does not hold for DDEs [41,42], because trajectories can cross themselves, which is in turn due to the fact that the current state depends on a whole history. The system is thus formally infinite-dimensional, since an infinite number of initial condition values (the whole history) must be specified to predict future behavior [56].

Despite this added complexity, Mallet-Paret and Sell proved that cyclic networks with monotonic, bounded regulation functions like the systems focused on in our work cannot result in chaotic behavior [42]. The activator-like Mackey-Glass equation [33], in contrast, has a non-monotonic regulation *X*(*T* − 1)/(1 + *X^n^*(*T* − 1)), and is known to have chaotic solutions [33,104]. Likewise, more complicated networks with non-cyclic topologies such as double-negative feedback (e.g., Eq 41), where multiple delays affect a single component, can have chaotic solutions. Future work can more fully explore the parameter spaces and derive analytical boundaries for double feedback and other chaos-producing networks [82].

### Outlook

Our results, many derived entirely analytically, provide simple behavioral descriptions of common network motifs in which the key phenomenological parameters of delay, regulatory strength, cooperativity, and leakage have defined roles. We believe that the simplicity of our models, despite their added mathematical complexity stemming from the use of delay differential equations, can serve as a powerful framework for analyzing regulatory networks. For example, determining whether biological networks can oscillate may amount to calculating an effective delay and regulatory strength and comparing to Fig 4E and Fig. S1. Designing synthetic biological oscillators could benefit from a similar procedure, adjusting parameters through genetic engineering to cross the Hopf bifurcation thresholds [23, 31].

We have shown quantitatively when delays are necessary for behaviors such as oscillations. In contrast, our results also provide a strong prediction of where delays are not important. Delays do not affect steady states or logic of independent inputs in any network, Only a difference of delays between regulation arms is important in determining feedforward behavior. Only the sum of delays is important for determining onset of transient oscillations in feedback loops.

Finally, we believe that the methods presented here for modeling biological networks with explicit delays may help resolve fundamental biological questions. For example, a comparison of negative feedback against multiple feedback may help determine whether chaos affects the cell cycle [99], or whether biology evolves to avoid double feedback due to its capacity for unpredictable output. Delays are often significant in multicellular signaling [32, 40], and thus comparing effective delays against bistability boundaries can inform how realistic certain models of biological pattern formation may be [24,34,40,118–121]. Exploring “contracted” networks in which linear and looped cascades are replaced with single-step delayed regulation may help uncover new functional motifs in biological networks that are larger and more complex than those known today [108]. We hope that our work on the detailed mechanics of delay models in biological contexts will be helpful in studying and engineering a wide variety of phenomena, including transcription factor networks [11,26,61], cell cycles [99] other biological clocks [22, 27, 35], and pattern formation [32, 122].

## Materials and methods

Analytics were in general performed by hand, and checked for validity using Mathematica. Numerical simulations were run in Matlab using the `dde23` delay differential equation solver for DDEs and `ode45` for ODEs. For autoregulation phase plots, simulations were run with 100 constant-history initial conditions spread logarithmically between 10^−4^ and 2*η* and run from *T* = 0 to *T* = 100(*γ* + 1). Solutions were considered stable if for all 100 simulations the maximum absolute value of the discrete derivative in the last three-quarters of the simulation time was less than 0.1. Stable solutions were sub-categorized as bistable if a histogram of final values over all 100 solutions had more than 1 peak. Solutions were considered oscillatory if the average Fourier transform of the last three-quarters of the simulation time for all 100 solutions had more than zero peaks with amplitude (square root of power) greater than 100. Solutions were considered spiral if this oscillation condition held for the first one-quarter of the simulation time only. For two-component loops, initial conditions were used that ranged between 0 and max(*η*_1_, *η*_2_), for equal *X* and *Y* and for apposing *X* and *Y*. Bistability was determined as for autoregulation, and a cutoff of 0.05 was used to determine “low” values. All simulation histories were constant except where indicated in Fig. S6. Specific parameter values and simulation details are given in the figures and/or made explicit in the MATLAB code in Data S1.

## Data availability

All raw data and code is included in the supplementary material (Data S1).

## Acknowledgments

We would like to thank A. Choksi, J. Prine, M. Aktas, Y. Yang, A. Mayo, U. Alon, and the Riedel-Kruse lab for their feedback and discussions. This work was supported in part by the Davies Family Engineering Fellowship, the Bio-X Bowes Graduate Student Fellowship, and the NSERC PGS fellowship. D.S.G is currently a member of the Zuckerman Postdoctoral Scholars Program.

## Author contributions

D.S.G., X.J., and I.H.R.K. all contributed to conception, analysis, and writing of this manuscript.

