## Supplementary Material for "Nonlinear delay differential equations and their application to modeling biological network motifs"

### Supporting information

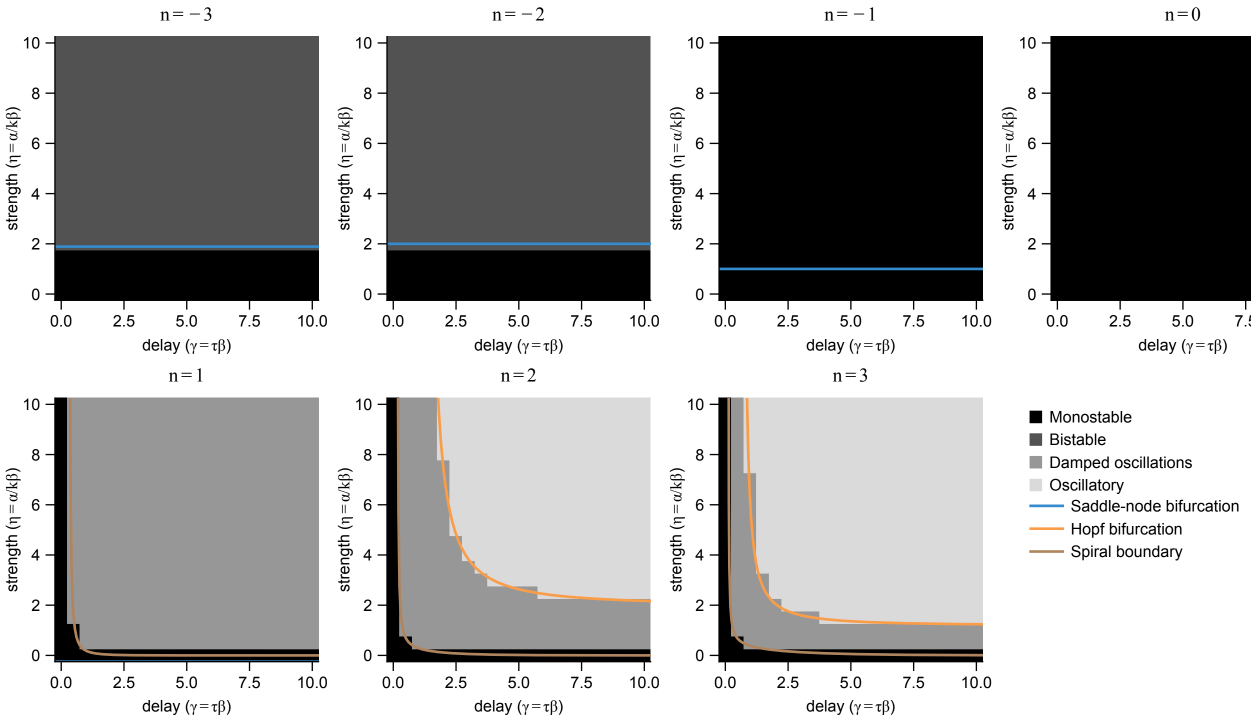


**Fig. S1 Phase diagrams for autoregulation with delay for a range of cooperativity.** As in Fig 4E, Shading represents simulation results and curves represent analytically derived phase boundaries (with an interval of 0.5 for both $\gamma$ and $\eta$ axes). Note that for $n = -1$, there is a degenerate case of the saddle-node bifurcation in which the system is monostable on both sides of the $\eta = 1$ boundary. For $\eta \leq 1$, the stable fixed point is located at the origin, and for $\eta > 1$, the location of the stable becomes non-zero ($X^* > 0$). Also note that the cases $n = \pm 2$ match Fig 4E.

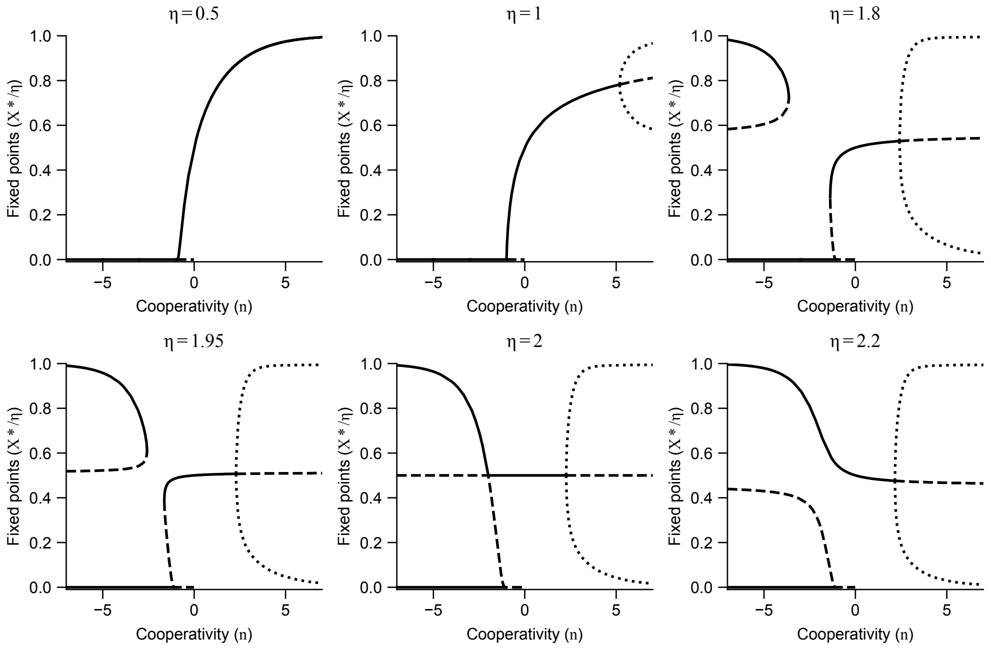


**Fig. S2 Bifurcation diagrams for autoregulation with delay.** These curves represent an alternative way to represent the results in Figs 4, Fig. S1, with more quantitative detail on how behavior changes when parameters cross the bifurcation curves shown in Fig 4E. Specifically, each plot is taken at a different $\eta$, while cooperativity $n$ is varied and the delay is set at $\gamma = 5$ for simplicity. Only the point at which oscillations begin would change with $\gamma$. Solid curves represent stable fixed point values, dashed lines unstable fixed points, and dotted lines the minimum/maximum values reached in oscillations. Solid and dashed lines were drawn based on the analytical curves; dotted lines were taken from simulation results. Note that there are only 2 fixed points for $-1 \leq n < 0$ due to the lack of inflection point in the Hill function.

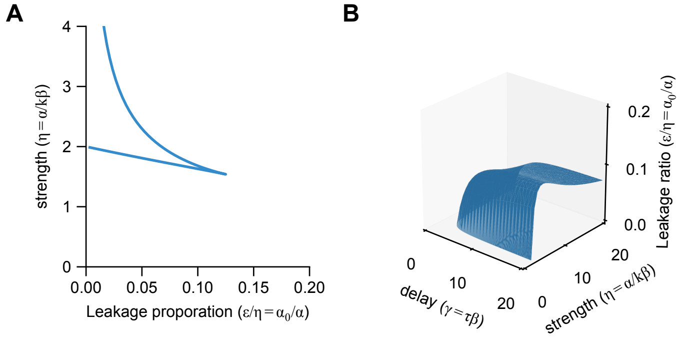


**Fig. S3 Phase boundaries for autoregulation with delay and leakage.** Leakage (i.e., basal expression) limits the parameter space in which bistability and oscillations are achieved. (A) Bistability boundary (saddle-node bifurcation, Eqs S22) as a function of fractional leakage ($\epsilon/\eta$) and regulation strength ($\eta$). Bistability is achieved to the left of the curve, and does not depend on delay. $n = -2$ (B) Oscillation boundary (Hopf bifurcation, Eqs S27) as a function of fractional leakage ($\epsilon/\eta$), strength ($\eta$), and delay ($\gamma$). Oscillations are achieved below the surface. $n = 2$. We show fractional leakage ($\epsilon/\eta$) rather than absolute leakage ($\epsilon$) to accentuate the upper bound on leakage, which does not occur for (B) in absolute terms.

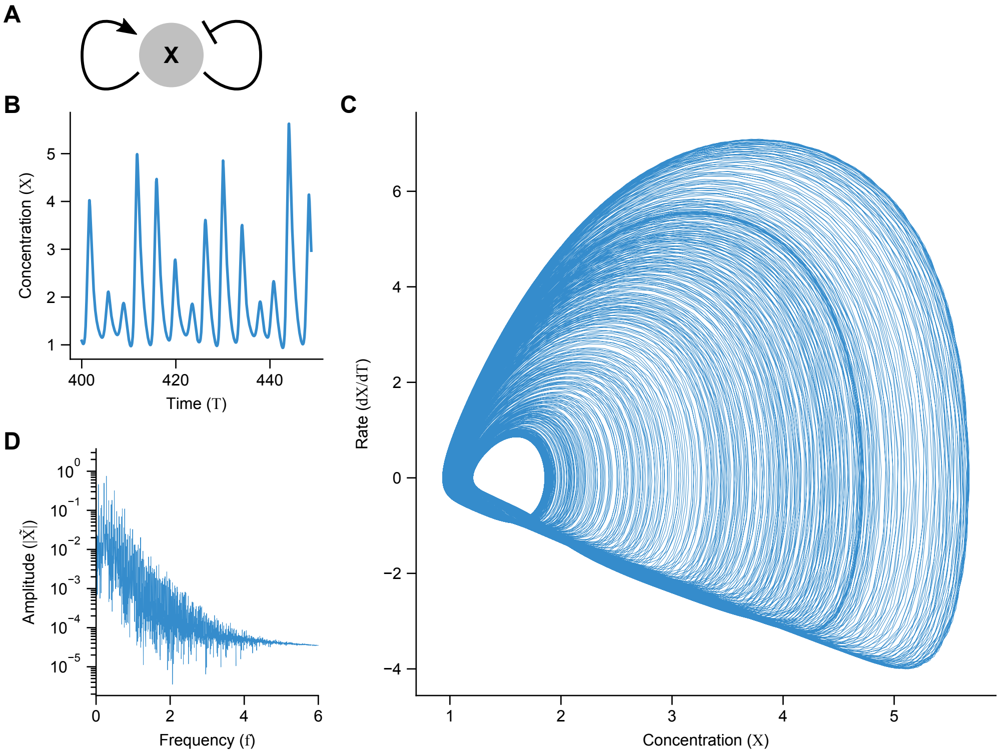


**Fig. S4 Positive/negative dual feedback induces chaotic behavior when the difference in delay times is significant.** (A) The double feedback motif, in which two regulation arms feed back directly, each with its own explicit delay. (B) Time trace of chaotic dynamics after initial transients. (C) Trace of dynamics in phase space, with the derivative on the vertical axis. While a simple oscillator would trace a loop (possibly with multiple sub-loops if the waveform is complicated), the chaotic dynamics appear to fill an entire region. (D) Fourier transform of chaotic dynamics show many peaks, indicating that there are no simple set of frequencies underlying the dynamics. $\eta_1 = 15$, $\eta_2 = 1$, $n_1 = 11$, $n_2 = -3$, $\gamma_1 = 1$, and $\gamma_2 = 9.5$.

**A**

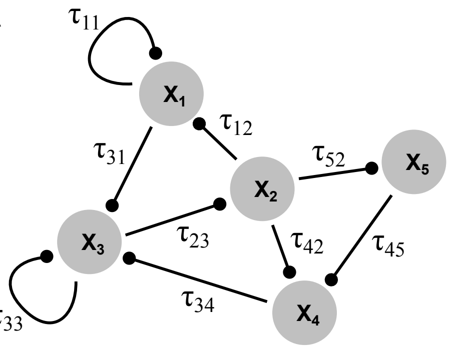


**B**

$$
J \quad = \quad (\lambda + 1)\ \mathrm{I}_5 \quad + \quad
\begin{pmatrix}
\eta_{11} M_1 e^{-\lambda \gamma_{11}} & \eta_{12} M_2 e^{-\lambda \gamma_{12}} & 0 & 0 & 0 \\
0 & 0 & \eta_{23} M_3 e^{-\lambda \gamma_{23}} & 0 & 0 \\
\eta_{31} M_1 e^{-\lambda \gamma_{31}} & 0 & \eta_{33} M_3 e^{-\lambda \gamma_{33}} & \eta_{34} M_4 e^{-\lambda \gamma_{34}} & 0 \\
0 & \eta_{42} M_2 e^{-\lambda \gamma_{42}} & 0 & 0 & \eta_{45} M_5 e^{-\lambda \gamma_{45}} \\
0 & \eta_{52} M_2 e^{-\lambda \gamma_{52}} & 0 & 0 & 0
\end{pmatrix}
$$

**Fig. S5 Decomposition of a complex network.** (A) An example of a complex network with many delays. (B) The interaction matrix $J$, defined in Eq S72, representing the network in (A). I refers to a $5 \times 5$ identity matrix. The diagonalization of $J$ leads to five eigenvalues $\Lambda_1$–$\Lambda_5$, each of which can be independently set to zero to describe the dynamics of five independent eigenmodes.

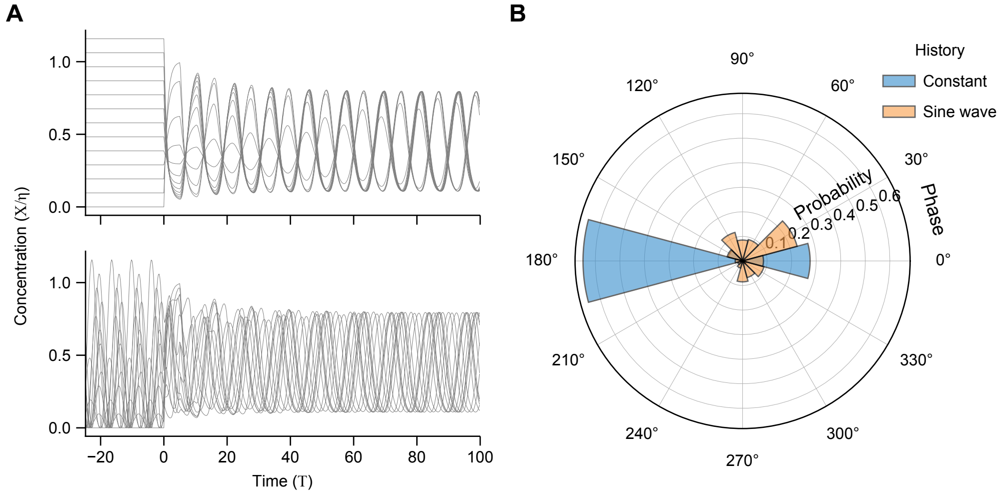


**Fig. S6 Non-constant histories may show non-trivial effects on oscillations.** (A) Time traces of autorepression with constant histories (top) and randomized-phase sine-wave functions (bottom). Amplitudes of the sine waves and values of the constant histories are spread linearly between 0 and $1.2\eta$. (B) Histograms of phases relative to the first constant-history oscillation, calculated by finding the maximum of the cross-correlation between the data and reference curve. $\eta = 4$, $\gamma = 5$, and $n = 1$.

| $n_1$ | $n_2$ | $X$ | $Y$ | $Z^*$ | non-`TRUE` | $\eta_{Z1} < 1$, $\eta_{Z2} < 1$, $\eta_{Z1}+\eta_{Z2} < 1$ | | $\eta_{Z1} < 1$, $\eta_{Z2} > 1$ | | $\eta_{Z1} > 1$, $\eta_{Z2} < 1$ | | $\eta_{Z1} < 1$, $\eta_{Z2} < 1$, $\eta_{Z1}+\eta_{Z2} > 1$ | | $\eta_{Z1} > 1$, $\eta_{Z2} > 1$ | |
|---|---|---|---|---|---|---|---|---|---|---|---|---|---|---|---|
| $> 0$ | $> 0$ | $\ll 1$ | $\ll 1$ | $\eta_{Z1} + \eta_{Z2}$ | $\eta_{Z1} + \eta_{Z2}$ | $< 1$ | `FALSE` | $> 1$ | `NOT` $Y$ | $> 1$ | `NOT` $X$ | $> 1$ | $X$ `NOR` $Y$ | $> 1$ | $X$ `NAND` $Y$ |
| $> 0$ | $> 0$ | $\ll 1$ | $\gg 1$ | $\eta_{Z1} + \eta_{Z2}/\eta_Y^{n_2}$ | $\eta_{Z1}$ | $< 1$ | | $< 1$ | | $> 1$ | | $< 1$ | | $> 1$ | |
| $> 0$ | $> 0$ | $\gg 1$ | $\ll 1$ | $\eta_{Z1}/\eta_X^{n_1} + \eta_{Z2}$ | $\eta_{Z2}$ | $< 1$ | | $> 1$ | | $< 1$ | | $< 1$ | | $> 1$ | |
| $> 0$ | $> 0$ | $\gg 1$ | $\gg 1$ | $\eta_{Z1}/\eta_X^{n_1} + \eta_{Z2}/\eta_Y^{n_2}$ | 0 | $< 1$ | | $< 1$ | | $< 1$ | | $< 1$ | | $< 1$ | |
| $> 0$ | $< 0$ | $\ll 1$ | $\ll 1$ | $\eta_{Z1} + \eta_{Z2}\eta_Y^{\lvert n_2\rvert}$ | $\eta_{Z1}$ | $< 1$ | `FALSE` | $< 1$ | $Y$ | $> 1$ | `NOT` $X$ | $< 1$ | $Y$ `NIMPLY` $X$ | $> 1$ | $X$ `IMPLY` $Y$ |
| $> 0$ | $< 0$ | $\ll 1$ | $\gg 1$ | $\eta_{Z1} + \eta_{Z2}$ | $\eta_{Z1} + \eta_{Z2}$ | $< 1$ | | $> 1$ | | $> 1$ | | $> 1$ | | $> 1$ | |
| $> 0$ | $< 0$ | $\gg 1$ | $\ll 1$ | $\eta_{Z1}/\eta_X^{n_1} + \eta_{Z2}\eta_Y^{\lvert n_2\rvert}$ | 0 | $< 1$ | | $< 1$ | | $< 1$ | | $< 1$ | | $< 1$ | |
| $> 0$ | $< 0$ | $\gg 1$ | $\gg 1$ | $\eta_{Z1}/\eta_X^{n_1} + \eta_{Z2}$ | $\eta_{Z2}$ | $< 1$ | | $> 1$ | | $< 1$ | | $< 1$ | | $> 1$ | |
| $< 0$ | $> 0$ | $\ll 1$ | $\ll 1$ | $\eta_{Z1}\eta_X^{\lvert n_1\rvert} + \eta_{Z2}$ | $\eta_{Z2}$ | $< 1$ | `FALSE` | $> 1$ | `NOT` $Y$ | $< 1$ | $X$ | $< 1$ | $X$ `NIMPLY` $Y$ | $> 1$ | $Y$ `IMPLY` $X$ |
| $< 0$ | $> 0$ | $\ll 1$ | $\gg 1$ | $\eta_{Z1}\eta_X^{\lvert n_1\rvert} + \eta_{Z2}/\eta_Y^{n_2}$ | 0 | $< 1$ | | $< 1$ | | $< 1$ | | $< 1$ | | $< 1$ | |
| $< 0$ | $> 0$ | $\gg 1$ | $\ll 1$ | $\eta_{Z1} + \eta_{Z2}$ | $\eta_{Z1} + \eta_{Z2}$ | $< 1$ | | $> 1$ | | $> 1$ | | $> 1$ | | $> 1$ | |
| $< 0$ | $> 0$ | $\gg 1$ | $\gg 1$ | $\eta_{Z1} + \eta_{Z2}/\eta_Y^{n_2}$ | $\eta_{Z1}$ | $< 1$ | | $< 1$ | | $> 1$ | | $< 1$ | | $> 1$ | |
| $< 0$ | $< 0$ | $\ll 1$ | $\ll 1$ | $\eta_{Z1}\eta_X^{\lvert n_1\rvert} + \eta_{Z2}\eta_Y^{\lvert n_2\rvert}$ | 0 | $< 1$ | `FALSE` | $< 1$ | $Y$ | $< 1$ | $X$ | $< 1$ | $X$ `AND` $Y$ | $< 1$ | $X$ `OR` $Y$ |
| $< 0$ | $< 0$ | $\ll 1$ | $\gg 1$ | $\eta_{Z1}\eta_X^{\lvert n_1\rvert} + \eta_{Z2}$ | $\eta_{Z2}$ | $< 1$ | | $> 1$ | | $< 1$ | | $< 1$ | | $> 1$ | |
| $< 0$ | $< 0$ | $\gg 1$ | $\ll 1$ | $\eta_{Z1} + \eta_{Z2}\eta_Y^{\lvert n_2\rvert}$ | $\eta_{Z1}$ | $< 1$ | | $< 1$ | | $> 1$ | | $< 1$ | | $> 1$ | |
| $< 0$ | $< 0$ | $\gg 1$ | $\gg 1$ | $\eta_{Z1} + \eta_{Z2}$ | $\eta_{Z1} + \eta_{Z2}$ | $< 1$ | | $> 1$ | | $> 1$ | | $> 1$ | | $> 1$ | |

**Table S1 Extended logic table.** Full set of data corresponding to Table 1 and Fig 5B. For each set of parameters the fixed-point value $Z^*$ is given based on Eq 19. The formulae in the "non-`TRUE`" column are given under the assumptions that $\eta_{Z1} \ll \max(\eta_X^{\lvert n_1\rvert})$ and $\eta_{Z2} \ll \max(\eta_Y^{\lvert n_2\rvert})$, outside of which $Z^*$ is always $> 1$ regardless of the values of $X$ and $Y$, meaning the output is logical TRUE. The last five columns give $Z^*$ in the parameter regions specified by the column headers, as well as the corresponding logic functions. The function X IMPLY Y = NOT(X) OR Y, and the function X NIMPLY Y = X AND NOT(Y). Note that the output at $Z$ is fuzzy logic, in that the value $Z^*$ can be close to 1 (the halfway point between FALSE and TRUE). The activation of a downstream reporter $R$ by $Z$ with high cooperativity digitizes this signal into non-fuzzy logic, essentially turning "$> 1$" and "$< 1$" in the last two columns of the table into "$\ll 1$" and "$\gg 1$", respectively.

| | | | | | | AND-type | | | OR-type | | |
|---|---|---|---|---|---|---|---|---|---|---|---|
| FALSE | | | TRUE | | | X AND Y | | | X OR Y | | |
| X | Y | Z | X | Y | Z | X | Y | Z | X | Y | Z |
| 0 | 0 | 0 | 0 | 0 | 1 | 0 | 0 | 0 | 0 | 0 | 0 |
| 0 | 1 | 0 | 0 | 1 | 1 | 0 | 1 | 0 | 0 | 1 | 1 |
| 1 | 0 | 0 | 1 | 0 | 1 | 1 | 0 | 0 | 1 | 0 | 1 |
| 1 | 1 | 0 | 1 | 1 | 1 | 1 | 1 | 1 | 1 | 1 | 1 |

| X | | | Y | | | X NOR Y | | | X NAND Y | | |
|---|---|---|---|---|---|---|---|---|---|---|---|
| X | Y | Z | X | Y | Z | X | Y | Z | X | Y | Z |
| 0 | 0 | 0 | 0 | 0 | 0 | 0 | 0 | 1 | 0 | 0 | 1 |
| 0 | 1 | 0 | 0 | 1 | 1 | 0 | 1 | 0 | 0 | 1 | 1 |
| 1 | 0 | 1 | 1 | 0 | 0 | 1 | 0 | 0 | 1 | 0 | 1 |
| 1 | 1 | 1 | 1 | 1 | 1 | 1 | 1 | 0 | 1 | 1 | 0 |

| NOT X | | | NOT Y | | | X NIMPLY Y | | | X IMPLY Y | | |
|---|---|---|---|---|---|---|---|---|---|---|---|
| X | Y | Z | X | Y | Z | X | Y | Z | X | Y | Z |
| 0 | 0 | 1 | 0 | 0 | 1 | 0 | 0 | 0 | 0 | 0 | 1 |
| 0 | 1 | 1 | 0 | 1 | 0 | 0 | 1 | 0 | 0 | 1 | 1 |
| 1 | 0 | 0 | 1 | 0 | 1 | 1 | 0 | 1 | 1 | 0 | 0 |
| 1 | 1 | 0 | 1 | 1 | 0 | 1 | 1 | 0 | 1 | 1 | 1 |

| X XOR X | | | X XNOR Y | | | Y NIMPLY X | | | Y IMPLY X | | |
|---|---|---|---|---|---|---|---|---|---|---|---|
| X | Y | Z | X | Y | Z | X | Y | Z | X | Y | Z |
| 0 | 0 | 0 | 0 | 0 | 1 | 0 | 0 | 0 | 0 | 0 | 1 |
| 0 | 1 | 1 | 0 | 1 | 0 | 0 | 1 | 1 | 0 | 1 | 0 |
| 1 | 0 | 1 | 1 | 0 | 0 | 1 | 0 | 0 | 1 | 0 | 1 |
| 1 | 1 | 0 | 1 | 1 | 1 | 1 | 1 | 0 | 1 | 1 | 1 |

**Table S2 Truth tables.** Truth table representations of all 16 2-input logic functions are given here for reference. Each function is given inputs X and Y and outputs Z. The function names are color-coded as responding effectively to 0 inputs (white), 1 input (light gray), 2 inputs (gray), and non-monotonically to 2 inputs (dark gray). Non-monotonic for logic functions means that $Z(X, Y = 1) > Z(X, Y = 0)$ and $Z(1, Y) > Z(0, Y)$. Note that the right two columns are the "AND-type" and "OR-type" logic gates. For example, X NOR Y = NOT X AND NOT Y, and also Y IMPLY X = X OR NOT Y.

**Data S1 Matlab simulation code and data.** Zip file containing all Matlab simulation code and binary-data results.

**Appendix S1 Full set of derivations related to the cascade motif.** Here we include a detailed derivation for the cascade-section equations.

##### Nondimensionalization

The dimensionful governing equations for a cascade in which $X$ regulates $Y$ which in turn regulates $Z$ are given by:

$$\begin{aligned} \dot{x}(t) &= \alpha_x - \beta_x x(t) \\ \dot{y}(t) &= \frac{\alpha_y k_x^{n_x}}{k_x^{n_x} + x^{n_x}(t)} - \beta_y y(t) \\ \dot{z}(t) &= \frac{\alpha_z k_y^{n_y}}{k_y^{n_y} + y^{n_y}(t)} - \beta_z z(t). \end{aligned} \tag{S1}$$

These can be nondimensionalized by setting $X = x/k_x$, $Y = y/k_y$, $Z = z/k_z$, $T = t\beta_x$ and assigning $\eta_x = \alpha_x/k_x\beta_x$, $\eta_y = \alpha_y/k_y\beta_x$, $\eta_z = \alpha_z/k_z\beta_x$) with $k_z$ being the half-maximal concentration of $z$ in its downstream use. This yields Eqs 8 from the main text.

Following the arguments in the main text for a delayed pseudo-steady state, we solve for $Y$ by setting $\dot{Y}(T) \approx 0$. We then plug this into the equation for $\dot{Z}(T)$ but with $X(T)$ replaced by $X(T - \gamma)$. This delay $\gamma = \beta_x/\beta_y$ reflects the time that $Y$ takes to change. This yields a compound Hill-of-Hill regulation term, given as:

$$\dot{Z}(T) = \frac{\eta_Z}{1 + \left(\frac{\eta_Y \gamma}{1 + X^{n_X}(T-\gamma)}\right)^{n_Y}(T)} - \frac{\beta_z}{\beta_x} Z(T). \tag{S2}$$

We next compare the composite regulation term to a regular Hill term with leakage, $\alpha_0 + \alpha k^h/(k^h + X^h)$ (main text Eq 9). By equating the values of the two terms at $X = 0$, $X = 1$, $X \to \infty$ and the derivatives at $X = 1$, we have four equations with four unknowns ($\alpha_0$, $\alpha$, $k$, and $m$). Solving yields the assignments of these four parameters in terms of those from the original set of equations, given in the main text (Eq 10).

Additionally, the sign of the composite regulation's derivative is $\operatorname{sgn} n_X n_Y$. The sign of the regular Hill term is $-\operatorname{sgn} h$. Neither are dependent on $X$, so both terms are monotonic, supporting the view of approximating the composite Hill-of-Hill term as a single Hill function.

#### Appendix S2 Full set of derivations related to the autoregulation motif.

Here we include a detailed derivation for the autoregulation-section equations other than leakage (see Appendix S3) and damped oscillations (see Appendix S4).

##### Nondimensionalization

The dimensionful governing equation for autoregulation is given by:

$$\dot{x}(t) = \alpha_0 + \frac{\alpha k^n}{k^n + x^n(t-\tau)} - \beta x(t). \tag{S3}$$

This equation can be non-dimensionalized by normalizing the protein concentration by the half-maximal input and the time by the degradation time scale $1/\beta$. Explicitly, this means performing the change of variables $X = x/k, T = t\beta$. This simplifies to the normalized, non-dimensional governing equation given in the main text(Eq 11). The normalized delay $\gamma = \tau\beta$ represents the ratio of delay to degradation times. The normalized regulation strength $\eta = \alpha/k\beta$ and leakage $\epsilon = \alpha_0/k\beta$ represent production of normalized concentration per degradation time.

##### Fixed points

The fixed points of Eq 11 are found by setting $\dot{X} = 0$ and $X(T) = X(T - \gamma) \equiv X^*$, we find (with $\epsilon = 0$ for simplicity) that all fixed points $X^*$ are given by Eq 12 or $X^* = 0$.

#### Linearization

The governing equation (Eq 11) can be linearized around the fixed points $X^*$, by setting $\delta X(T) = X(T) - X^* \ll 1$ and expanding to first order:

$$\delta\dot{X}(T) + \eta M(X^*)\delta X(T-\gamma) + \delta X(T) = 0, \tag{S4}$$

with the constant function of the fixed point value $M(X^*)$ given as in the main text (Eq 14). This value $M$ is positive for the repressor case and negative for the activator case, since all terms in $M$ are positive due to $X^* > 0$ except the prefactor $n$.

Plugging an assumed solution $\delta X(T) = A\exp(\lambda T)$ into Eq S4 yields the transcendental characteristic equation (Eq 13 from the main text) for the eigenvalues $\lambda$.

#### Bifurcations

For the saddle-node case ($\lambda = 0$), Eq 13 reduces to:

$$\eta M = -1. \tag{S5}$$

Since $\eta > 0$ always, it must be the case that $M < 0$ (i.e., $n < 0$), meaning that a saddle-node bifurcation can only occur for autoactivation. Taken together, Eqs 14 and 13 imply that the for the fixed point $X^* = 0$, $M = 0$ as long as $-n > 1$, and thus $\lambda = -1$, making this fixed point unconditionally stable. For $-n < 1$, the derivative near $X = 0$ is positive, making it unstable. In general, there are other fixed points as long as $-n > 1$, because then there is an inflection point in the regulation curve. For these other fixed points, we insert the value of $\eta$ from Eq 12 and the value of $M$ from Eq 14 into Eq S5 to yield

$$n\frac{X^{*n}}{1+X^{*n}} = -1. \tag{S6}$$

Becuase the fraction involving $X$ is always between 0 and 1, we must further require that $-n > 1$ to balance the equation. Solving for the fixed point $X^*$ yields

$$X^* = (-n-1)^{-1/n}. \tag{S7}$$

Finally, plugging back into Eq 12 yields the bifurcation line for $\eta$ given by Eq 15 in the main text.

For the Hopf case ($\lambda = i\omega$), Eq 13 can be separated into real and imaginary parts, yielding:

$$\begin{aligned} \eta M\cos(\gamma\omega) &= -1 \\ \eta M\sin(\gamma\omega) &= \omega \end{aligned}. \tag{S8}$$

Solving for $\eta M$ in the first equation and plugging back into the second, then solving for $\gamma$ yields:

$$\gamma = \frac{1}{\omega}\left(-\tan^{-1}\omega + \pi k\right), \qquad k = 0, 1, \ldots \tag{S9}$$

which are a set of bifurcation boundaries on $\gamma$, with smaller $k$ bifurcating first. Plugging this solution for $\gamma$ back into the first of Eqs S8, and substituting in the values for $\eta$ from Eq 12 and $M$ from 14 yields

$$(-1)^{k+1}n \cdot \frac{1}{\sqrt{1+\omega^2}} \cdot \frac{X^{*n}}{1+X^{*n}} = 1 \qquad . \tag{S10}$$

For the signs on each side of the equation to hold, we must have $k$ even for $n < 0$ and $k$ odd for $n > 0$; in other words, we have $(-1)^{k+1}n = |n|$. Since both of the fractions are always less than 1, we must also have than $|n| > 1$ in order to balance the equation. Solving this for $X^*$, we find

$$X^* = \left( \frac{|n|}{\sqrt{1+\omega^2}} - 1 \right)^{-1/n} \quad . \tag{S11}$$

Finally, plugging back into Eqs S8, we get the Hopf boundary parameterized by $\omega$ as found in the main text (Eq 16).

#### Estimation of oscillatory period

To estimate the oscillatory period of oscillations for $n > 0$ beyond the Hopf boundary, we follow the method from Julian Lewis [26].

We first rewrite the governing equation (Eq 11, with $\epsilon = 0$) with the degradation term on the left:

$$\dot{X}(T) + X(T) = \frac{\eta}{1 + X^n(T-\gamma)}. \tag{S12}$$

The right side of this equation is a monotonically decreasing function, so it has a maximum when the left side has a minimum, and vice versa.

When $\gamma \gg 1$, the degradation-driven equilibration occurs much faster than the delay time. In this case, the system shifts between psuedo-steady states (i.e., $\dot{X}(T) \approx 0$), and switches between these steady states approximately every $\gamma$ time steps. Thus the system approximates a cascade $X(T-2\gamma) \to X(T-\gamma) \to X(T) \to \cdots$. The pseudo-steady states are then given approximately by the cascaded regulation with $\dot{X}(T) = 0$:

$$X = \frac{\eta}{1 + \left( \frac{\eta}{1+X^n} \right)^n} \tag{S13}$$

with $X$ being the pseudo-steady state value. These pseudo-steady states are similar though not generally equal to the bistability steady states for $-n < 0$. In any case, the period is approximately $2\gamma$, because if $X(T-2\gamma)$ is near a maximum, then (because the regulation is monotonically decreasing) $X(T-\gamma)$ is near a minimum, and $X(T)$ is near a maximum again.

For a more general estimate of the period, when $\gamma$ is not necessarily large compared to 1, we will assume a minimum of the right-hand side of Eq S12 at some value $X(T_0 - \gamma)$ for some time $T_0$. Then the left-hand side will have a maximum at $X(T_0)$. Furthermore, since from the above discussion we expect the difference in time between consecutive extrema to be approximately $\gamma$, we expect that $X(T)$ will have a maximum near $T_0 - \gamma + \gamma + \delta = T_0 + \delta$, for some small $\delta$. We can then expand $X(T)$ around $T_0 + \delta$ using a Taylor series:

$$X(T) \approx X(T_0 + \delta) + \frac{1}{2}(T - T_0 - \delta)^2 \ddot{X}(T_0 + \delta) \tag{S14}$$

where the first derivative term is absent since $X(T)$ is a maximum at time $T_0 + \delta$, making $\dot{X}(T_0 + \delta) = 0$. Note that $\dot{X}$ and $\ddot{X}$ values in this equation are constants, since they are evaluated at $T_0 + \delta$.

Now, let

$$\Psi(T) = \dot{X}(T) + X(T) \tag{S15}$$

be the left-hand side of Eq S12. Substituting in the expanded $X(T)$ from Eq S14 yields

$$\Psi(T) = (T - T_0 - \delta)\ddot{X}(T_0 + \delta) + X(T_0 + \delta) + \frac{1}{2}(T - T_0 - \delta)^2 \ddot{X}(T_0 + \delta). \tag{S16}$$

Finally, since $\Psi$ is at a maximum at $T = T_0$ (see above), its derivative is zero:

$$\dot{\Psi}(T) = \ddot{X}(T_0 + \delta) + (T - T_0 - \delta)\ddot{X}(T_0 + \delta) = (T - T_0 - \delta + 1)\ddot{X}(T_0 + \delta) = 0. \quad \text{(S17)}$$

Since we assumed the system is in the oscillatory regime, $\ddot{X}(T_0 + \delta) \neq 0$, leaving

$$\delta = 1 \quad \text{(S18)}$$

at $T = T_0$. Thus the time difference between consecutive extrema of $X(T)$ is approximately $(T_0 + \delta) - (T_0 - \gamma) = \gamma + 1$, and the period is twice this:

$$\text{Oscillation period} \equiv \Delta T_{\text{o.p.}} \approx 2(\gamma + 1). \quad \text{(S19)}$$

This period is in non-dimensionalized time units, but if we re-introduce the time dimension by returning to $\Delta t_{\text{o.p.}} = \Delta T_{\text{o.p.}}/\beta$ (see above nondimensionalization section of this appendix), we see that dimensionful period is

$$\Delta t_{\text{o.p.}} = 2(\tau + 1/\beta). \quad \text{(S20)}$$

Thus the oscillation period is twice the delay time plus the "intrinsic delay" due to degradation, which in the nondimensionalized equations is normalized to 1.

**Appendix S3 Analysis of autoregulation with leakage.** In this discussion we derive the boundaries for bistability and oscillations when a leakage (basal expression) is present.

To derive the bifurcation boundaries for autoregulation with non-zero leakage, we allow $\epsilon > 0$. The derivation follows similar steps to the non-leaky autoregulation case. First, the fixed points are given by setting $\dot{X} = 0$:

$$(X^* - \epsilon)(1 + X^{*n}) = \eta \quad \text{(S21)}$$

and the linearized equation is the same as without leakage, since $\dot{\epsilon} = 0$. Note, however, that the coefficient $M(X^*)$ depends on the fixed point value, which in turn depends on the leakage. With this information in hand, the derivation of saddle-node and Hopf bifurcation curves can be carried out as in the non-leaky case, but Eq S21 cannot be solved explicitly for $X^*$, making it impossible to solve for the bifurcation boundaries entirely analytically.

For the saddle-node bifurcation ($\lambda = 0$), $\eta$ and $\epsilon$ can be solved directly from Eq S21 and Eq S5 in terms of $X^*$, which serves as a parameterizing variable:

$$\begin{aligned} \epsilon &= \frac{X}{n}\left(n + 1 + X^{-n}\right) \\ \eta &= -\frac{X}{n}\left(1 + X^{-n}\right)\left(1 + X^{n}\right) \end{aligned} \quad \text{(S22)}$$

with the asterisks left off for clarity. For $\eta > 0$, Eq S22 requires $n < 0$, and therefore for $\epsilon > 0$, we must have at least $-n > 1$. The value of $\epsilon$ has a maximum and $\eta$ has a minimum for the same value of $X^*$, which can be calculated by setting the derivatives to zero. These extrema occur at

$$X^* = \left(\frac{n - 1}{n + 1}\right)^{1/n} \quad \text{(S23)}$$

with corresponding values

$$\begin{aligned} \epsilon &= \left(\frac{n - 1}{n + 1}\right)^{\frac{1-n}{n}} \\ \eta &= \frac{-4n}{(1 + n)^2}\left(\frac{n - 1}{n + 1}\right)^{\frac{1-n}{n}} \end{aligned} \quad \text{(S24)}$$

and a ratio

$$\frac{\epsilon}{\eta} = -\frac{(1+n)^2}{4n} \qquad . \tag{S25}$$

That these extrema correspond to a maximum of $\epsilon$ and a minimum of $\eta$ can be found by explicitly calculating the second derivatives of Eq S22 at the given value of $X^*$. Altogether, these results imply that for large enough leakage, the bistability is lost, while for smaller leakage, the minimum $\eta$ required for bistability actually decreases with increasing leakage. Furthermore, for $-n > 3 + 2\sqrt{2} \approx 6$, the cooperativity is high enough that bistability is possible even when the leakage is larger than the regulation strength ($\epsilon/\eta > 1$). This is because the $\eta$ requirement becomes small ($\eta \to 0$ as $n \to \infty$), while the $\epsilon$ requirement plateaus ($\epsilon \to 1$ as $n \to \infty$). The limit on $\epsilon$ also implies that leakage $\epsilon > 1$ never permits bistability, regardless of the value of $n$ or $\eta$. Fig. S3A shows this boundary for $n = -2$.

For the Hopf bifurcation ($\lambda = i\omega$), a similar procedure is followed. However, both $\omega$ and $X^*$ are now used as parameterizing variables. The boundary for $\gamma$ is the same as without leakage, because the $M$ values cancel in its derivation (see Appendix S2). For the conditions on $\eta$ and $\epsilon$, we follow the same procedure as for the no-leakage case. We plug in the value for $\gamma$ into the first of Eqs S8 and substitute the definition of $M$ (Eq 14) and $\eta$ (with leakage, Eq S21) to yield:

$$(-1)^{k+1} n \cdot \frac{1}{\sqrt{1+\omega^2}} \cdot \frac{X^{*n}}{1+X^{*n}} \cdot \frac{X-\epsilon}{X} = 1 \qquad . \tag{S26}$$

To balance this equation, we must have $(-1)^{k+1}n = |n|$, and since each fraction term is positive and less than 1 (the last term must be less than 1 to have positive $\eta$ in Eq S21), we must also have $|n| > 1$. Solving this equation for $\epsilon$ gives the $\epsilon$ boundary, and plugging back into the value of $\eta$ (Eq S21) yields the boundary for $\eta$. Altogether, we have an oscillation boundary defined by:

$$\begin{aligned}
\gamma &= \frac{1}{\omega}\left(-\tan^{-1}\omega + \pi k\right), \qquad k = 0, 1, \ldots \\
\epsilon &= X - X(1+X^{-n})\frac{\sqrt{1+\omega^2}}{|n|} \\
\eta &= \frac{\sqrt{1+\omega^2}}{|n|} X \left(1+X^{-n}\right)(1+X^n)
\end{aligned} \tag{S27}$$

For this boundary, there is no bound on $\epsilon$, but there is a bound on $\epsilon/\eta$. In particular, setting to zero the derivative of the $\eta$ bound with respect to $X$ and plugging into $\epsilon$ shows that there is a minimum $\eta$ for positive $\epsilon$ as long as

$$n > 1 + 2\sqrt{1+\omega^2} > 3 \qquad . \tag{S28}$$

There is also a bound on the ratio $\epsilon/\eta$, found by taking the derivative of the ratio with respect to $X$ and setting to zero. This maximum fractional leakage occurs at

$$\frac{1}{2}\left(\frac{n^2+1+\omega^2}{2n\sqrt{1+\omega^2}} - 1\right) \tag{S29}$$

where the individual values are

$$\begin{aligned}
\eta &= \frac{4n\sqrt{1+\omega^2}}{n^2-(1+\omega^2)}\left(\frac{n^2+1+\omega^2+2n\sqrt{1+\omega^2}}{n^2-(1+\omega^2)}\right)^{\frac{1}{n}} \\
\epsilon &= \left(\frac{n^2+1+\omega^2+2n\sqrt{1+\omega^2}}{n^2-(1+\omega^2)}\right)^{\frac{1-n}{n}}
\end{aligned} \tag{S30}$$

and with the slightly less restrictive requirement on the cooperativity

$$n > \sqrt{1+\omega^2} > 1 \qquad . \tag{S31}$$

Again, for large $n$ the minimum $\eta$ goes to zero, while the maximum $\epsilon$ goes to 1.

The bifurcation boundary for $n = 2$ are plotted in Fig. S3B.

**Appendix S4 Analysis of damped oscillations in autoregulation.** Here we sketch a derivation of why damped oscillations are expected in auto-repressors but not auto-activators.

The Hopf bifurcation for auto-repressors implies by definition oscillatory behavior on one side of the boundary and damped oscillations on the other side, for lower delay $\gamma$ and regulatory strength $\eta$. At $\gamma = 0$ there should be no damped oscillations because the Poincareé-Bendixson theorem for ODEs prevents the 1-dimensional system from crossing its own path, as required by any oscillations, damped or undamped. Similarly, $\eta = 0$ which lacks any feedback regulation has a trivial exponential solution without oscillations. The same is true for $n = 0$. Intuitively, we should expect damped oscillations somewhere between these lines ($\gamma = 0$, $\eta = 0$, $n = 0$) and the Hopf boundary, and monotonic (overdamped) behavior for $n < 0$.

Mathematically, for any $\gamma$ and $\eta$, we can take Eq 13 and rewrite it in the following form:

$$\gamma(\lambda+1)e^{\gamma(\lambda+1)} = -\eta M \gamma e^{\gamma}. \tag{S32}$$

Formally, the solution for $\lambda$ can be given in terms of Lambert's $W$ function [123], where $W(z)$ is the solution $z$ to the transcendental equation $f(z) = ze^z$ (i.e., the inverse of $f(z)$). That is,

$$\lambda = -1 + \frac{1}{\gamma} W(-\eta M \gamma e^{\gamma}). \tag{S33}$$

Lambert's $W$ function has an infinite number of branches, and thus an infinite number of solutions. It also has the properties that for $z \geq 0$, there is one real solution ($\operatorname{Im} z = 0$), and it is the solution with the largest real part of any solution. For $-1/e < z < 1$, there are two real solutions, again with the largest real parts of any solutions. For $z < -1/e$, there are no real solutions.

Setting $z = -\eta M \gamma e^{\gamma}$ from Eq S32, we can observe two facts. First, since the sign of $M$ is given by the sign of $n$, the largest eigenvalue $\lambda$ is always real for $n < 0$. For $n > 0$, we can find the boundary in which the largest eigenvalue is real by setting $-\eta M \gamma e^{\gamma} = -1/e$, noting as in the main text that $1/\eta M = (1 + X^{*-n})/n$. The boundary is then as follows:

$$\eta = n\gamma e^{\gamma+1}\left(n\gamma e^{\gamma+1} - 1\right)^{-\frac{n+1}{n}}. \tag{S34}$$

This curve has a vertical asymptote at $\gamma = W(1/ne)$ that approaches $\gamma = 0$ for $n \to \infty$ and infinity for $n \to 0$, and a horizontal asymptote that approaches $\eta = 0$ as $\gamma \to \infty$ for all $n$. The curve is not defined to the left of the vertical asymptote, because then arriving at Eq S34 would assume a negative steady state ($X^* < 0$).

Thus, for $n < 0$ and for values of $\eta$ and $\gamma$ above the boundary given by Eq S34, the largest (least negative) eigenvalue is real. This means that the slowest-decaying mode is non-oscillatory. Thus, there may be short transients depending on initial conditions that overshoot (for $\gamma > 0$) due to other non-real eigenvalues, but the approach to steady state after the initial transients, does not oscillate. That is, for autoactivation ($n < 0$) and for autorepression ($n > 0$) below the boundary in Eq S34, decay to steady state is overdamped; for autorepression above the boundary in Eq S34 (and below the Hopf boundary), decay to steady state is characterized by damped oscillations.

**Appendix S5 Full set of derivations related to the logic motif.** Here we include a detailed derivation for the logic-section equations.

#### Nondimensionalization

The dimensionful equations governing the logic motif are:

$$
\begin{aligned}
\dot{z}(t) &= \frac{\alpha_{z1}k_x^{n_1}}{k_x^{n_1} + x^{n_1}(t-\tau_1)} + \frac{\alpha_{z2}k_y^{n_2}}{k_y^{n_2} + y^{n_2}(t-\tau_2)} - \beta z(t) \\
\dot{r}(t) &= \frac{\alpha_r k_z^{n_3}}{k_z^{n_3} + z^{n_3}(t-\tau_z)} - \beta r(t).
\end{aligned} \tag{S35}
$$

Using the substitutions

$$
\begin{aligned}
&\eta_{Z1,2} = \frac{\alpha_{z1,2}}{k_z\beta}, \quad \gamma_{1,2,Z} = \tau_{1,2,Z}\beta, \quad \eta_R = \frac{\alpha_r}{k_r\beta}, \\
&X = \frac{x}{k_x}, \quad Y = \frac{y}{k_y}, \quad Z = \frac{z}{k_z}, \quad R = \frac{r}{k_r}, \quad T = t\beta
\end{aligned} \tag{S36}
$$

yields Eq 19 from the main text. The notation as before is with $n_i > 0$ for repressors and $n_i < 0$ for activators, ignoring leakage. The parameter $k_r$ is arbitrary, and could refer to, for example, the half-maximal input of $r$ on some downstream gene if $r$ is a transcription factor.

#### Alternative logic formulations

In the main text, we use a sum of Hill terms to describe logical behavior. Previous work has shown that other formulations can describe a range of logic gates as well [75, 76]. The fixed points given by various formulations will of course differ from one another. However, the analysis in this paper depends almost exclusively on linearized equations. Thus, for example, the logic based on sum of Hill terms (Eq 19) has a linearization around a fixed point $(X^*, Y^*, Z^*)$

$$
\dot{\delta Z}(T) + \eta_{Z1}M(X^*; n_1)\delta X(T-\gamma_1) + \eta_{Z2}M(Y^*; n_2)\delta Y(T-\gamma_2) + \delta Z(T) = 0. \tag{S37}
$$

This regulation terms are found by taking the derivatives with respect to $X(T-\gamma_1)$ and $Y(T-\gamma_2)$. The same is true for a multiplicative logic function, or in fact any function $g(X(T-\gamma_1), Y(T-\gamma_2))$ (see, e.g., Eqs 6 and 7 of Bhalekar 2019 [124]). For the multiplication of two Hill terms, we get:

$$
\begin{aligned}
\dot{\delta Z}(T) &+ \frac{\eta_{Z2}}{1 + {Y^*}^{n_2}}\eta_{Z1}M(X^*; n_1)\delta X(T-\gamma_1) \\
&+ \frac{\eta_{Z1}}{1 + {X^*}^{n_1}}\eta_{Z2}M(Y^*; n_2)\delta Y(T-\gamma_2) + \delta Z(T) = 0.
\end{aligned} \tag{S38}
$$

Thus, even though the terms are multiplied in the nonlinear equation, the linearized form produces separate terms, one for each delay. The multiplicative form only introduces an extra added constant in front of each term. The same argument can be carried over to feedforward loops (setting $Y \to X$), double feedback (setting $Y \to X$, $Z \to X$), and complex networks. Where we specify input functions for feedforward loops, the same input functions can easily be applied to a multiplicative logic function as well. For example, Eq S41 hold true for a product-of-Hill function with the + replaced by $\times$, resulting in very similar conclusions.

**Appendix S6 Full set of derivations related to the feedforward motif.** Here we include a detailed derivation for the feedforward-section equations.

#### Nondimensionalization

The dimensionful equation for the feedforward motif is:

$$\dot{z}(t) = \frac{\alpha_1 k_1^{n_1}}{k_1^{n_1} + x^{n_1}(t-\tau_1)} + \frac{\alpha_2 k_2^{n_2}}{k_2^{n_2} + x^{n_2}(t-\tau_2)} - \beta z(t) \tag{S39}$$

Normalizing yields Eq 20 from the main text, with

$$\eta_{1,2} = \frac{\alpha_{1,2}}{k_1 \beta}, \quad \gamma_{1,2} = \tau_{1,2}\beta, \quad K = \frac{k_1}{k_2}, \quad X = \frac{x}{k_1}, \quad Z = \frac{z}{k_z}, \quad T = t\beta \tag{S40}$$

#### Pulse response

For the square input pulse (Eq 22), $X$ only takes on two values, so the Hill terms can be evaluated explicitly as follows:

$$\begin{aligned}
&\dot{Z}(T) = \\
&\quad \eta_1 \left[ \frac{1}{1+X_0^{n_1}} + \left( \frac{1}{1+\eta_X^{n_1}} - \frac{1}{1+X_0^{n_1}} \right) (\Theta(T) - \Theta(T-\omega)) \right] \\
&+ \eta_2 \left[ \frac{1}{1+K^{n_2} X_0^{n_2}} + \left( \frac{1}{1+K^{n_2}\eta_X^{n_2}} - \frac{1}{1+K^{n_2} X_0^{n_2}} \right) (\Theta(T-\Delta\gamma) - \Theta(T-\Delta\gamma-\omega)) \right] \\
&- Z(T) \qquad .
\end{aligned} \tag{S41}$$

Eq 21 is linear in all time-dependent terms, making the solution straightforward using Laplace transforms. Denoting the Laplace transform of $Z(T)$ as $\tilde{Z}(s)$, we have that the Laplace transform of $\dot{Z}(T)$ is $\tilde{Z}(s) - Z(0)$, and the Laplace transform of $a\Theta(T-\gamma)$ is $(a/s)e^{-\gamma s}$. Solving for $\tilde{Z}(s)$ and then taking the reverse Laplace transform yields Eqs 23 and 24.

#### Frequency response

The sinusoidal input (Eq 25) oscillates between zero and twice the amplitude $2A$ at a frequency $f$. Note that rearranging Eq 21 as

$$Z(T) + \dot{Z}(T) = \frac{\eta_1}{1+X^{n_1}(T)} + \frac{\eta_2}{1+K^{n_2} X^{n_2}(T-\Delta\gamma)}, \tag{S42}$$

we see that the input and output variables can be placed on opposite sides of the equation. Since $X(T)$ is periodic with frequency $f$, and the right-hand side has no other time-dependent terms, the whole right-hand side must be periodic with a fundamental frequency of $f$ as well. The same applies to both terms on the right-hand side independently. We can decompose the first term in a Fourier series as in Eq 26 where the Fourier coefficients $a_k^{x;n}$ (with $n = n_1$) are determined by Eq 27 for all $f > 0$. Since the input is chosen to be a pure cosine, there are no sine terms in the Fourier expansion.

The second term on the right of Eq S42 is shifted in time and uses a different Hill coefficient. The shift in time introduces a phase shift, but no change in the amplitudes. Thus, for the second term we have

$$\begin{aligned}
&\frac{\eta_2}{1+X^{n_2}(T)} = \\
&\quad \eta_2 \left( \frac{a_0^{x;n_2}}{2} + \sum_{k=1}^{\infty} a_k^{x;n_2} \left[ \cos(2\pi f k \Delta\gamma)\cos(2\pi k f T) + \sin(2\pi f k \Delta\gamma)\sin(2\pi k f T) \right] \right)
\end{aligned} \tag{S43}$$

using the same Eq 27 to determine the coefficients (with $n = n_2$). If $K \neq 1$, then we must also replace $A \to AK$ in the integral. We will not have need to discuss $K$ further, but one may keep in mind that all values of $a_k^{x;n_2}$ below reflect this modification in $A$.

Now, since the right-hand side of Eq S42 (the input) is periodic with fundamental frequency $f$, the left hand side of the equation (the output) must have the same fundamental frequency in order for both sides of the equation to be simultaneously periodic with period $1/f$. The same must hold true for both the output $Z(T)$ and its derivative $\dot{Z}(T)$. Thus we can decompose $Z(T)$ in a Fourier series as well and the calculate the derivative of this decomposition:

$$\begin{aligned} Z(T) &= \frac{a_0^z}{2} + \sum_{k=1}^{\infty} a_k^z \cos(2\pi f k T) + b_k^z \sin(2\pi f k T) \\ \dot{Z}(T) &= \sum_{k=1}^{\infty} 2\pi f k b_k^z \cos(2\pi f k T) - 2\pi f k a_k^z \sin(2\pi f k T) \end{aligned} . \tag{S44}$$

Summing the Fourier decompositions on either side of Eq S42, we can equate the Fourier coefficients on either side to determine the expressions for the output coefficients $a_k^z$ and $b_k^z$, yielding the assignments in Eq 29.

The Fourier decomposition can also be written in terms of magnitude and phase instead of cos and sin components. In this format, $Z(T) = \frac{a_0^z}{2} + \sum_{k=1}^{\infty} I_k \cos(2\pi f k T - \phi_k)$ (see Eq 30) with

$$\begin{aligned} I_k &= \sqrt{{a_k^z}^2 + {b_k^z}^2} \\ \phi_k &= \operatorname{atan2}\left(b_k^z, a_k^z\right) \end{aligned} , \tag{S45}$$

which upon plugging in the values from Eq 29 yields Eq 30.

Because a Hill function for $n < 0$ is equal to one minus the same Hill function for $|n|$, it is easy to see from Eq 27 that $a_k^{x;-n} = -a_k^{x;n}$. Thus, at frequencies $f$ that are integer multiples of $1/k\Delta\gamma$, the output magnitude $I_k$ increases to a local maximum for coherent feedforward motifs ($\mathrm{sgn}(n_1) = \mathrm{sgn}(n_2)$) and decreases to a local minimum for incoherent feedforward motifs ($\mathrm{sgn}(n_1) = \mathrm{sgn}(n_2)$). The opposite holds for frequencies that are half-integer multiples of $1/k\Delta\gamma$. For the special case of perfectly balanced incoherent feedforward motifs ($\eta_1 = \eta_2$, $n_1 = -n_2$), the magnitudes $I_k$ decrease to zero for frequencies that are half-integer multiple of $1/\Delta\gamma$; otherwise, the maxima (for coherent) and minima (for incoherent) equal $I_k(f = 0)$.

**Appendix S7 Full set of derivations related to the multi-component feedback motif.** Here we include a detailed derivation for the feedback-section equations.

#### Nondimensionalization

The dimensionful equations for two-component feedback (assuming equal degradation rates for simplicity) are given by:

$$\begin{aligned} \dot{x}(t) &= \frac{\alpha_1 k_1^{n_1}}{k_1^{n_1} + y^{n_1}(t-\tau_1)} - \beta x(t) \\ \dot{y}(t) &= \frac{\alpha_2 k_2^{n_2}}{k_2^{n_2} + x^{n_2}(t-\tau_2)} - \beta y(t) \end{aligned} . \tag{S46}$$

Setting $X = x/k_2$, $Y = y/k_1$, $T = \beta t$, $\eta_1 = \alpha_1/k_1\beta$, $\eta_2 = \alpha_2/k_2\beta$, $\gamma_1 = \tau_1\beta$, and $\gamma_2 = \tau_2\beta$ results in the nondimensionalzed equations found in the main text (Eqs 33).

#### Fixed points

The fixed points correspond to

$$\begin{aligned}\eta_1 &= X(1+Y^{n_1})\\ \eta_2 &= Y(1+X^{n_2})\end{aligned} \tag{S47}$$

or, similar to the stability of the origin for direct autoactivation,

$$X = Y = 0 \tag{S48}$$

for two activators.

#### Linearization

Linearizing around these fixed points and assuming solutions of the form $\delta X(T) = A\exp(\lambda_1 T), \delta Y(T) = B\exp(\lambda_2 T)$, we find that:

$$\begin{aligned}\lambda_1 + \eta_1 M_1 \frac{B}{A} e^{(\lambda_2-\lambda_1)T} e^{-\lambda_2\gamma_1} + 1 = 0\\ \lambda_2 + \eta_2 M_2 \frac{A}{B} e^{(\lambda_1-\lambda_2)T} e^{-\lambda_1\gamma_2} + 1 = 0.\end{aligned} \tag{S49}$$

In principle, the two $\lambda$'s could bifurcate separately. However, Eqs 34 can only reflect an actual solution of the linearized system if $\lambda_1 = \lambda_2 \equiv \lambda$; otherwise there is an exponential term $\exp((\lambda_1 - \lambda_2)T)$ will not be true for all time. Thus the characteristic equations are

$$\begin{aligned}\lambda + \eta_1 M_1 \frac{B}{A} e^{-\lambda\gamma_1} + 1 = 0\\ \lambda + \eta_2 M_2 \frac{A}{B} e^{-\lambda\gamma_2} + 1 = 0.\end{aligned} \tag{S50}$$

Since $B$ and $A$ can in general be complex, we will write the ratio $B/A$ in the form $B/A = Re^{i\phi}$, where $R$ is the magnitude of the ratio and $\phi$ is the phase. We assume neither $A$ nor $B$ equal zero, because that would imply a lack of regulation.

Subtracting the two equations in Eq S50 reduces to

$$R^2 e^{2i\phi} = \frac{\eta_2 M_2}{\eta_1 M_1} e^{\lambda(\gamma_1-\gamma_2)} \tag{S51}$$

If we let $\lambda = \mu + i\omega$ to designate separately the real ($\mu$) and imaginary ($\omega$) parts of $\lambda$, we can solve for $R$ and $\phi$ to yield:

$$\begin{aligned}R &= e^{\mu(\gamma_1-\gamma_2)/2}\sqrt{\frac{\eta_2|M_2|}{\eta_1|M_1|}}\\ \phi &= \frac{\omega}{2}(\gamma_1-\gamma_2) + \frac{\pi}{4}(\operatorname{sgn} n_1 - \operatorname{sgn} n_2) + \pi k\end{aligned} \tag{S52}$$

for integer $k$. The second term of $\phi$ is simply a way of writing that there is an extra $\pm\pi/2$ if the signs of $n_1$ and $n_2$ are not the same. The $\pi k$ accounts for a possible negative sign on taking the square root of $R^2$. We leave off the $k$ in the main text (Eq 39), as we account for it elsewhere (see below). One can also freely add integer multiples of $2\pi$ to $\phi$ with no effect, since it is a phase. Note also that the phase does not depend on $\mu$, and so holds anywhere in the parameter space, not only at a bifurcation where $\mu = 0$.

We can also write Eqs S50 in matrix form. First, we multiply the top equation by $A$ and the bottom by $B$ (Eq S50 from the main text). Then, we can write the equation as:

$$J\vec{a} = \begin{pmatrix} 1+\lambda & \eta_1 M_1 e^{-\gamma_1\lambda} \\ \eta_2 M_2 e^{-\gamma_2\lambda} & 1+\lambda \end{pmatrix}\begin{pmatrix} A \\ B \end{pmatrix} = 0 \tag{S53}$$

where $J$ refers to the matrix and $\vec{a}$ to the vector of displacement coefficients ($\delta X, \delta Y$ at $T=0$). $J$ can then be diagonalized, $J = S\Lambda S^{-1}$ for diagonal $\Lambda$. The diagonal elements of $\Lambda$ are the eigenvalues of $J$,

$$\Lambda_{\pm} = \lambda \pm \sqrt{\eta_1 M_1}\sqrt{\eta_2 M_2} e^{-\lambda(\gamma_1+\gamma_2)/2} + 1 \tag{S54}$$

Note that the square roots cannot be combined because $\sqrt{-1}\sqrt{-1} = i^2 = -1$ but $\sqrt{(-1)(-1)} = \sqrt{1} = +1$. The columns of $S$ are the eigenvectors of $J$. After pulling out a constant factor of $\sqrt{\eta_1 M_1}\exp(\gamma_2 - \gamma_1)\lambda/4$ to clean up the form (since an eigenvector times a constant is still an equivalent eigenvector), we have eigenvectors

$$\vec{v}_{\pm} = \begin{pmatrix} \pm 1 \\ \frac{\sqrt{\eta_2 M_2}}{\sqrt{\eta_1 M_1}} e^{(\gamma_1-\gamma_2)\lambda/2} \end{pmatrix} \tag{S55}$$

Substituting in $\lambda$ from Eq S51 yields

$$\vec{v}_{\pm} = \begin{pmatrix} \pm 1 \\ Re^{i\phi} \end{pmatrix} \tag{S56}$$

As noted above, we do not need to account for the possible $\pi$ phase in $\phi$ (if $k$ is odd), because the possible difference of $\pi$ in phase is exactly accounted for by the two possible eigenvectors. We thus set $k=0$ in the use of $\phi$ here and do not include $k$ in further discussion.

Due to this diagonalization, we can rewrite Eq S53 as

$$\Lambda S^{-1}\vec{a} = \Lambda \vec{a}' = 0 \tag{S57}$$

with the eigenvector-basis components $\vec{v} \equiv S^{-1}\vec{a}$. For this to hold, we have either $\Lambda_i = 0$ or $\vec{a}'_i = 0$. Now, $S^{-1}\vec{a}$ is the vector of coordinates of $\vec{a}$ in the eigenvector basis. This means that if the first element of $\vec{a}'$ is zero, then $\vec{a}$ is proportional to the second eigenvector, and therefore Eq S57 is satisfied if the second eigenvalue of $\Lambda$ also equals zero. Likewise, if the second element is zero, then $\vec{a}$ is proportional to the first eigenvector and the first eigenvalue of $\Lambda$ must be zero for Eq S57 to hold for both components.

Thus, instead of considering when $S^{-1}\vec{a} = 0$, we can focus entirely on the eigenvalues and eigenvectors of $J$. Explicitly, we can write $\vec{a}$ in the basis of the eigenvectors $\vec{v}_{\pm}$ with components parallel to the two eigenvectors $\nu_+$ and $\nu_-$. Thus, from Eq S57:

$$\Lambda S^{-1}\vec{a} = \Lambda S^{-1}(\nu_+\vec{v}_+ + \nu_-\vec{v}_-) = \nu_+\Lambda S^{-1}\vec{v}_+ + \nu_-\Lambda S^{-1}\vec{v}_- = \nu_+\Lambda \begin{bmatrix} 1 \\ 0 \end{bmatrix} + \nu_-\Lambda \begin{bmatrix} 0 \\ 1 \end{bmatrix} \tag{S58}$$

which in turn implies that we have two independent equations to satisfy Eq S57:

$$\Lambda_{\pm} = 0 \tag{S59}$$

corresponding to independent behavior of the two eigenvectors, $\vec{v}_{\pm}$, respectively. In fact, $J$ (for $\gamma_1 = \gamma_2$) is often considered a matrix characteristic equation on its own and $\vec{a}$ is ignored entirely [58, 125, 126].

##### Bifurcations

Like the autoregulation case, we expect the system to undergo bifurcations at $\lambda = 0$ (saddle-node bifurcation) and at $\lambda = i\omega$ (Hopf bifurcation). Because we have two dimensions, however, we can ask what happens to each eigenmode for each type of bifurcation. The eigenvectors, or eigenmodes $v_{\pm}$, represent relative changes in $\delta X$ and

$\delta Y$ that grow or shrink independent of the changes in the other mode. We expect each mode $\vec{v}_i$ to grow when $\Lambda_i > 0$ and shrink when $\Lambda_i < 0$. The bifurcation for a given mode $\vec{v}_i$ thus happens at $\Lambda_i = 0$, in keeping with the matrix characteristic equation (Eq S57). We thus look at $\lambda = 0$ and $\lambda = i\omega$ for $\Lambda_+ = 0$ and for $\Lambda_- = 0$. Because purely real values of $\lambda$ solving $\Lambda_\pm = 0$ have greatest real part (see Appendix S4), if a mode bifurcates via a saddle-node bifurcation, we will not observe meaningful Hopf bifurcations in that mode. However, the other mode may still show a Hopf bifurcation. As described below and in the main text, this is exactly what leads to transient oscillations.

For the saddle-node bifurcations, we set $\lambda = 0$. Then Eq S59 reduces to

$$\eta_1 \eta_2 M_1 M_2 = +1 \tag{S60}$$

which corresponds to $\Lambda_-$ for $n_1 > 0, n_2 > 0$ or $\Lambda_+$ for $n_1 < 0, n_2 < 0$. No solutions are possible if $\operatorname{sgn} n_1 \neq \operatorname{sgn} n_2$. Plugging in the definitions of $M$ (Eq 14) and $\eta_1, \eta_2$ (Eq S47), we arrive at the equation $f(X, Y) = 1$ of Eq 36 in the main text. Solving for $X$ in terms of $Y$ and vice versa, and plugging into Eq S47 yields the $\eta_1, \eta_2$ boundaries found in Eq 36, completing the definition of the saddle-node boundaries. The delays do not appear in the boundary, and are thus not a factor in this bifurcation.

For the Hopf bifurcations, we set $\lambda = i\omega$. To stay agnostic to the signs of $n_1$ and $n_2$, we set $M_j = \operatorname{sgn} n_j |M_j|$ and note that $\sqrt{n_j} = \exp(\frac{i}{2} \operatorname{Arg} n_j)$, where $\operatorname{Arg} x$ yields 0 for $x \geq 0$ and $\pi$ for $x < 0$. This allows us to then separate the real and imaginary parts of Eq S59, yielding

$$\begin{aligned} \pm\sqrt{\eta_1 \eta_2 |M_1||M_2|} \cos\left(-\omega \frac{\gamma_1 + \gamma_2}{2} + \frac{\operatorname{Arg} n_1 + \operatorname{Arg} n_2}{2}\right) &= -1 \\ \pm\sqrt{\eta_1 \eta_2 |M_1||M_2|} \sin\left(-\omega \frac{\gamma_1 + \gamma_2}{2} + \frac{\operatorname{Arg} n_1 + \operatorname{Arg} n_2}{2}\right) &= -\omega \end{aligned} \quad . \tag{S61}$$

Solving one of the equations for the square-root term and plugging into the other, then solving for $\gamma_1 + \gamma_2$ yields:

$$\gamma_1 + \gamma_2 = \frac{2}{\omega}\left(-\tan^{-1}\omega + \frac{\operatorname{Arg} n_1 + \operatorname{Arg} n_2}{2} + k'\pi\right) \tag{S62}$$

for integer $k'$. We can also rearrange Eq S52 to solve for $\gamma_1 - \gamma_2$:

$$\gamma_1 - \gamma_2 = \frac{2}{\omega}\left(\phi - \frac{\pi}{4}\left(\operatorname{sgn} n_1 - \operatorname{sgn} n_2\right)\right) \quad . \tag{S63}$$

Adding these two equations and dividing by two yields an equation for $\gamma_1$. Subtracting the two and dividing by two yields an equation for $\gamma_1$. Replacing all the $\operatorname{Arg} x$ terms with the equivalent $(1 - \operatorname{sgn} x)\pi/2$ yields the equations given in the main text (Eq 37).

Next, we can substitute Eq S62 into Eq S61 and simplify using the relations $\cos(\tan^{-1} x) = 1/\sqrt{1 + x^2}$ and $\sin(\tan^{-1} x) = x/\sqrt{1 + x^2}$. Then, substituting in the definitions of $M$ (Eq 14) and $\eta_1, \eta_2$ (Eq S47), we get:

$$\sqrt{\frac{|n_1 n_2|}{1 + \omega^2} \cdot \frac{X^{n_2} Y^{n_1}}{(1 + X^{n_2})(1 + Y^{n_1})}} = (-1)^{k''} \tag{S64}$$

where $k'' = k'$ for $\Lambda_- = 0$ and $k'' = k' + 1$ for $\Lambda_+ = 0$. Squaring this equation yields $f(X, Y) = 1$ of Eq 37 in the main text. However, because un-squared the left side is still positive, it also requires that $k'$ in Eq S62 be even for $\Lambda_- = 0$ and odd for $\Lambda_+ = 0$. In particular, the minimum bifurcation boundaries will then be for $k' = 0$ and $k' = 1$, respectively.

Taking $f = 1$ (or Eq S64), one can solve for $X$ in terms of $Y$ and vice versa, and then plug into Eq S47 to yield the $\eta_1, \eta_2$ boundaries found in Eq 37. This completes the definition of the Hopf boundaries.

Because $\Lambda_-$ (for $n_1 > 0, n_2 > 0$) and $\Lambda_+$ (for $n_1 < 0, n_2 < 0$) had already bifurcated via saddle-node, the noticeable bifurcations will be for the other mode in each case ($\Lambda_+$ for $n_1 > 0, n_2 > 0$ and $\Lambda_-$ for $n_1 < 0, n_2 < 0$). In fact, one can check that the boundaries in Eq 37 lie entirely above those in Eq 36. Thus, due to the definitions of $\vec{v}_\pm$ (Eq S56), $X$ and $Y$ will oscillate synchronously ($\vec{v}_+$, $\phi = 0$) for $n_1 > 0, n_2 > 0$ and anti-synchronously ($\vec{v}_-$, $\phi = \pi$) for $n_1 < 0, n_2 < 0$). For the cases in which $\operatorname{sgn} n_1 \neq \operatorname{sgn} n_2$, both $\Lambda_+$ and $\Lambda_-$ may show results; however, because the minimum bifurcations occur for $k' = 0$ for $\Lambda_-$ and $k' = 1$ for $\Lambda_+$, the $\Lambda_+$ boundary lies above the $\Lambda_-$ boundary. By the definition of $\vec{v}_-$ (Eq S56), these oscillations will have a phase difference of $\phi = \pi/2$ ($Y$ lags $X$) for $n_1 < 0, n_2 > 0$ and $\phi = -\pi/2$ ($Y$ leads $X$) for $n_1 > 0, n_2 < 0$.

**Appendix S8 Full set of derivations related to double feedback networks.** Here we include a detailed derivation related to the double feedback motif. (Appendix S7).

##### Nondimensionalization

The dimensionful equation governing the double feedback motif are:

$$\dot{x}(t) = \frac{\alpha_1 k^{n_1}}{k^{n_1} + x^{n_1}(t - \tau_1)} + \frac{\alpha_2 k^{n_2}}{k^{n_2} + x^{n_2}(t - \tau_2)} - \beta x(t) \tag{S65}$$

in which we assume the values of $k$ in the two terms are identical for simplicity. Using the substitutions

$$\eta_{1,2} = \frac{\alpha_{1,2}}{k\beta}, \quad \gamma_{1,2} = \tau_{1,2}\beta, \quad X = \frac{x}{k}, \quad T = t\beta \tag{S66}$$

yields Eq 41 from the main text.

##### Nonmonotonicity

The derivative of Eq 41 is $-\eta_1 M_1 - \eta_2 M_2 = -(n_1\eta_1|M_1| + n_2\eta_2|M_2|)$, where $M_1$ and $M_2$ are defined as usual, corresponding to the two regulation terms, respectively. All the variables on the right side of this equation are positive except the Hill coefficients. If they have the same sign ($\operatorname{sgn} n_1 = \operatorname{sgn} n_2$), then the sign of the derivative never changes, meaning the overall regulation is monotonic. If they are of opposite sign, but equal magnitude ($n_1 = -n_2 \equiv n$), then the derivative reduces to $n(\eta_1 - \eta_2)|M|$ (where $M_1 = M_2 \equiv M$). This derivative also has a constant sign, equal to the sign of $n(\eta_1 - \eta_2)$. If the Hill coefficients are of opposite sign and have unequal magnitudes, there will be some $X$ for which the derivative is either positive or negative, making the overall regulation non-monotonic.

**Appendix S9 Full set of derivations related to arbitrary-sized networks.** Here we include a detailed derivation related to arbitrary networks, as mentioned in the Discussion section. The arguments follow the derivation for two-component feedback (Appendix S7).

We can write a set of delay differential equations corresponding to $N$ components using sum-of-Hills regulation terms as follows:

$$\dot{X}_i(T) = \sum_{j\,\forall\, j\,:\, X_j \text{regulates} X_i}^{N} \frac{\eta_{ij}}{1 + X_j^{n_{ij}}(T - \gamma_{ij})} - X_i(T) \tag{S67}$$

where the sum is over all values of $j$ for which $X_j$ regulates $X_i$. This includes cases of autoregulation in which $j = i$. We assume all half-maximal input parameters can be normalized to 1 for simplicity, which is true as long as all regulation terms of a given gene have the same half-maximum parameter). Otherwise, there will be an additional parameter $K_{ij}^{n_{ij}}$ in multiplying $X_j$ in the denominator. We also ignore leakage, which can be added back as a single $\epsilon_i$ outside the sum. The fixed points are given by

$$X_i^* = \sum_{j \,\forall\, j\,:\, X_j \text{ regulates } X_i}^{N} \frac{\eta_{ij}}{1 + X_j^{*\,n_{ij}}} \tag{S68}$$

and $X_i^* = 0$ for all $X_i$ that are only regulated by activators.

Linearizing around the fixed points yields the linear governing equations

$$\delta \dot{X}(T) + \eta_{ij} M(X_j^*) \delta X_j(T - \gamma_{ij}) + \delta X_i(T) = 0 \tag{S69}$$

Plugging in assumed solutions of $\delta X_i(T) = A_i e^{\lambda_i T}$ yields a set of characteristic equations. If all the components are coupled, all the eigenvalues must be identical ($\lambda_i \equiv \lambda$) to avoid time dependence. If there are decoupled groups (i.e., completely separate networks), then each should be analyzed separately. Assuming they are all interconnected, the characteristic equations are then:

$$A_i(\lambda + 1) + A_j \sum_{j}^{N} \eta_{ij} M_j e^{-\lambda \gamma_{ij}} = 0. \tag{S70}$$

These equations can be rewritten in matrix form as

$$J \vec{a} = 0 \tag{S71}$$

in which we identify $a_i = A_i$ and

$$J_{ij} = (\lambda + 1)\delta_{ij} + \begin{cases} \eta_{ij} M_j e^{-\lambda \gamma_{ij}} & j \text{ regulates } i \\ 0 & \text{otherwise} \end{cases} \tag{S72}$$

which is graphically depicted in Fig. S5.

In this form, all the $X_i$ are coupled together. Diagonalizing $J = S \Lambda S^{-1}$ yields an equation $\Lambda \vec{a}' = 0$ in which $\vec{a}' = S^{-1}\vec{a}$ and the columns of $S$ are eigenmodes of the diagonal matrix $\Lambda$. Each row of this matrix equation can be true when either $\Lambda_i = 0$ or $\vec{a}' = 0$. However, we can write $\vec{a}$ in the basis of the eigenvectors $\vec{v}_i$ with components $\nu_i$ parallel to the eigenvectors. Explicitly,

$$\Lambda S^{-1} \vec{a} = \Lambda S^{-1} \left( \sum_{i}^{N} \nu_i \vec{v}_i \right) = \sum_{i}^{N} \nu_i \Lambda S^{-1} \vec{v}_i = \sum_{i}^{N} \nu_i \Lambda \vec{e}_i = \sum_{i}^{N} \nu_i \Lambda_i \vec{e}_i \tag{S73}$$

in which $\vec{e}_i$ are the standard basis vectors with the $j^{\text{th}}$ component equal to 1 if $j = i$ and zero otherwise. The penultimate expression follows because $S^{-1}$ by definition yields the components in the eigenbasis of the vector it acts on. The last expression, where $\Lambda_i$ denotes the $i^{\text{th}}$ diagonal element of $\Lambda$, then immediately follows because $\Lambda$ is diagonal. This implies that we have $N$ independent equations to satisfy Eq S71:

$$\Lambda_i = 0 \tag{S74}$$

corresponding to independent behavior of the eigenvectors $\vec{v}_i$, respectively. Thus, instead of considering when $S^{-1}\vec{a} = 0$, we can focus entirely on the eigenvalues and eigenvectors of $J$. Saddle-node bifurcations ($\lambda = 0$) and Hopf bifurcations ($\lambda = i\omega$) can be found for each mode independently. In fact, as noted in Appendix S7, $J$ (for a single delay $\gamma_{ij} \equiv \gamma$) is often considered a matrix characteristic equation on its own and $\vec{a}$ is ignored entirely [58, 125, 126].
